# PiggyBac Transposable Element-derived 1 controls Neuronal Progenitor Identity, Stress Sensing and mammal-specific paraspeckles

**DOI:** 10.1101/2021.05.19.444448

**Authors:** Tamás Raskó, Amit Pande, Kathrin Radscheit, Annika Zink, Manvendra Singh, Christian Sommer, Gerda Wachtl, Orsolya Kolacsek, Gizem Inak, Attila Szvetnik, Spyros Petrakis, Mario Bunse, Vikas Bansal, Matthias Selbach, Tamás Orbán, Alessandro Prigione, Laurence D. Hurst, Zsuzsanna Izsvák

## Abstract

The evolution and functional integration of new genes, especially those that become core to key functions, remains enigmatic. We consider the mammal-specific gene, *piggyBac* transposable element derived 1 (PGBD1), implicated in neuronal disorders. While it no longer recognises *piggyBac* transposon-like inverted repeats and transposase functionality having been lost, it has evolved a core role in neural homeostasis. Depletion of PGBD1 triggers accumulation of mammal-specific paraspeckles and neural differentiation. It acts by two modalities, DNA binding and protein-protein interaction. As a transcriptional repressor of (lnc)NEAT1, the backbone of paraspeckles, it inhibits paraspeckle formation in neural progenitor cells (NPCs). At the protein level it is associated with the stress response system, a function partially shared with (lnc)NEAT1. PGBD1 thus presents as an unusual exemplar of new gene creation, being a recently acquired multi-function, multi-modal gene. Mammalian specificity associated with control of a mammal-specific structure implies coevolution of new genes with new functions.

---

Where do new genes come from and how do they integrate into existing systems? The origin of novelty more generally is intriguing in its own right, but is especially so for new genes that evolve to become key to important cellular or organismic processes. How, one must question, did the organism cope prior to the evolution of what is now a key gene? The generation of new genes by duplication is a well understood route to the origin of novelty^1^, including key genes. Duplicate genes have already existing functions and, depending on the mode of duplication, they can be born with functional promoters too. “Tinkering”^2^ with such extant functionality is thus an accessible route to the evolution of new genes with new (neofunctionalization) or divided (subfunctionalization) functions. In the latter instance, partitioning two different key functions between two duplicates is an accessible route to the evolution and retention of new core genes. There is, however, increasing realization that this is by no means the only route to new gene creation, *de novo* origination or co-option of inactivated transposable elements being two alternative paths^3–5^.In these instances, the mechanisms behind the evolution of novel core functionality is less evident.

By definition, when inactivated transposable elements (TEs) are domesticated^6^they evolve into novel genes with a function beneficial for the host. TE derived sequences are ripe for co-option, as the transposable elements most successful at propagating through a genome after horizontal transfer, are those with effective promoter elements and functional products. Domesticated TEs have been incorporated in various host biological processes. The well-known examples include the adaptive immune system (*RAG1* in V(D)J recombination) in jawed vertebrates^7–9^ and in bacteria(*CRISPR/Cas*)^10, 11^, defence against other TEs (e.g. *Abp1*, *Schizosaccharomyces pombe*)^12^, immunity and far-red light phototransduction in plants *(Arabidopsis thaliana, FAR1/FHY3*)^13^, telomere integrity control (TERT) in eukaryotes^14, 15^, mammalian placentation (syncytin)^16, 17^, toti/pluripotency regulation in mice/primates (*MERVL/HERVH*, *L1TD1*)^18–21^, regulated elimination of DNA to decrease genome complexity of *Paramecium* genomes (*PiggyMac*)^22^.

Besides providing sequence for new proteins (as above), TE-derived sequences can also provide transcriptional signals (e.g. enhancers, polyA, etc)^19, 23, 24^ or structural elements, and affect gene expression at both DNA and RNA levels. For example, inverted repeated *Alu* elements (IRAlus) affect nuclear retention of mRNAs of host-encoded genes by generating secondary structure at 3’ UTRs of host genes^25–28^. Some IRAlus carrying mRNAs, such as Nicolin 1 (NICN1) and LIN28, are regulatory factors of pluripotency^27, 29–31^ and/or associated with stress response (e.g. long intergenic noncoding RNA (Linc-p21)^32^. These IRAlu mRNAs are specifically recruited to certain non-membrane-bound nuclear bodies notably paraspeckles. Paraspeckles are mammal-specific dynamic structures in the nucleus^33–36^, consisting of a special set of coiled-coil domain (<u>NO</u>NO/<u>P</u>ara<u>s</u>peckle, NOPS) containing RNA-binding proteins (e.g. SFPQ, NONO, PSPC1), assembled around the long noncoding RNA (lnc)NEAT1 (Nuclear Enriched Abundant Transcript 1 isoform2).

The domestication of TEs often involves co-option of pre-existing functionality. For example, TE proteins domesticated to suppress TEs or viruses in response to genetic conflicts had at the time of population invasion interactions with the cellular machinery and with the TEs themselves^37^. Another example, *PiggyMac*, sharing sequence similarity to the *piggyBac* superfamily of transposases, utilizes the catalytic activity of the transposase, and performs regulated genome-wide self-splicing^38, 39^. Similarly, mammalian syncytins employ the fusogenic capacity of the ancestral *env* gene in generating the placental syncytial cells^17, 40^. The evolutionary route to incorporation of such co-option of existing functions is relatively straightforward to comprehend. By contrast, co-option of TE proteins and sequences for new functions is more enigmatic. Examples include the adaptive immune systems of prokaryotes and vertebrates (e.g. CRISPR/Cas and V(D)J recombination, respectively). In such instances their interaction with the host cellular machinery needs to be better characterized to obtain a more complete picture of how the novel biological processes evolved. In these examples, TE-derived function is only rarely deciphered, not least because such characterization requires multiple approaches and manipulations^41–43^.

Here, motivated by this gap in knowledge, we sought to provide an in-depth analysis of mammalian *piggyBac* transposable element derived 1 (PGBD1). The domestication of *piggyBac* transposases has been observed in several evolutionary lineages: *Paramecium tetraurelia*^39^, *Tetrahymena thermophila*^44^, *Xenopus tropicalis*^45^. The human genome has five genes that are related to the *piggyBac* transposon (*PB*), hence the name, *piggyBac* transposable element derived 1-5, PGBD1-5^46^. The *PB* transposon, a widely used molecular tool for transgenesis^47^,was isolated from the genome of an insect, the cabbage looper moth (*Trichoplusiani*)^46^. The ancestral *PB*-like transposons were transferred horizontally to vertebrates in multiple waves^48^. The transposase-like open reading frames (ORFs) of extant PGBDs are subject to purifying selection^49^, with low nonsynonymous (*Ka*) to synonymous (*Ks*) nucleotide substitution ratio(*Ka*/*Ks* < 0.4)^49^,indicative of functionality^50^. Nevertheless, their current function (or functions) in humans is unknown.

PGBD1 is especially interesting for several reasons. First, PGBD1 (along with PGBD2) appears to be mammal-specific and hence evolutionarily highly taxonomically restricted^49^. PGBD5 by contrast is seen widely within the vertebrates^49^. Second, there is prior evidence to suppose that PGBD1 might not just be functional (as evidenced by Ka/Ks <1) but that it has evolved some core functionality. Notably, as with PGBD3 and 5^48, 51–53^, PGBD1 may be so well integrated that mutations within it are disease-associated. Genome-wide association studies (GWASs) identified single-nucleotide polymorphisms in the first intron of PGBD1 as a susceptibility locus for schizophrenia (SCZ) in independent studies^54–56^. Given high neuronal activity of the gene this may well be more than coincidence. Third, *prima facie* evidence suggests that it may not have evolved new functionality by adopting its prior functionality. In contrast to PGBD3 and PGBD4, PGBD1 does not have the transposase-derived open reading frame (ORF) flanked by transposon terminal inverted repeat sequences. Similarly, PGBD1, unlike it close relative PGBD2, is missing an intact C terminal cysteine-rich (CRD) domain. By contrast, unlike other PGBD genes, the transposase-derived ORF of PGBD1 is fused to an upstream exon of a SCAN-domain^46^ (named after Sre-ZBP, Ctfin51, Aw-1 and Number 18 cDNA).More generally, PGBD1 has been annotated as a protein of unknown function, consisting of a SCAN and a transposase-derived domain (Uniprot)^46, 49^. The SCAN-domain itself derives from a retrotransposon^57^, and functions as a protein multimerization domain, mediating homo- and hetero-oligomerisation^58, 59^, commonly found in the N-terminal regions of zinc-finger (ZNF) transcription factors^60^. The members of the SCAN domain family are expressed in neuronal cells^61, 62^. The addition of the SCAN domain we hypothesise may enable the evolution of functionalities, possibly in the neuronal context, not accessible to other PGBD domesticates. Here then we decipher the domesticated function of PGBD1 in humans with a view to better understanding its role in pathology.

## Results

### PGBD1 has an N-terminal SCAN/KRAB domain

Prior to functional assay, we sought to clarify the domain structure of the gene using sequence homology search and protein structure prediction (Phyre2 server).Phyre2 could detect the *PiggyBac* transposase domain (100% confidence, 20% sequence identity, template: <u>c6x68D</u>) and the SCAN domain, where the template of ZNF24-SCAN leads to the best match (100% confidence, 64% sequence identity, template: <u>c3lhrA_</u>). The transposase-derived domain includes a recognisable dimerization and DNA-binding domain (DDBD1/2) and catalytic sub-domain (DDD) (Fig. 1a,d).

**Fig. 1.**
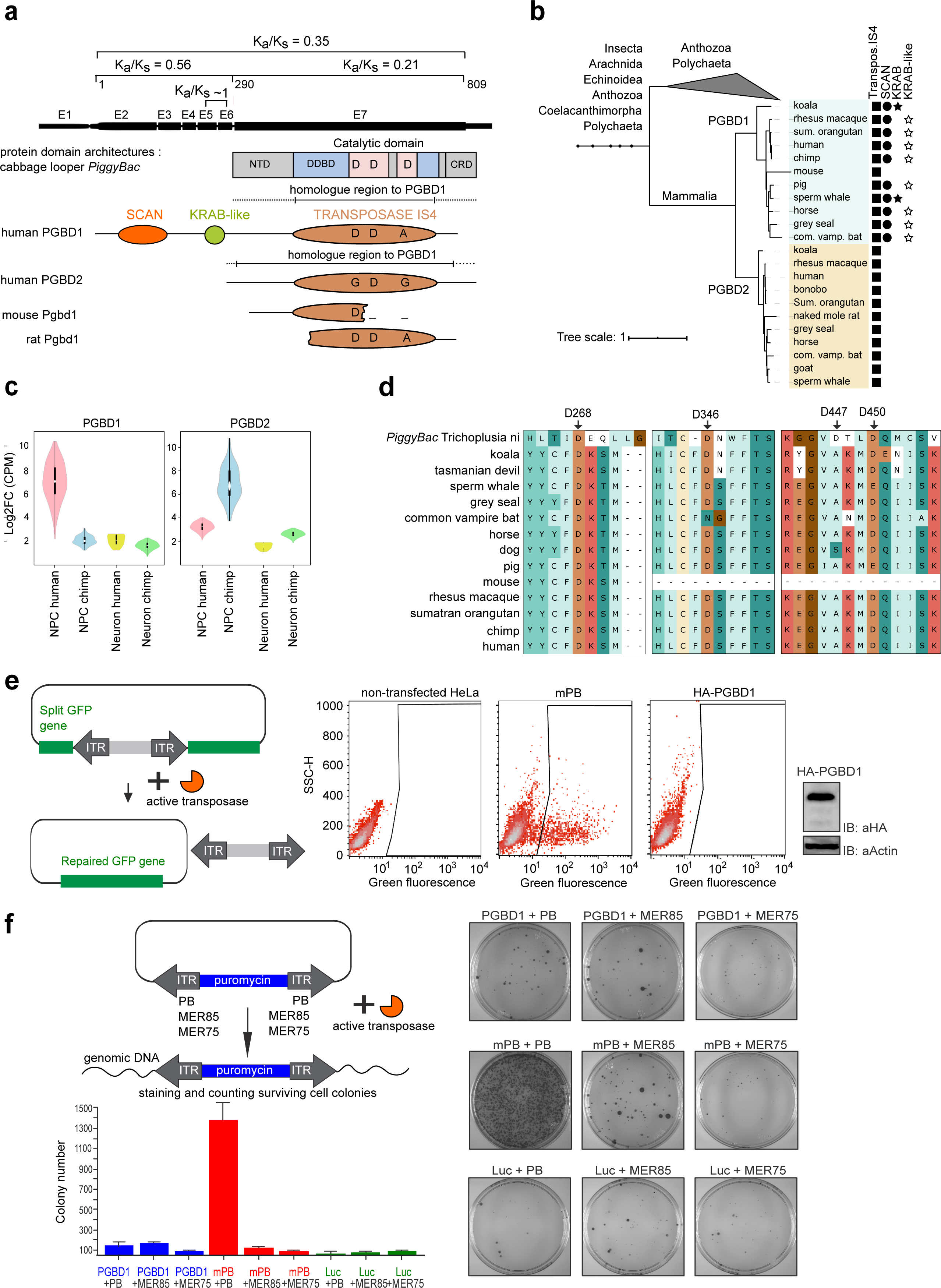
The domesticated PGBD1 possesses a SCAN-, KRAB- and transposase-derived domains, but has no catalytic activity as a transposase. **a,** PGBD1 domain structure in comparison to *PiggyBac* of the cabbage looper, human PGBD2, rat PGBD1 and mouse PGBD1. The transposase-derived domain (IS4) includes dimerization and DNA binding domains (DDBD) as well as the catalytic domains of PiggyBac^80^. NTD, N-terminal domain; CRD, C-terminal cysteine rich domain; E1-7 are exons 1 to 7. The ‘D’s in the transposase-derived domains represent the catalytic triad DDD (D268, D346, D447). D447 is replaced by A in PGBD1. PGBD2 and PGBD1 are highly similar (average pairwise similarity score of ∼ 63% the aligned region which spans 1324 bp exceeds the borders of the annotated transposase IS4 domain, calculated by distance matrix of Ugene). Note that the ZN-finger containing CRD domain, required for ITR binding in the piggyBac transposase is missing in PGBD1^223^. The PGBD1 sequences in rodent animal models are truncated, resulting in degenerated copies. The Ka/Kv values of the entire PGBD1 as well as for various subdomains are shown. Note the ∼1 value for the KRAB domain. (overall = 0.35, N-terminal (aa 1-290) = 0.56, C-terminal (aa 291-809) = 0.21, SCAN (aa 40-142) = 0.32, KRAB (aa 211-267) = 1.02, DDBD1 (aa 405-541) = 0.19, DDBD2 (aa 750-804) = 0.26, catalytic domain 1 (aa 541-651) = 0.14 and catalytic domain 2 (aa 726-750) = 0.07, reference is the human amino acid sequence of PGBD1). **b,** Phylogenetic tree of PGBD1 and PGBD2. The presence of the transposase-derived, the SCAN and KRAB domains are shown. The human PGBD1 and PGBD2, with the most closely related sequences (containing transposase IS4) were aligned with *muscle* and a tree was built using *MrBayes*. Protein domains were annotated with hmmerscan and CDD (NCBI). The KRAB domain was annotated with Phyre2. **c,** Relative expression levels of PGBD1 and PGBD2 in human and chimpanzee NPCs and neurons^73^(GSE83638). Note that the in cross-species comparison the expression of PGBD1 and PGBD2 is specifically enriched in human and chimp, respectively. **d,** Protein sequence alignment of the transposase-derived DDD catalytic domain of PGBD1. The first raw of the alignment shows the corresponding sequence of the *piggyBac* transposase, identified in *Trichoplusiani* (cabbage looper). The alignment includes koala and grey seal, from where the KRAB domain was reported (see also Fig. S1B), and various mammalian species. The conserved amino acids D268/D346/D447 of the conserved DDD catalytic domain and D450 of the *piggyBac* transposase are arrowed^46^. The numbers refer to their position using the piggyBac amino acid sequence as reference. **e,** *Transposon excision repair assay* detects no activity of the PGBD1. (Left panel) Schematic representation of the reporter assay of *PiggyBac* excision. The *PiggyBac* transposon (flanked by inverted terminal repeats, ITRs) splits the coding sequence of the green fluorescence protein (GFP) reporter. In the presence of an active transposase, transposon excision occurs, and the readout is the restored GFP reporter signal. (Middle panel) Quantitative FACS analysis of GFP positive cells generated in the transposon excision repair assay. HeLa cells were co-transfected with plasmids harbouring HA-tagged PGBD1 along with the reporter construct. Non-transfected HeLa and cells transfected with mPB (codon optimized, mouse *piggyBac* transposase) along with the reporter served as controls. (Right panel) Western blot analysis of the HA-tagged PGBD1 (HA-PGBD1) protein tested in the excision repair assay. **f,** *Transposition assay* detects no activity of the PGBD1 protein. (Left panel) Schematic representation of the colony forming transposition assay to detect stable integration of the puromycin resistance gene marked reporter in HEK293 cells. In case of active transposition, the transposase cuts at the ITRs - (inverted terminal repeats), and inserts the reporter-marked transposon into the genome, providing antibiotic resistance for the transfected cells. In addition to the piggyBac ITRs, reporters were also built with PiggyBac-derived miniature inverted repeat (MITEs) ITRs of MER75, MER85. See also Fig. 2a. (Right panel) Puromycin resistant HEK293 colonies as the readout of the assay. The constructs were transfected in various combinations. HEK293 cells transfected with the mPB transposase and the non-relevant Luciferase expression construct (Luc) along with the reporter served as controls. (Bottom Left panel) Quantification of the transposition assay. Colonies were quantified in a 75S model gel imager, using the Quantity One 4.4.0 software (Bio-Rad). Error bars indicate *s.d*. Note that the mPB transposase (positive control) was able to mobilize the PB transposon, but not the PB-inverted repeat related reporters.

The above accords with prior annotation. However, we also identified in some species an additional protein domain downstream of SCAN, a KRAB domain (Fig. 1a,b, Extended Data Fig. 1a,b). The KRAB domain is also part of the KRAB-ZNF (Zn-finger) family of sequence-specific transcriptional regulators, involved in cell differentiation and development^63^ (Extended Data Fig.1c). Automated protein domain search algorithms could detect the KRAB domain in the PGBD1 protein in only a few species (Fig. 1b). We investigated the homology of the KRAB-like region in a multiple sequence alignment (MSA) (Extended Data Fig. 1b) and predicted the protein structure with Phyre2. This aligns the KRAB domain (human PGBD1 query, template: <u>d1v65a_</u>) in the region aa 211-267 with a confidence score of 99.3% and 16% identity. We added the templates used by Phyre2 to our MSAs (Fig. 1a).

To decipher the evolutionary history of the conjugation between SCAN, KRAB and the transposase-derived domains seen in human PGBD1, we constructed a phylogenetic tree of all transposase IS4 domain containing sequences (∼12k), so as to capture all PGBD-related sequences. From these we extracted the subtree consisting of PGBD1, PGBD2 and some outgroup sequences (Extended Data Fig. 1b and Extended Data Fig. 1d). We detect bothPGBD1 and PGBD2from marsupials (diverged 148 Mya), but not before (e.g. Monotremes). The sequences of PGBD1 and PGBD2 are closely related, forming two well defined gene families, but differ most profoundly in their N-terminal domains (Fig. 1b). Species with the KRAB domain include marsupials (e.g. koala, Extended Data Fig. 1b, Extended Data Fig. 1d, 2a), supporting the hypothesis that KRAB inclusion is the ancestral condition but with numerous loss/decay events.

**Fig. 2.**
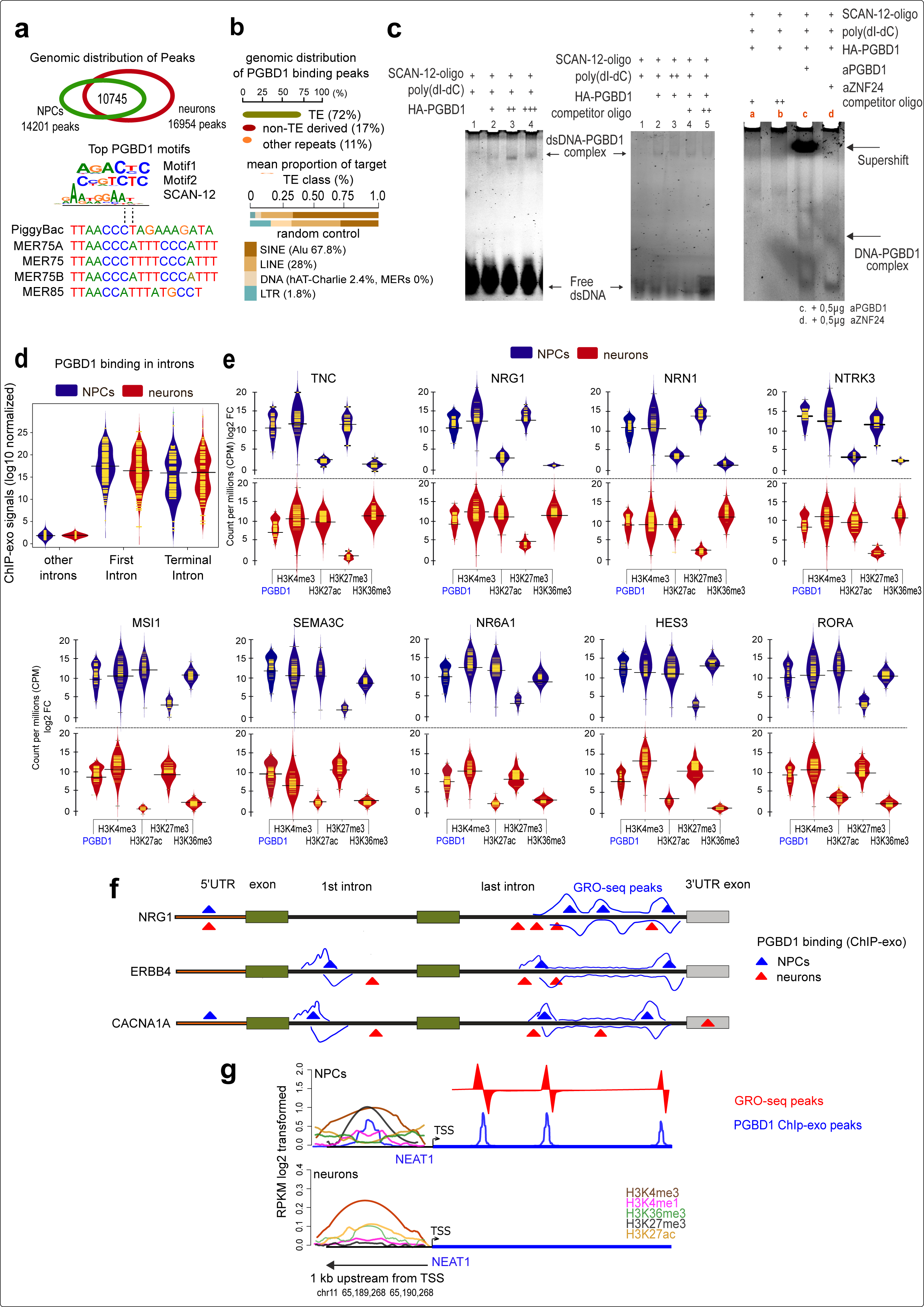
PGBD1 target genes influence human neural progenitor identity. **a,** ChIP-exo analysis of PGBD1 binding motifs with an overlapping, but specific distribution pattern in NPCs and in differentiated neurons. (Upper panel) Venn diagram shows the number of identified PGBD1 ChIP-exo peaks in hESC-derived NPCs, in hESC-derived neurons and common to both. (Middle panel) The top two PGBD1 motifs (Motif1 and Motif2) derived from over-represented sequences identified from both NPCs and neurons. SCAN-12 represents the shared consensus DNA-binding motif identified between PGBD1 and twelve SCAN-ZNF-proteins, including ZNF167, ZNF174, ZNF18, ZNF232, ZNF274, ZNF394, ZNF483, ZNF496, ZNF500, ZNF498, ZNF187, ZNF323. (Lower panel) Sequence alignment of the *piggyBac* inverted repeat sequences (ITRs) and ITR-like motifs of the *piggyBac* transposon-derived miniature inverted repeat elements (MITEs) of MER75A, MER75, MER75B and MER85. Note the lack of similarity between PGBD1 ChIP motifs and the *PiggyBac* or the MITE ITR sequences. **b,** PGBD1 binding is overrepresented on transposable elements (TEs). (Upper panel) Genomic distribution analysis reveals an overrepresentation (72%) of the PGBD1 binding peaks on TEs in NPCs, when compared to a random control (mean Z score =3.7). (Lower panel) Further dissection of TE-enriched PGBD1 ChIP-exo signals shows that the majority (67.8%) of the binding sites are mappable to *Alu* elements. **c,** Electromobility shift assay (EMSA) confirms that PGBD1 directly binds to the SCAN-12 consensus DNA motif.(Left panel) EMSA detects stable complexes formed between HA-PGBD1 (HA-tagged and purified) and SCAN-12 oligonucleotidesin the presence of nonspecific competitor (polydI-dC). (Middle panel) The stability of the complexes formed between HA-PGBD1 and SCAN-12 oligonucleotidesis not changed in the presence of increasing amount of the non-specific inhibitors but mildly decrease in presence of competitor oligonucleotides. (Right panel) In the presence of anti-PGBD1 antibody a supershifted complex is formed (c), confirming the specific binding of the SCAN-12 probe by HA-PGBD1. Note that the complex is resistant to form a supershifted complex with the anti-ZNF24 antibody (d), suggesting that HA-PGBD1 binds the SCAN-12 motif directly, and not via ZNF24. dsDNA, double stranded SCAN-12 oligo. **d,** Distribution of PGBD1 ChiP-exo binding signals in introns. Note that the ChIP-exo signals (log10 normalized, introns were length-normalized by plotting empirical densities of PGBD1 signals in bins of equal read count) were seen specifically in the first and terminal introns in both NPC and neurons. **e,** Binding of PGBD1 to the promoter region of selected target genes influence neural progenitor identity. Analysis of regulatory genomics of the promoter region (1kb upstream from TSS). Panels show the log2-Fold change (log2FC) values (CPM) of the PGBD1 binding signals (ChIP-exo) and epigenetic histone marks of a selected set of gene promoters in NPCs (blue) and in neurons (dark red). The selected genes are known markers of (Upper row) neuronal differentiation or (Lower row) cell proliferation maintenance. **f,** Representative examples of differential PGBD1 binding in NPCs and neurons. Differentially bound PGBD1 peaks between NPCs (blue triangle) and neurons (red triangle) at the NRG1, ERBB4 and CACNA1A genes (not to scale). Only first/last exons and 5’ upstream/downstream genic regions are shown. Blue wavy lines represent GRO-seq peaks in NPCs. **g,** PGBD1 binding suppresses NEAT1 expression in NPCs. PGBD1 ChIP-exo binding peaks (blue) and regulatory genomic analysis of the NEAT1 (1Kb upstream from the TSS) in NPCs and in neurons. Note that PGBD1 specifically binds NEAT1 in NPCs, but not in differentiated neurons. The NEAT1 promoter is in a repressed state in NPCs, supported by GRO-seq analysis (red), whereas activated in differentiated neurons.

### PGBD1 is unique to non-monotreme mammals

The above analysis suggesting that PGBD1 is mammal-specific agrees with the more limited EggNOG phylogenetic reconstruction^64^ (Extended Data Fig. 2a,b) and earlier analysis^49^. Apparent mammal-specificity may be consistent with incorporation of the gene in the mammals or with failure of homology search outside of mammals^65^. That, by contrast, PGBD5 (EggNog group ENOG5028KT9) has numerous non-mammalian vertebrate orthologs suggests that lack of evidence for homology of PGBD1/2 outside of mammals reflects absence rather than a lack of ability to resolve orthology.

To further consider this issue, we apply a recent method to determine whether absence of detectable homology is likely to be an artefact/failure of homology searching or real^65^. This method considers the likelihood that a gene would be called absent in a given taxon, given homology search relevant parameters (rate of evolution, gene size) and distance between taxa. It is tuned by parameter estimates from a minimum of three well described orthologs in species of known evolutionary distance. We find that absence of PGBD1 in monotremes and earlier lineages (e.g. reptiles) cannot be accounted for in terms of failure of homology search (probability of homolog detection failure=0 (E=0.001), 99% confidence interval, a=1705.0, b=1.2, r^2^=0.98) (Extended Data Fig. 3a).The same is true for PGBD2 (Extended Data Fig. 3a), consistent with the approximately synchronous colonization of the genome of the marsupial/eutherian common ancestor by PGBD1/2.

**Fig. 3.**
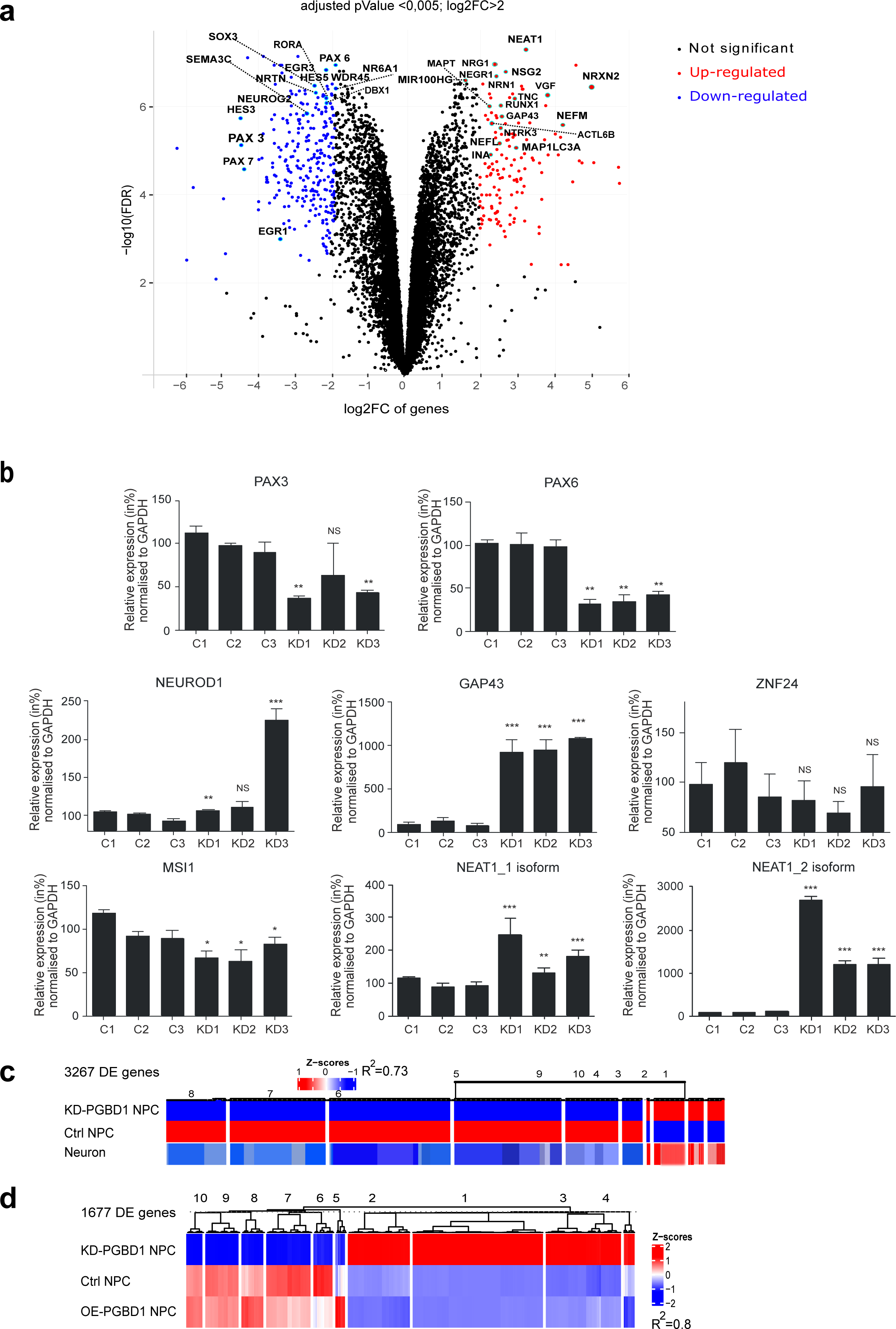
Depletion of PGBD1 compromises the progenitor state of NPCs. **a,** Volcano plot shows the 762 differentially expressed genes (DEGs) in the transcriptome of knockdown (KD) PGBD1 neural progenitors, KD-PGBD1_NPC (log2 fold > 2 change (log2FC), pValue< 0.05. Depletion of PGBD1 resulted in the up-regulation of 475 genes (red), whereas 287 genes were down-regulated (blue). The highlighted up-regulated genes are mostly associated with neural differentiation, whereas several of the down-regulated genes have relevant functions in maintaining the self-renewal of NPCs. (for the list of DEGs, see Supplementary Table 4). **b,** Quantitative polymerase chain reaction (qPCR) confirms the effects of PGBD1 knockdown on transcription of selected neuronal marker genes in NPCs. Data shown are representative of three independent experiments with biological triplicates per experiment. P-values: * p<0.05,** p<0.005, *** p<0.0005. Error bars indicate *s.d*. Note that while both isoforms are upregulated upon KD, the values are a magnitude higher for NEAT1_2 isoform. **c,** The transcriptomes of the PGBD1-depleted NPCs (KD-PGBD1) and differentiated neurons are highly similar (R^2^ = 0.73) (3267 DEGs). Global comparative analysis of the transcriptomes of PGBD1-depleted NPCs, untreated control (Ctrl) NPCs and differentiated neurons^222^. **d,** Overexpression (OE) of PGBD1 in NPCs does not generate robust transcriptome changes. The heat map shows the transcriptional changes of 1677 genes in KD-PGBD1, in OE-PGBD1 in comparison to scrambled-miRNA transfected control NPCs (Ctrl NPC). The transcriptome of the OE-PGBD1 NPCs are highly similar to the Ctrl NPCs (R^2^=0.8).

While within vertebrates we observed PGBD1 and PGBD2 in marsupials and eutherians exclusively (Extended Data Fig. 1b and Extended Data Fig. 1d), they nonetheless show homology to arthropod PGBD sequences (Extended Data Fig. 1d and Fig. 2b). The most likely explanation for the origin of PGBD1/PGBD2 sequences is thus horizontal transfer (rather than *de novo* origination) early in mammalian evolution. Such horizontal transfer is not without precedent for transposable elements^66–68^. The high similarity of PGBD1 and PGBD2 and their seemingly contemporaneous integration into the mammalian genome is consistent with a gene duplication event shortly after their integration. This duplication occurred either prior to the acquisition by PGBD1 of the SCAN domain or after the gain of the SCAN domain, but with immediate loss of the SCAN domain in PGBD2.Our analysis finds no evidence for the SCAN domain or homology across the relevant region in any PGBD2 ortholog (Extended Data Fig. 1b,d) supporting the first scenario. Evoking parsimony (minimum number of events) would support SCAN acquisition once by PGBD1 in the common ancestor of eutherians and marsupials shortly after the duplication (or parallel integration) event. Indeed, PGBD1 is found in a genomic domain rich in SCAN domain proteins suggesting gene fusion after integration into this site.

We conclude that PGBD1 incorporated in the common ancestor of marsupials and eutherian mammals and that this incorporation included the SCAN domain as the ancestral condition with KRAB inclusion also likely to be ancestral.

### The SCAN domain has been lost in model rodents and elsewhere

While we observe a relatively high homology between the transposase domains of the murine and human PGBD1s (87%) (Extended Data Fig. 3b), we could not identify the SCAN domain by HMMERsearch, nor could we observe homology to the SCAN domain within multiple sequence alignment when employing the annotated sequences of mouse and rat (Fig. 1a and Extended Data Fig. 1a,d, 2a). These findings suggest loss of the SCAN domain in some rodents.

To experimentally confirm the lack of the N-terminal SCAN/KRAB domains in the murine PGBD1s, we cloned the 5’ end of the rat Pgbd1 gene following reverse transcription of total RNA isolated from a rat cell line (Extended Data Fig. 3c,d). Instead of the SCAN/KRAB domains, molecular analysis of the rat Pgbd1 gene identified several STOP codons upstream of the transposase-like domain at the genomic locus (Extended Data Fig. 3e). Thus, while the SCAN-transposase domain fusion is present in the majority of mammalian genomes, it has been lost from murine rodents (Fig. 1a,b and Extended Data Fig. 1a,d). The murine loss can be dated to post the common ancestor between murines and fellow rodents, ground squirrels (Extended Data Fig. 2). Consequently, standard rodent models (e.g. mice/rat) are not suitable to study the function of (intact) human PGBD1. Broader and alternative phylogenetic analysis concurs with this rodent loss and suggests further losses of the SCAN domain in cats, grey lemurs and some marsupials (Extended Data Fig. 2 and Fig. 1d).

### PGBD1 is commonly under purifying selection

Given the flux in the structure of the gene with domains being lost or unrecognised across the mammals, we return to the issue of whether the gene is subject to purifying selection. Under the assumption that genes and domains that are functional are more conserved than expected, the gene should have Ka/Ks << 1, this being the hallmark of purifying selection^50^. To investigate evolutionary forces, PGBD1 mRNA (CDS) sequences from 11 mammalian and 18 primate species were analysed with PAML^69^.

For all regions, we detected an overall Ka/Ks ratio <= 0.35, indicative of purifying selection (Fig. 1a). For the KRAB domain, however, our analysis of branch-specific ratios revealed several organisms with Ka/Ks > 1 (horse, grey seal, common vampire bat and primates). To test if the evolution of the KRAB(-like) domain might be better explained by neutral or adaptive forces, we tested the two hypotheses and found that adaptive evolution explains the data better than neutral evolution (chi² = 7.47, df = 2, p-value = 2.38e-02).Given the broad evolution distances (and associated problems of synonymous site saturation), we repeated the analysis exclusively in primates (Ka/Ks ∼0.44 and 1.13, overall protein and KRAB region, respectively). We found that adaptive evolution was explaining the processes in this region significantly better (chi² = 7.43, df = 1, p-value = 6.42e-03) than neutral evolution. In particular, two sites were identified positively selected, beneficial mutations: 227V (prob ∼ 0.98) and 252M (prob ∼ 0.96, naive empirical bayes, reference: human full-length aa sequence). While 252M is typically not well conserved in other KRAB domains, and its functional relevance is unknown, position 227 is a well conserved F in other in other KRAB domains, and is important for binding the transposable element suppressor TRIM28^70, 71^. We conclude that the gene is functional and mostly under purifying selection but in one subregion in some lineages it is functional and subject to positive selection for reasons unknown.

### PGBD1 is highly expressed inhuman neuronal progenitors but not in chimp

A key question to understand the functions of any gene are the foci of high expression. In humans, both PGBD1 and PGBD2 show a broad range of expression in most tissues, with PGBD1 expression enriched in the cerebellum and the cerebellar hemisphere of the brain, consistent with their frequent involvement in neuronal disease^72^. PGBD2, by contrast, is mostly expressed in spleen and thyroid tissues (Extended Data Fig. 4a,b).

**Fig. 4.**
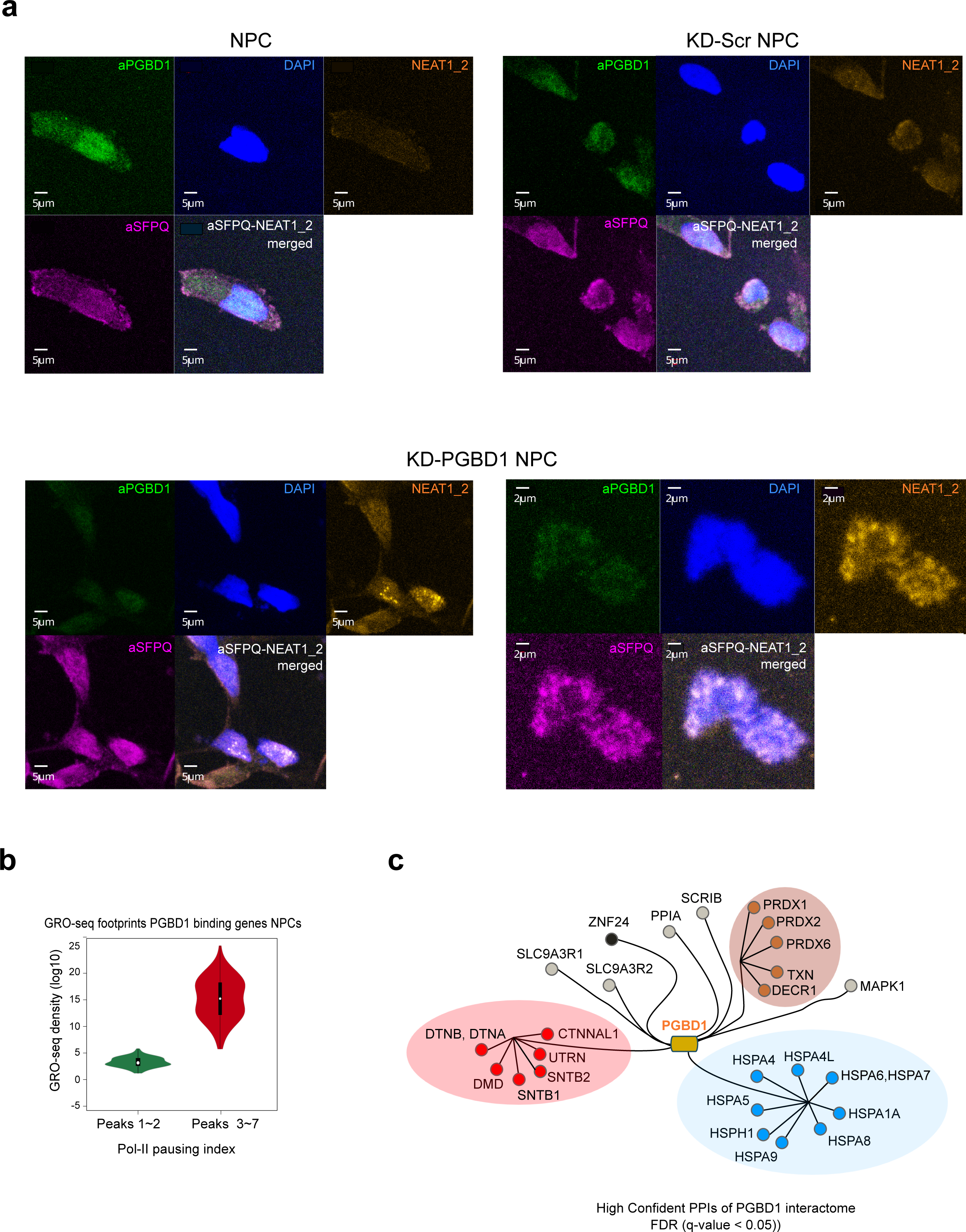
The protein interacting partners of PGBD1 include structural proteins of paraspeckles. **a,** Depletion of PGBD1 induces paraspeckle formation of in NPCs. Representative fluorescent microscopy images using antibodies against PGBD1 (green), SFPQ paraspeckle (purple) proteins, combined with FISH visualisation of NEAT1_2 RNA (yellow-brown) and DAPI (blue) staining. (Upper panels) No significant level of NEAT1_2 RNA-FISH signal is detectable in (Left panel) untreated NPCsor (Right panel) in scrambled-miRNA transfected KD-Scr NPC controls. Note the colocalization of the aPGBD1 and aSFPQ signals mostly in the nuclei (merged image). (Lower panels) Representative images of paraspeckle formation in KD-PGBD1 NPCs (at two different magnifications). In sharp contrast to controls, upon PGBD1 depletion, a robust NEAT1_2 RNA-FISH signal appears that accumulates in speckles (yellow-brown). The NEAT1_2 RNA-FISH signal colocalization with the aSFPQ immunostaining (merged image) define the nuclear structures paraspeckles. **b,** GRO-Seq plot demonstrating global pausing indices of transcriptionally active RNA polymerase II (Pol-II) overlapping PGBD1 binding sites with peaks (N=1-2) compared to (N=3-7). Note that the pausing index in genes having multiple PGBD1 peaks was higher ∼4.1, when compared with genes having less peaks ∼1.2. **c,** The high confidence protein interaction partners of PGBD1 (Log2FC(H/L ratio) L-HA-PGBD1<-2,0 and Log2FC(H/L ratio) H-HA-PGBD1>2,0) identified by MS-SILAC (MDC-PGBD1 PPIs). PPI: protein-protein interactor. PPIs of similar function are colour coded: Blue, chaperones; Brown, oxidation/reduction status modifiers; Red, member of the dystrophin-associated proteins (DAGs); Black, SCAN-ZNFs; Grey, unclassified.

To narrow down the neuronal activity profile, we analysed PGBD1/2 expression levels in neuronal progenitors (NPCs) and in differentiated neurons of human and chimpanzee (GSE83638) ^73^. Both PGBD1 and PGBD2 have a higher expression in progenitor vs differentiated cells (Fig. 1c). The observed expression pattern of PGBD1 in human NPCs vs neurons is consistent with regulatory genomics analyses over the PGBD1 genomic locus (Extended Data Fig. 4c). Notably, upon differentiating human embryonic stem cells (hESCs) to neurons, PGBD1 is expressed highest (both RNA and protein) in hESCs, followed by NPCs and its differentiated derivatives (Extended Data Fig. 4d,e). The relative enhancement of PGBDs in NPCs is, however, species-specific (e.g. PGBD1-human; PGBD2 -chimp), suggesting functional diversification during primate evolution (Fig. 1c).

### The human PGBD1 has no detectable transposase activity

Transposable elements have transposase activity key to their successful genomic colonization. This same functionality can be co-opted on domestication^38, 39^. We thus sought to test for activity of the catalytic domain of PGBD1. Sequence considerations suggest that the activity might have been lost. In PGBD1, we observe a replacement (by A) at the third D within the DDD motif of the catalytic domain (D447A) (Fig. 1d). As the DDD motif is highly conserved^46, 74^,and mutation at D447 abolishes catalytic activity^46, 74^, reduced functionality is to be expected. This loss is seen in koala and Tasmanian devil (Fig. 1d), suggesting that this loss was an early event.

To test for (potential residual) transposase activity, we designed two assays. First, we used a tissue culture-based *excision assay*^75^ that restores the open reading frame of the GFP reporter, following the excision of the piggyBac (PB) transposon (Fig. 1e). As PGBD1 has no obvious inverted repeats (IRs), flanking the transposase-derived sequence, we utilised the piggyBac IRs. Using a mouse codon-optimized mPB transposase as a positive control, we detected no transposon excision activity in the presence of the human PGBD1 (Fig. 1e). As the assay requires precise excision of the transposon, we also performed a less restrictive *transposition assay* (Fig. 1f). For both assays, in addition to the piggyBac IRs, we generated additional reporter constructs, where we used IR sequences, flanking the piggyBac-derived genes of PGBD3 and PGBD4 that have been also amplified as <u>M</u>iniature Inverted-repeat <u>T</u>ransposable <u>E</u>lements (MITEs) of MER75B or MER85, respectively in the human genome^76, 77^ (Fig. 1f and Extended Data Fig. 4f). However, no detectable catalytic activity of the PGBD1 was observed using any of the reporters while the positive controls worked as expected.

Collectively, these results indicate that the catalytic function of the transposase-derived domain in PGBD1 has indeed inactivated, consistent with the D447A replacement in the DDD motif. Given that this D->A event is an early post integration event, any domesticated function is unlikely to be connected to the catalytic activity of the transposase.

### PGBD1 does not recognize *piggyBac* transposase recognition sequences

The transposase function of *PiggyBac* transposons enables recognition of certain DNA motifs. With the transposase inactive in PGBD1 we can ask whether it still recognizes known *PiggyBac* recognition motifs or other motifs and if, so how. The SCAN domain might provide alternative functionality, PGBD1 being classified as a SCAN cerebellum-specific transcription regulator (GTEx portal) and Supplementary Table 1). Structurally, several members of this family also possess a KRAB domain followed by a ZNF domain (Extended Data Fig. 1c). In contrast, in PGBD1, the sequence-specific ZNF domain is replaced by the PB transposase-derived region with a sequence-specific binding potential (Extended Data Fig. 1c).

To determine whether PGBD1 has sequence-specific DNA binding, we performed a ChIP-exo assay in both NPCs and neurons, differentiated from the human embryonic stem cell line, hESC_H1^78, 79^. The assay identified two top ChIP-exo motifs of PGBD1, indicating its sequence specific binding capacity (Fig. 2a). Notably, despite the shared motifs, the individual binding peaks show distinct NPC- and neuron-specific binding sites with an overlap (Fig. 2a), suggesting that PGBD1 has both a shared and cell type-specific binding pattern in NPCs and neurons.

First, we checked if the identified ChIP-exo motifs have any similarity to the *piggyBac* transposase recognition sequences, located in the inverted repeats (IRs) of the transposon or the IRs, flanking the PGBD3 and PGBD4 genes (also MER75B and MER85). However, we found no obvious similarity between the PGBD1 ChIP-exo motifs and the IRs (Fig. 2a), suggesting that PGBD1 does not bind *piggyBac* transposase-related recognition sequences in the human genome.

### PGBD1 genomically does not recognise piggyBac-related sequences

Having defined what PGBD1 does not bind, we sought to identify what it does bind. We analysed PGBD1 peaks genome wide (Extended Data Fig. 5). Given that it does not recognise itself, we asked whether it might recognise other transposable elements, particularly those with sequence resemblance. We observed no enriched intersection at PB-like MER75/85 sequences^76, 77^. However, we find a relative enrichment (72%) of TEs compared to a random generated control (Fig. 2b). On questioning whether the co-occurrence was more common for a particular TE family-derived sequences, we found that the TE/PGBD1 binding sites were enriched in *Alu* elements (Fig. 2b and Extended Data Fig. 5b) (subfamily *AluSx*) as against the general distribution of TE classes (χ2 test p value <0.01). We can conclude that PGBD1 has lost ability to recognise itself or related sequences, consistent with its loss of the CRD^49^, this domain being key to terminal inverted repeat recognition^80^.

**Fig. 5.**
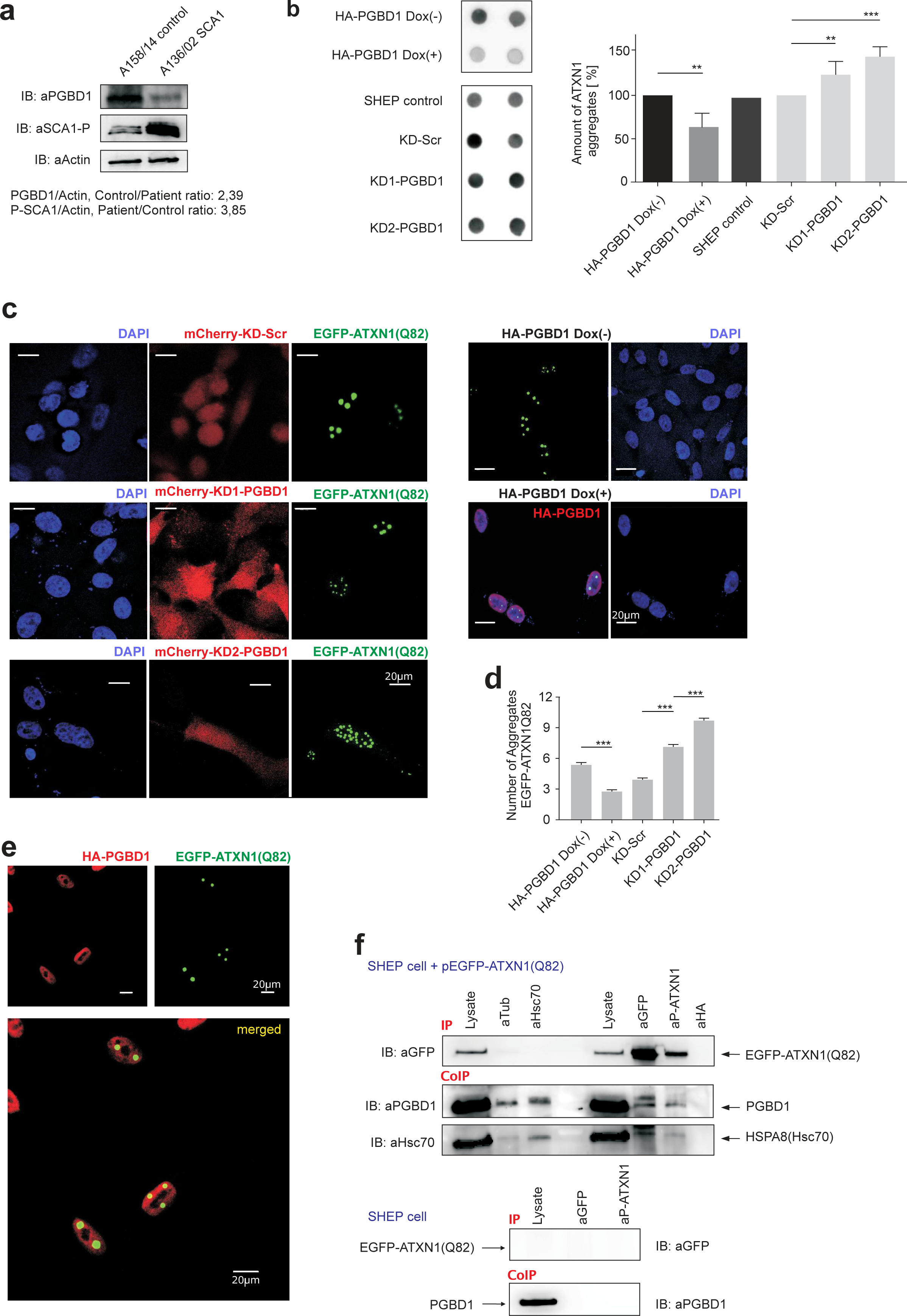
PGBD1 dissolves protein aggregates in the polyQ-Ataxin1 protein aggregation model. **a,** Decreased protein level of PGBD1 in spinocerebellar ataxia type 1 (SCA1) patients. Western blot analysis shows PGBD1 protein levels in human brain cerebellum homogenates derived form a healthy (Acc.Nr: A158/14) and a SCA1 patient (A136/02). IB, immunoblot; aSCA1-P, antibody against the phosphorylated (phospho S776) Ataxin 1 (ATXN1); The ratio of the aPGBD1/aActin and the aSCA1-P/aActin signal ratios are indicated in the panel. In the SCA1 patient, note the increased and the decreased levels of phosphorylated ATXN1 (aSCA1-P) and aPGBD1 signals, respectively. **b,** Detection of SDS-insoluble nuclear aggregates of ATXN1(Q82) using *filter retardation assay* in SHEP neuroblastoma cells. (Left panel) Induction or knocking-down (KD) effects of PGBD1 on the formation of spontaneous ATXN1(Q82) nuclear aggregates in SHEP cells. (Upper box) SHEP cells, stably expressing hemagglutinin (HA)-tagged PGBD1 from an (DOX)-inducible expression construct transfected with the EGFP-tagged ATXN1(Q82) reporter. (Lower box) SHEP cells, stably expressing various RNAi constructs, KD-Scr, KD1-PGBD1 and KD2-PGBD1 transfected with the ATXN1(Q82) reporter. UN, un-transfected control. (Right panel) Densitometric quantification of insoluble ATXN1(Q82) aggregates detected using *filter retardation assay* in SHEP neuroblastoma cells, presented on the right panel by AIDA Image Analyser software. Scr, scrambled control. DOX(+) doxycycline induced and (DOX-) non-induced Student’s t-test, ** p<0.005, *** p<0.0005. **c,** Depletion of PGBD1 results in an elevated number of spontaneous EGFP-tagged ATXN1(Q82) aggregates in SHEP neuroblastoma cells. (Left panel) Detection of EGFP-ATXN1(Q82) aggregates by confocal microscopy. Red, mCherry-tag on the RNAi constructs: Scr, scrambled; green, EGFP-tag on the ATXN1(Q82); blue, DAPI. SHEP cells stably express both EGFP-ATXN1(Q82) and the RNAi constructs, mCherry-KD-Scr, mCherry-KD1/2-PGBD1.(Right panel) Induced HA-PGBD1 expression results in a decreased number of spontaneous ATXN1(Q82) aggregates in SHEP neuroblastoma cells. Green, EGFP-tag on the ATXN1(Q82); red, anti-HA-PGBD1; blue, DAPI. Note that the induced PGBD1 efficiently dissolves nuclear protein aggregates. Sale bar, 20 µm. **d,** Quantification of the aggregation assay. Student’s t-test, *** p<0.0005. **e,** PGBD1 co-localizes with the EGFP-ATXN1(Q82) aggregates in the nuclei of SHEP cells. Representative confocal microscopy image of the HA-tagged PGBD1 and EGFP-tagged ATXN1(Q82) aggregates in human SHEP cells. Green, EGFP-tag on the ATXN1(Q82); red, anti-HA-PGBD1. **f,** PGBD1 recruits the Hsc70(HSPA8) chaperon molecule to nuclear protein aggregates formed by ATXN1(Q82). (Upper panel) Immunoprecipitation (IP) and Co-immunoprecipitation (CoIP) identifies the endogenous PGBD1 as well as Hsc70(HSPA8) in the nuclear protein aggregates formed by ATXN1(Q82) in SHEP cells. SHEP cells are transfected with pEGFP-ATXN1(Q82) and subjected to analyses. IB:immunoblot. (Bottom panel) Control immunoprecipitation experiment by using non-transfected SHEP cells.

### PGBD1 genomically binds protein coding genes and the consensus motif of SCAN domain family members

Many of the *AluSx* elements are intronic^81, 82^. More generally, in both NPCs and neurons, we observe binding sites mapping predominantly in or around (+/- 1 kb) protein coding genes (∼84%) (Extended Data Fig. 5a) (Z=4.2), rather than non-coding regions (∼16%). Generally, in genic regions we commonly observed multiple binding peaks on targeted genes that were mappable to introns and upstream regulatory regions (1kb from TSS). Indicative of regulatory roles, the PGBD1 ChiP-exo signals were exclusively distributed in either the first and last introns (NPC, 51 and 49%; neurons, 50% each), 5’ and 3’ UTRs, frequently associated with regulatory features (e.g. transcriptional regulation, alternative splicing/polyadenylation)^83–85^. By contrast, no peaks were mappable to introns located more centrally in the gene bodies (Fig. 2d).

This focus on gene bodies but not internal introns is consistent with a regulatory role. If so, we might also expect a concentration in near gene regulatory regions as well. We therefore asked if the PGBD1 Chip-exo peaks in non CDS map to regulatory regions, frequently associated with active enhancers. Intersecting PGBD1 Chip-exo peaks, genomic regions >2kb upstream TSS of RefSeq annotated genes and H3K4me1 histone mark signals revealed ∼2000 PGBD1 binding peaks (68% NPC, 72% neurons) overlapping with regulatory regions genome-wide (Extended Data Fig. 5e). *AluSx* was enriched in this set against a randomised background (Z=3.8). Taken together this evidence suggests that PGBD1 has evolved gene regulatory functions.

Via their multimerization domain, the proteins of the SCAN-domain family might interact with each other^86, 87^. Indeed, PGBD1 has been reported to have several protein interaction partners of the SCAN family (e.g. SCAND1, ZKSCAN1,3,4,8; ZSCAN1,12, 18,20,22,25,32) (BioGRID, STRING)^57, 88, 89^. Is this possible partnership reflected in DNA binding? Data mining of reported binding motifs of the SCAN-ZNF transcription factors predicts a 9 bp sequence, shared by 12 ZNF proteins (SCAN-12) and PGBD1 (Fig. 2a), suggesting that several family members can bind the same consensus, potentially modulating binding. 40% of PGBD1 peaks correspond to this 9 bp sequence.

To confirm that PGBD1 can bind the consensus, we performed electrophoretic mobility shift assay (EMSA) with fluorescently labelled oligonucleotides, containing the consensus SCAN-12-motif and purified human HA-tagged PGBD1 protein. EMSA supported a stable DNA substrate-PGBD1 association (Fig. 2c). Can PGBD1 bind the consensus SCAN-12 motif alone or only in cooperation with other SCAN family members, possibly co-purifying with PGBD1? To address this, we added antiPGBD1 antibody to the reaction mixture. The observed super-shifted complex suggests that PGBD1 is able to specifically bind the SCAN-12-motif alone, even without the other SCAN-ZNF family members (Fig. 2c, right).

### PGBD1 provides access to both repressing and activation factors at regulatory regions of target genes involved in neural progenitor maintenance

What is the function (if any) of PGBD1 binding to DNA and what genes are associated with such binding? In both NPCs and neurons, among the most significant ontology (GO) terms of targeted protein coding genes we observed *neuron development*, *neuron differentiation and neurogenesis* (Extended Data Fig. 5c and Supplementary Table 2).

Since the ChIP-exo peaks shared the binding motifs, but had specificities in NPCs and neurons (Fig. 2a), we hypothesized that the sites would have differential accessibility. To test this hypothesis, we first analysed the common target genes in both cell types around their transcriptional start sites (TSSs) (1 kb upstream) for various histone marks of active and repressive activities in promoter regions (e.g. H3K4me3, H3K36me3, H3K27me3, H3K27ac) and chromatin accessibility (ATAC-seq) (Extended Data Fig. 5d.e). This integrative analysis revealed a set of common target genes, where the PGBD1 peaks overlapped with either activating or repressive histone marks (Extended Data Fig. 5f).

To determine what the repressed and activated biological processes are, we selected a set of common target genes from the most significant GO categories e.g. *nervous system development* (NPC: n=887) and *neuronal differentiation (*NPC: n=545), and determined their chromatin activation status. This revealed that the PGBD1-targeted gene set which overlaps with activating histone marks include several genes associated with the maintenance of the progenitor state (e.g. MSI1, SEMA3C, NR6A1, HES3, RORA), whereas the repressed genes are generally involved in inhibiting NPC differentiation (e.g. TNC, NRG1, NRN1, NTRK3) (Fig. 2e and Supplementary Table 2).This suggests that in NPCs PGBD1 is involved in keeping the regulatory regions of genes responsible for neural progenitor maintenance in an active state, while simultaneously repressing those that could initiate the differentiation process. These results are consistent with PGBD1 maintaining the progenitor state of NPCs.

In the common gene set of PGBD1 targets in NPCs and neurons, we observe both shared and differential binding peaks (Supplementary Table 2), supporting the observation that PGBD1 binding might exert differential, cell type specific modulation of gene activity. In addition to the genes of distinct accessibility at their upstream regulatory regions (Fig. 2e), among the targets with a differential intronic binding pattern, we identify NRG1 (neuregulin 1) and its associated cell surface receptor, ERRB4 (Erb-B2 Tyrosin kinase 4) or CACNA1A (Calcium Voltage-Gated Channel Subunit Alpha1 A), all required for normal development of the embryonic central nervous system^90–93^ (Fig. 2f).

While the identified NPC-specific targets did not show up in any GO category significantly, we observed several lncRNAs among the repressed non-protein coding genes, including NEAT1, DICER1-AS1, ATP1A1-AS1, CCDC18-AS1. As NEAT1, a hallmark of differentiated cells, is not observed in embryonic stem cells or in neural progenitors^27, 94^, PGBD1 binding at the NEAT1 locus might be associated with its suppression (Fig. 2g).

What might PGBD1 be doing that is specific to neurons, and not seen in NPCs? We identified 191 stress response genes from the GO categories of *homeostatic process regulator* and *cellular stress regulator*. Notably, PGBD1 binds 23% (44/191) of these stress response genes at their upstream/regulatory region. This is consistent with PGBD1 regulating response to environmental stress in neurons (Supplementary Table 3).

### PGBD1 helps maintain the fate identity of human NPCs

To test for PGBD1 target gene activation/repression obtained from the integrative analysis of ChIP-exo and histone mark data, we performed transcriptome analysis on PGBD1-depleted NPCs. As the knockout (KO) strategy interfered with cell renewal (Extended Data Fig. 6a), preventing stable maintenance of a colony, it was more feasible touse a knockdown (KD) approach (KD-PGBD1) to deplete PGBD1 (Extended Data Fig. 6b-d). A comparison of the NPC and KD-PGBD1_NPC transcriptomes (Log2-fold change, L2FC) identified 763 differentially expressed genes (DEGs) (Fig. 3a and Supplementary Table 4), some of them also validated by qPCR (Fig. 3b).

**Fig. 6.**
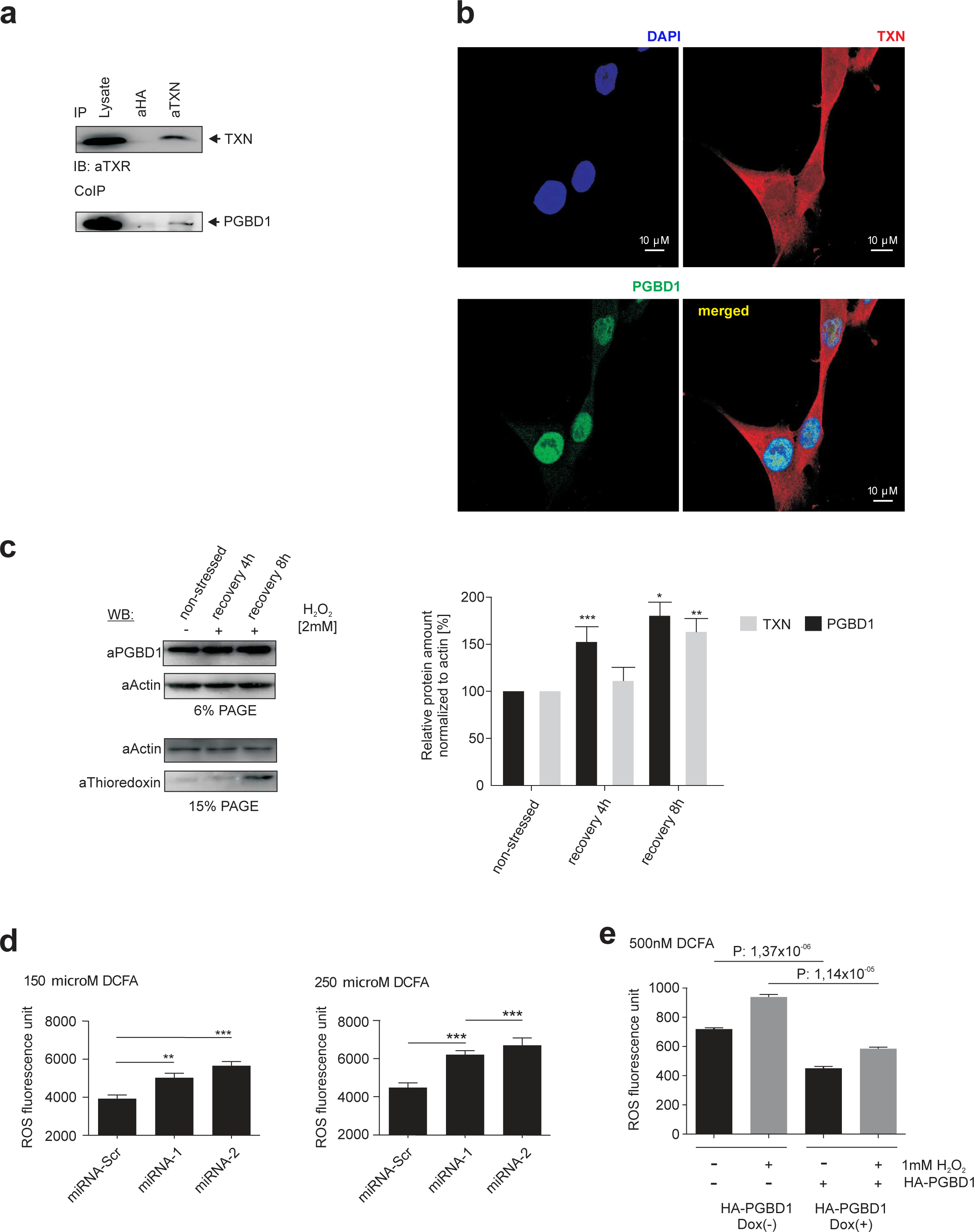
The presence of PGBD1 reduces reactive oxidative species (ROS). **a,** PGBD1 physically interacts with thioredoxin (TXN). Co-immunoprecipitation assay used to confirm the PGBD1-TXN protein interaction. (IB: immunoblot, IP: immunoprecipitation reaction, CoIP: co-immunoprecipitation, Lysate: undiluted cell lysate). **b,** Co-localisation of the endogenous PGBD1 and TXN protein in human neuroblastoma SHEP cells. Confocal microscopy. Red-Alexa Flour® 555: TXN; green-Alexa Flour® 488: PGBD1; blue, DAPI. **c,** Increased PGBD1 level is detected in SHEP cells recovering after oxidative stress exposure. (Left panel) Western blot (WB) analysis of the protein levels of PGBD1 and TXN in SHEP cells exposed to oxidative stress. (PAGE: polyacryl-amide gel electrophoresis, % shows the PAGE gel concentration). (Right panel) Densitometric quantification of the relative levels of PGBD1 and TXN proteins between stressed and non-stressed conditions visualized on Western blot. * p<0.05, ** p<0.005, *** p<0.0005. Note that the PGBD1 and TXN levels increased after 4h and 8 h recovery, respectively. **d,** Depletion of endogenous PGBD1 by RNAi (miRNA-Scr/1/2) results in elevated ROS level. FACS measurement of the ROS signals were determined in the stable knock-down PGBD1 SHEP cells using different DCFA concentrations. (Left panel) 150 μM DCFA; (Right panel) 250 μM DCFA. **e,** Elevated level of HA-tagged PGBD1 protein provides a protection against H_2_O_2_ (1 mM) exposure in human SHEP cells. Measuring of ROS extent in stable, DOX-inducible SHEP cells expressing HA-tagged PGBD1 protein. * p<0.05.

The overall GO analysis of the DEGs in KD-PGBD1_NPC suggests that the function of PGBD1 is associated with *gene regulation of cell differentiation, nervous system development* and *neurogenesis* (Supplementary Table 4). Importantly, from the 763 significant DEGs, we found 212 genes (over 1/3), with significant PGBD1 ChIP-exo signals consistent with differential expression being caused directly by PGBD1 DNA binding (see above). In this overlapping dataset, the peaks for PGBD1 binding are located in the upstream regulatory (1kb upstream from TSS) regions of 38 DEGs, again consistent with PGBD1 playing a transcriptional regulatory role of these targets.

Analysis of the 38 genes revealed the down-regulation of SEMA3C, NR6A1, HES3, RORA, NRTN and DBX1 genes in KD-PGBD1, implicated in maintaining the proliferative status of NPCs^95–97^(www.rndsystems.com/research-area/neural-stem-cell-and-differentiation-markers). Among the upregulated genes, by contrast, we identified genes associated with neuronal differentiation (e.g. TNC, NRG1, NRN1, ACTL6B and NTRK3) (Fig. 3a). Notably, (lnc)NEAT1 (primarily lncNEAT1_2), the structural RNA component of paraspeckles, was among the most significant DEGs in KD-PGBD1 and differentiated neurons, but lncNEAT1_1 was also affected, both validated by qPCR (Fig. 3a,b). This supports a key role of PGBD1 in suppressing transcription of the NEAT1 locus in NPCs, consistent with binding data. Among the DEGs we identify another essential lncRNA MIR100HG (Fig. 3a), implicated in neuronal differentiation regulation (e.g., encoding for the miRNA cluster, including LET7a-2)^98, 99^.

Collectively, our transcriptome analysis, in conjunction with the integrative chromatin status determination, suggests that the depletion of PGBD1 compromised the identity of NPCs, and triggered the cells to activate their differentiation program. To test this hypothesis, we compared the transcriptome of NPCs, KD-PGBD1_NPCs and *in vitro* differentiated neurons from hESCs (4 weeks)^79^. The comparison revealed a strong correlation (R^2^=0.73) between the transcriptomes of PGBD1-depleted NPCs and differentiated neurons, and anticorrelation between NPCs to both, supporting a high similarity between the transcriptomes of KD-PGBD1 and differentiated neurons (Fig. 3c). In addition, analysing the transcriptional changes of 99 key neuronal markers supports the hypothesis that depleting PGBD1 drives cells toward a differentiated phenotype (Extended Data Fig. 6e). Thus, PGBD1 depletion activates NPC differentiation, arguing for an essential role of PGBD1 in the maintenance of the progenitor state of neuronal cells. In contrast to the robust transcriptional changes generated by PGBD1 depletion, overexpression of PGBD1 results in no dramatic changes in NPCs (Fig. 3d), and the OE-PGBD1 transcriptome stays close to the control (R^2^=0.8).

Finally, in order to probe whether PGBD1 influences expression of additional SCAN family member(s), we also determined the transcriptional status of other SCAN domain family members in PGBD1-depleted cells. Among the significant DEGs, we found only the brain/cerebellum-specific SCAN-KRAB domain ZNF483 with moderate expression alteration (L2FC-1.79), while other family members were not affected transcriptionally (not shown). This observation is in line with the assumption that the family members are more likely to modulate each other’s activity via protein-protein interaction (via the SCAN domain).

### PGBD1 inhibits paraspeckle formation in neural progenitor cells via transcriptional control of NEAT1

The longer NEAT1 isoform, NEAT1_2, localizes to paraspeckles^100^ and has an essential role in their formation, the tRNA-like triple helix at its 3’-end being required to stabilize paraspeckles. Given the specific DNA binding of PGBD1 at the NEAT1 locus (Fig. 2g) and the regulatory effect of PGBD1 on NEAT1_2 transcription (Fig. 3b), we hypothesise a role of PGBD1 in the regulation/biogenesis of paraspeckles, among other functions.

In order to test this prediction, we used confocal microscopy to visualize NEAT1, PGBD1 and SFPQ (splicing factor proline glutamine-rich protein) as a paraspeckle marker, in WT and PGBD1-depleted NPCs. To visualize NEAT1, we performed a FISH assay by using a specific probe for the NEAT1_2 transcript. We combined the FISH with immunohistological staining against PGBD1 and SFPQ. Our expectation under a model in which PGBD1 suppresses NEAT1_2 is that in the presence of PGBD1 (which should be intranuclear and diffuse), we should not observe SFPQ/NEAT1_2 colocalised foci, the SFPQ should be intranuclear and diffuse and NEAT1_2 largely absent. On PGBD1 depletion, SFPQ and NEAT1_2 should now colocalise in intranuclear foci (paraspeckles).

In WT-NPCs and in scrambled control cells, no obvious signs for either elevated level of NEAT1_2 transcripts or SFPQ-marked paraspeckles were seen (Fig. 4a). Both PGBD1 and SFPQ are predominantly nuclear and diffusely distributed as predicted. These observationsare consistent with previous reports that paraspeckles are not detectable in NPCs^94^. In PGBD1-depleted NPCs, by sharp contrast, we detect a robust nuclear NEAT1_2 signal, indicating intensive NEAT1_2 transcription. The NEAT1_2 signal accumulated in multiple SFPQ-marked nuclear bodies, colocalized with SFPQ immunostaining, which is no longer diffuse (Fig. 4a). These observations indicate that a decreased level of PGBD1 is associated with extensive paraspeckle formation (Fig. 4a), consistent with a key suppressor role of PGBD1 in the biogenesis of paraspeckles in NPCs. We also repeated the visualization experiment in a stable, PGBD1-depleted neuroblastoma cell line (SHEP) with a similar result (Extended Data Fig. 8). These data suggest that the release of the NEAT1_2 transcription from PGBD1-mediated suppression is a requirement of paraspeckle formation.

How does PGBD1 enforce such an effective suppression of NEAT1_2 transcription? A detailed integrative analysis (e.g. PGBD1-ChIP-exo; histone marks, ChIP-seq; RNA-seq) finds, besides the upstream regulatory region of NEAT1, three further PGBD1 ChIP signals over the gene body in NPC (Fig. 2g), whereas no significant binding signal was detectable over the NEAT1 locus in differentiated neurons. In NPCs, PGBD1 efficiently inhibits NEAT1 transcription, likely by forming a ribonuclear/protein complex as a physical barrier that interferes with both TF binding at the regulatory region of NEAT1 and its POL2-mediated transcription. Removing the suppression of NEAT1 by knocking down PGBD1 in NPCs results in a robust upregulation of NEAT1_2 in the transcriptome (∼17-fold elevation) (Fig. 3b), suggesting that PGBD1 plays a key role in repressing NEAT1_2, and thus paraspeckle formation.

To determine whether PGBD1 functions as a transcriptional barrier at multiple targets, we performed a genome-wide GRO-seq analysis in NPC. Protein coding target genes bound by PGBD1 were divided into two groups based on the number of the observed ChIP-exo peaks (1-2 vs 3-7). In order to quantify transcriptional pausing for regions of interest, we computed the quotient of binned expression per analysed feature (PGBD1 intersecting GRO-seq peaks) and the entire gene body. Our findings (density of GRO-seq reads) (Fig. 4b) suggest that regions with multiple PGBD1 binding sites are prone to transcriptional pausing, consistent with blocking NEAT1 transcription.

### Protein interacting partners are enriched for dystrophin associated proteins, chaperones and stress response genes

The above concerns the role of PGBD1 as a DNA binding protein. At least via its SCAN domain, PGBD1 is expected to also have multiple potential interaction protein partners. What processes, if any, might these interactions mediate? As a first step to evaluate this we determined PGBD1 global protein interactome, and performed a differential mass spectrometry-based proteomics approach, SILAC (**S**table **I**sotope **L**abelling by **A**mino acids in **C**ell culture) in HEK293 cells.

SILAC-based quantitative affinity purification mass spectrometry (q-AP-MS) of overexpressed hemagglutinin peptide (HA) tagged PGBD1 and non-tagged PGBD1 overexpressing control cells quantified1,103 proteins in both a forward and a reverse (label swap) experiment. From these, we identify 19 highly confident interacting partners of PGDB1 (defined as FDR < 0.05) that also have expression >= 10 TPM in NPCs (Fig. 4c, Extended Data Fig. 7a and Supplementary Table 5).

**Fig. 7.**
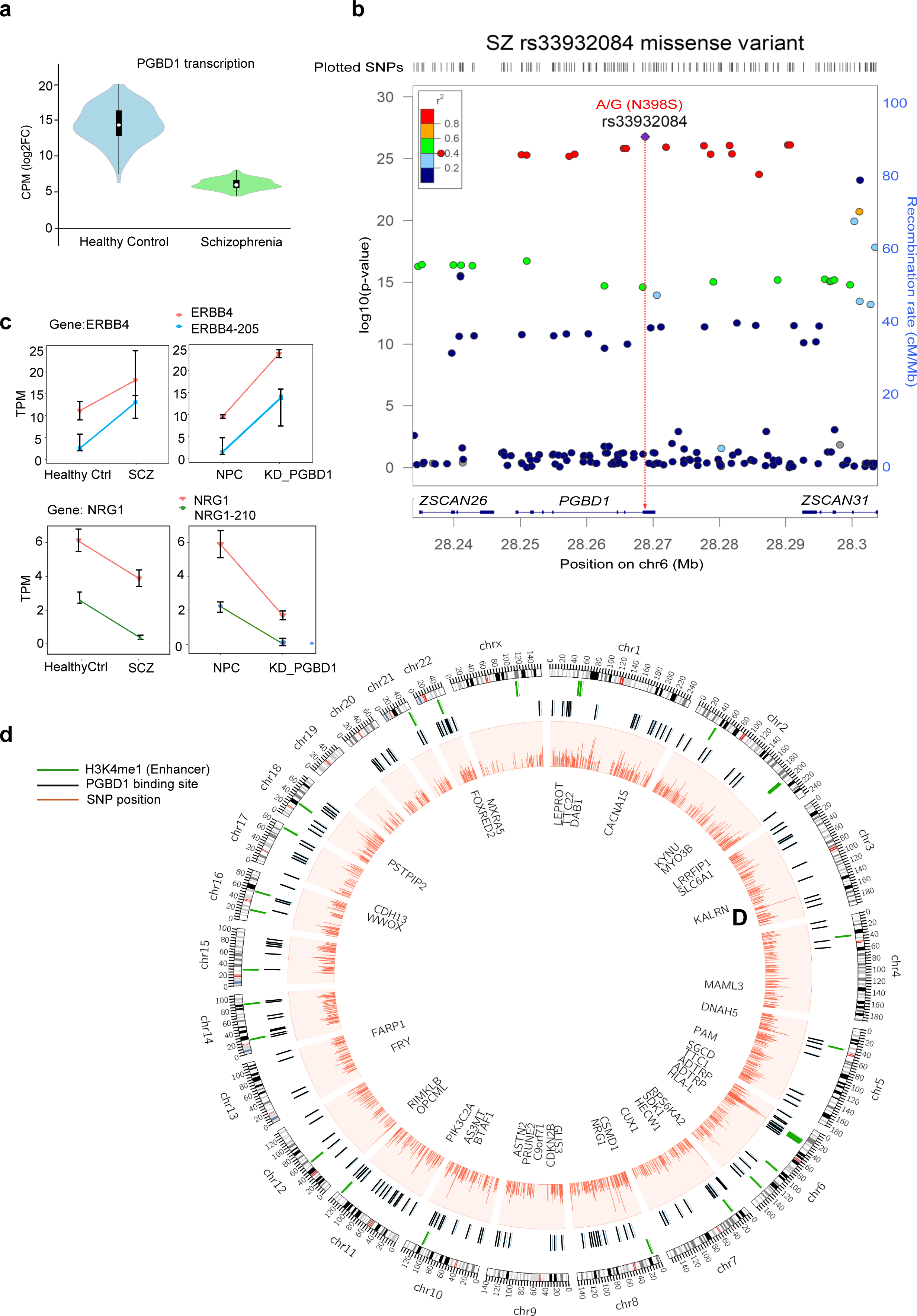
The KD-PGBD1_NPC model mimics certain aspects of schizophrenia. **a,** Transcription levels of PGBD1 in healthy control (HC) and schizophrenia (SCZ) patients (GSE118313 and GSE121376). **b,** Schizophrenia (SCZ) associated SNPs mapping on the chr6 in and around the PGBD1 gene. SCZ patients (N = 443,581) and a replication cohort (1169 controls; 1067 cases)^224^. Y axes shows the strength of association (log10-p-value). Colour code: LD r^2^. Rs33932084 was identified in last exon of PGBD1, resulting in a missense mutation (N398S). **c,** The *in vitro* model of PGBD1-depleted NPC (KD-PGBD1_NPC) mimics certain features of schizophrenia. Comparative isoform analysis of NRG1 and ERBB4 derived transcripts between various contrast groups (NPC, neural progenitor cell; NPC KD-PGBD1, PGBD1 depleted NPC; HC, healthy control; iPSC-derived neurons from schizophrenia patients). Note that both NRG1 and ERBB4 genes have similar transcript isoform-enrichment in PGBD1-depleted NPC and in schizophrenia patient-derived cells. **d,** PGBD1 binding is enriched at the enhancer regions of a subset of schizophrenia (SCZ) susceptibility genes. SCZ-associated SNPs (N=1251), PGBD1 ChiP-exo peaks (N∼2000), mapping >2kb upstream of TSS of genes were overlaid with the H3K4me1 histone marks (indicative of active enhancers). CIRCOS shows the schizophrenia susceptibility genes indicated with PGBD1 binding in their enhancer region.

The SILAC experiments involved overexpression of either HA-tagged PGBD1 or untagged PGBD1. Hence, there was overexpression in both cases, so we don’t expect a systematic bias to be introduced by over-expression per se. Nonetheless, to address whether their transcript abundance might be affected by PGBD1 expression levels, we determine the changes in NPCs where PGBD1 was overexpressed or knocked down. Overexpression (OE) of PGDB1 elevated the abundance of transcript for HSPA4/A4L and SNTB2.From these HSPA4L and SNTB2 were also affected by knockdown (Extended Data Fig. 7b), whereas the remainder of the high confidence interacting partners of PGBD1 show a clear knockdown-specific pattern (Extended Data Fig. 7b).

As expected of a SCAN domain protein, PGBD1 does indeed bind at least one member of the SCAN family. Among the high confidence interacting partners of PGBD1, we identified ZNF24 (Zn-finger protein 24), a SCAN-ZNF family member (Fig. 4c, Extended Data Fig. 1c and Supplementary Table 5). The gene is implicated in maintaining the neural progenitor fate^61^ and is expressed at >70TPM in NPCs. Neither TRIM28, a KRAB interactor, nor any of the members of the TRIM28-associated gene regulation complex, are within the set of significant interactors.

The interactome of PGBD1 however is by no means restricted to SCAN domain proteins. The dystrophin associated protein (DAP) complex comprises at least ten proteins^101^, from which, we identify (e.g. Dystrophin (DMD), Dystrobrevin (DTNA, DTNB), Uthrophin (UTRN), Syntrophin beta 1 and 2 (SNTB1, SNTB2), Catenin alpha like 1 (CTNNAL1) in the interactome of PGBD1. While, dystrophin is primarily expressed in the skeletal muscle, DTNB is a member of the brain-specific dystrophin complex, expressed in the spinal cord, the hippocampus and within the cerebral cortex^102, 103^.The brain-specific DTNB regulates distinct aspects of neurogenesis, including terminal differentiation (reviewed in^104^). The specific interaction of PGBD1 with multiple DAPs suggests that PGBD1 could have a significant modulatory effect on the processes, controlled by the brain-specific dystrophin associated protein complex.

In addition to the DAP complex, two further gene set clusters are evident (Fig. 4c). Using gene ontology (*GO*)-based classification, DMD, SNTB1, SNTB2 and UTRN are also belong to the *Go glycoprotein complex.* This analysis identifies *Go chaperon cofactor dependent protein refolding*, r*eactome regulation of Hsf1 mediated heat shock response* and *reactome cellular response to heat stress* as the most significant categories (Extended Data Fig. 7c). Indeed, there are significant interactions with multiple stress-responsive chaperones, involved in protein quality control (e.g. HSPA4, HSPA4L, HSPA5, 6/7, 8, 9, HSPH1, HSPA1A)^105, 106^ (Fig. 4c and Extended Data Fig. 7c).

### PGBD1 dissolves nuclear protein aggregates via recruited chaperones

An association with a diversity of chaperones may be little more than reflecting the need for PGBD1 to fold correctly or indeed the notion that chaperones tend to be sticky proteins. Nonetheless, given an association with a diversity of chaperones, we asked whether, via its associated chaperones, PGBD1 might generally affect the polymerization/aggregation properties of proteins. To test this hypothesis, we employed nuclear Ataxin1 protein (ATXN1) as a protein aggregation model^107–111^. In the neurodegenerative disease of spinocerebellar ataxia type 1 (SCA1), the ATXN1 protein, has a pathologically expanded polyglutamine (polyQ) region and susceptibility for a spontaneous misfolding^112^.A possible dissolution of the nuclear aggregates is thought to depend on chaperone activity^105,113,114^.

Importantly, ATXN1, similar to PGBD1, is expressed in the cerebellum (Gtex/ATXN1), and PGBD1 protein levels were lower, relative to a healthy control in a human cerebellum homogenate derived from a SCA1 patient (A136/02) (Fig. 5a), suggesting that the ATXN1 model might be a highly relevant system to test PGBD1 function.

Our aggregation model consisted of fluorescently labelled ATXN1 reporters with expanded polyQ track (Q) of increasing lengths (Q2/30/82) in stable human neuroblastoma SHEP cell lines. We asked if knocking-down (KD) or overexpressing (OE) PGBD1 affected the aggregation of the reporter proteins. First, we performed a cellulose acetate membrane-based filter retardation assay. This approach revealed that depleting PGBD1 results in an increased amount of Q82 aggregates (Fig. 5b). We also determined the KD/OE PGBD1 effects on the number of Q82 aggregates using confocal fluorescence microscopy. These experiments visualize that KD-PGBD1 cells accumulate an increased number of protein aggregates, whereas the excess of PGBD1 results in a reduced number of nuclear inclusions formed by both Q82 and Q30 (Fig. 5c, d and Extended Data Fig 9). Collectively, the aggregation assays suggest that the presence of PGBD1 mitigates the improper folding of Q30 and Q82 variants in a chaperone-like mechanism. Using confocal microscopy, we observe that PGBD1 co-localizes with the nuclear aggregates of EGFP-ATXN1 (Q82) (Fig. 5e).

To find out whether PGBD1 specifically recognizes the ATXN1 or the aggregate, we performed co-immunoprecipitation (co-IP) experiments using the Q82 variant of ATXN1 and Q2 as a control. The co-IP experiments revealed that while PGBD1 co-precipitates with the Q82 variant (Fig. 5f), it does not form a stable complex with the minimal Q2 (Extended Data Fig. 10a). Furthermore, PGBD1 did not affect the number of aggregates formed by EGFP-ATXN1(Q2) Q2 (Extended Data Fig. 10b-d). Thus, PGBD1 has an affinity to the misfolded protein structure (e.g. the polyQ aggregate) and not to ATXN1 itself.

To see if a chaperone molecule is present in the aggregate together with PGBD1, as suggested by the proteome, we performed a co-IP experiment between ATXN(Q82), PGBD1 and a chaperone, HSPA8/HSC70, previously predicted to be an interacting partner of PGBD1 by SILAC based AP-MS. HSPA8/HSC70 was detectable in both forward and reverse co-IP experiments with PGBD1 and EGFP-ATXN1(Q82) (Fig. 5f), demonstrating that PGBD1 and HSPA8(HSC70) are both present in the aggregates.

While these experiments are not suitable to eliminate the possibility that PGBD1 might act as a chaperone itself, it is more likely that the chaperone(s), recruited by PGBD1 are responsible for the observed effect. Therefore, PGBD1 appears to dissolve the nuclear aggregates by recruiting nuclear chaperones.

### PGBD1 is involved in oxidative stress mitigation via its protein interacting partners

Dissolving aggregations (via recruited chaperones) could also contribute to mitigate nuclear stress in both NPCs and neurons. The interactome also suggests some coupling with a stress response. More generally, there appears to be a complex stress response/paraspeckle coupling. Indeed, beside differentiation^115, 116^, paraspeckles are potential markers of cellular stress, including oxidative stress^117^. Furthermore, NEAT1was shown to deplete members of the peroxiredoxin family of antioxidant enzymes PRDX1,5,6, and mitigate stress in various cell types^118, 119^. It is surely no coincidence that, among the significant potential interacting partners of PGBD1 we find PRDX1/2/6 and thioredoxin (TXN) (Fig. 4a) (validated by additional co-immunoprecipitation and co-localization immunofluorescence microscopy studies) (Extended Data Fig. 6a and 6b), implicated in regulating redox signalling^120, 121^. In addition, we identify SCRIB (Fig. 4a), which interacts directly with the NADPH oxidase (NOX) complex and is required for NOX-induced reactive oxygen species (ROS) generation in culture and *in vivo*^122^. SCRIB additionally determines cell polarity and cell proliferation. It is involved in synaptic function, neuroblast differentiation, and epithelial polarization^123^.

To find out, whether PGBD1modulates the oxidative stress response, we first tested whether oxidative stress activates PGBD1 expression (Fig. 6c and Extended Data Fig. 11). To determine this, human SHEP neuroblastoma cells were treated with hydrogen peroxide. We monitored both PGBD1 and the antioxidant TXN protein (interacting partner) levels in a short recovery period of 4 and 8 hours post-treatment. After a short oxidative stress exposure, the level of both PGBD1 and TXN proteins increases rapidly in the first 8 hours (Fig. 6c). We observed that the speed of PGBD1 accumulation in the treated cells is similar (even faster) when compared to TXN (Fig. 6c). To see whether PGBD1 modulates oxidative stress signalling, we determined reactive oxygen species (ROS) (Extended Data Fig. 11)^124^ in knockdown/overexpressing (KD/OE) SHEP cells (Extended Data Fig. 6d and 6e).Using this assay in stable KD-PGBD1 cells, we observed a significantly elevated ROS signal, whereas a tendency of decreased ROS signal is detectable upon induced PGBD1 expression (Fig. 6e). These findings suggested that PGBD1 was capable of both sensing oxidative stress and eliminating toxic oxidative elements from cells.

### PGBD1 and neuronal pathologies

NEAT1 is a negative regulator of neuronal excitability and axonal maintenance^125^, which are the hallmarks of neurodegenerative disorders. Dysregulated NEAT1 expression has been reported from Amyotrophic lateral sclerosis (ALS), Huntington disease (HD), Alzheimer’s disease (AD), Parkinson’s disease (PD)^33, 125–127^and schizophrenia (SCZ)^128^. PGBD1’s association with NEAT1 provides a potential coupling with neuronal disease. Might then PGBD1 also be involved in neuronal diseases?

PGBD1 is a highly transcribed gene in the human cerebellum, followed by the cortex (Extended Data Fig. 4a). The dysfunction of both regions is connected with several neuronal disorders, including aggregation disorder, such as SCA1, but also schizophrenia (SCZ), whose pathophysiology includes disturbed cerebellar/cortical circuits^129–132^. Due to its expression profile and its role in maintaining neuronal homeostasis, including cellular stress mitigation in differentiated neurons, impaired PGBD1 function is a likely candidate associated with neuronal pathologies. In line with this, the expression level of PGBD1 is decreased in SCA1 (Fig. 5a), and mutations of a PGBD1 target, CACNA1A (Fig. 2f) are associated with neurological disorders of ataxia (Spinocerebellar Ataxia 6 and Episodic Ataxia 2)^133^.

Mining of GSE118313 and GSE121376 data revealed^134^ a reduced PGBD1 level also in schizophrenia (SCZ) patients (Fig. 7a)^135–137^, suggesting its potential association with SCZ.However, while independent genome-wide association studies (GWASs) identified several PGBD1-associated SNPs (rs2142731, rs1150772, rs3800324, rs13211507)^55, 138–140^ in SCZ patient cohorts, and PGBD1 has been previously mentioned among the potential candidate list of risk genes of schizophrenia, the significance of these findings was somewhat weakened by the facts that (i) the suggested SNP-SCZ association (e.g. rs2142731) was not confirmed in all ethnic groups^56, 141, 142^, and that (ii) the identified SNPs are located in an intronic region of PGBD1, with no obvious mechanistic explanation as to its potential contribution to schizophrenia.

Were analysed a large-scale single nucleotide polymorphism (SNP)/schizophrenia GWAS data^139^of 36,989 cases and 113,075 controls collected in multi-ethnic populations, but using a narrowed 25 Mb window around the MHC region (that now includes the PGBD1 locus). This strategy supports PGBD1 as a high susceptibility gene of schizophrenia (Fig. 7b). Notably, this analysis revealed rs33932084 (chr6:28,268,824) as a missense mutation (N398S) in the transposase-derived domain of PGBD1 (Fig. 7b), predicted as a potential regulatory SNP (rSNP) in a large-scale integrative analysis of two SCZ-GWAS datasets with functional annotation datasets^143^.

Do we find schizophrenia susceptibility genes among PGBD1 target genes? In fact, PGBD1 binds several reported susceptibility genes, including ERBB4, NRG1, ATXN3, SNAP91 and SRPK2 (Supplementary Table 2)^135–137, 139, 144–146^. Notably, these genes are from the overlapping cluster of PGBD1 target genes between NPCs and neurons and have distinct binding patterns in NPCs and neurons (Fig. 2e), suggesting their differential expression regulation during development. Certain transcript isoforms of NRG1 and ERBB4, specifically enriched in the neurons of schizophrenia patients^134^ are also accumulated upon PGBD1 depletion in NPCs (Fig. 7c). This suggests that the *in vitro* KD-PGBD1_NPC depletion model might mimic certain features SCZ.

An open question is the extent to which any effects of PGBD1 in schizophrenia might act through the NEAT1 axis. Despite PGBD1 suppressing NEAT1 expression in NPCs, in schizophrenia the expression of both NEAT1 and PGBD1 are reduced^134^ (Fig. 7a). While NEAT1_2 is a structural RNA for paraspeckles, both NEAT1_1and _2 RNA isoforms have DNA binding properties^147^. We might therefore also suspect dysregulation along the NEAT1 controlled transcriptome. Given antagonism between PGBD1 and NEAT1 (at least in NPCs), we ask whether there is an overlap between PGBD1 and lncNEAT1 target genes. Indeed, a comparison of PGBD1 target genes in NPCs with human NEAT1-ChIRP-seq data^128^ confirmed a large set of commonly targeted genes (75) involved in neurogenesis (21) and neuronal development (19) (Supplementary Table 6). Of these 75 common target genes 68 are SCZ susceptibility genes (Jaccard index=0.67). Specifically, PGBD1 targets those SCZ susceptibility genes involved in Oligodendrocytes development (Supplementary Table 6). Aberrant development of Oligodendrocytes is suggested as a hallmark of SCZ^148–151^, with NEAT1 also being suggested as a SCZ modulator via oligodendrocytes transcription^128^. Consistent with some coordination, both PGBD1 and (lnc)NEAT1 bind the target genes at the same positions, namely transcriptional start and termination sites (TSS and TTS)^147^. This pattern is different from MALAT1/NEAT2 (TTS only) that co-localize on many target genes with NEAT1^147^.

SNPs in enhancer regions might contribute to dysfunctional gene regulation of their targets^152, 153^. Thus, finally we asked whether a potential association exists between the enhancer-promoter (EP) gene regulatory network of PGBD1-bound genomic sites and SCZ susceptibility. We analysed the promoter interactions of PGBD1-bound, H3K4me1-enriched putative enhancer regions (∼2000) (Extended Data Fig. 5e) in the HACER database (<u>H</u>uman <u>Ac</u>tive <u>E</u>nhancers to interpret <u>R</u>egulatory variants), and intersected them with SNPs, identified by SCZ GWAS^139^. This revealed that around 53% of the target genes interacting with PGBD1-bound enhancer regions (1982 and 2101 in NPCs and neurons, respectively) are also associated with schizophrenia susceptibility, defined by GWAS risk SNPs studies^154^ (Fig. 7d).

Collectively, in NPCs, PGBD1 is required for the maintenance of the neural progenitor status (e.g. by antagonising NEAT1 expression), whereas in differentiated neurons, it has a stress mitigation regulatory function, shared with that of lncNEAT1(e.g. via targeting an overlapping set of target genes).

## Discussion

While we know little in detail about recently domesticated TEs, one might expect that their co-option would typically employ their catalytic abilities and that, being new genes, they might normally control just one function via one mechanism of action. PGBD1 confounds all of these expectations: PGBD1’s function is not related to the catalytic activity of a transposase, it has multiple functions and does so via multiple modes, i.e. DNA binding and protein-protein interactions. As regards its functions it is, however, not so unusual. For example, through its unfolded structure^155, 156^, the transposase-derived domain is likely involved in recruiting proteins involved in stress response (e.g. chaperones), mirroring classical TE role in stress responsiveness^157^. Similarly, like other domesticated TEs (i.e. HERVH)^19^, PGBD1 has a role in self-renewal regulation.

The loss of potential DNA binding activity and the gain of new activity is at first sight surprising for a domesticated gene. However, recent analysis has suggested that chimeric TE – KRAB-SCAN genes are a common mode of TE domestication^158^. PGBD1 present a paradigmatic example of such a process. The loss of catalytic ability is in line with the assumption that the DDD catalytic domain of the *piggyBac* transposase is not conserved among the domesticated *piggyBac*-derived PGBD sequences^46, 48, 159^ and hence probably not key to their domestication^49^. This contrasts with PGBD5 that has been suggested to have a residual DNA transposase-like activity, capable of mobilizing a synthetic DNA transposon in human cells, resulting in genome-wide DNA damage and genomic rearrangements^51, 160, 161^.

### A mammal-specific gene to control a mammal-specific structure, paraspeckle

The fact that depletion of this gene compromises self-renewal of the neural progenitors in human suggests that this is an unusual case in which a new gene has evolved a cell-level core function. Such circumstances are intrinsically paradoxical as we must query how organisms survived before the evolution of the new core gene. In this instance the resolution appears to be, at least in part, that the new gene is associated with a new core process. Indeed, it is likely to be significant that a mammal-specific gene controls a mammal-specific structure, paraspeckles.

The new mammalian gene PGBD1 and new mammalian structure (lnc)NEAT1/paraspeckles are indeed very closely intertwined and appear to regulate similar biological processes, including differentiation and stress responses in humans^94, 115, 162^. As regards co-regulation, we observe a significant overlap between PGBD1 DNA targets and genes regulated by (lnc)NEAT1binding^128, 147^. The PGBD1-mediated inhibition of paraspeckle formation supports a continuously stable condition for self-renewal of neural progenitors. In PGBD1-depleted cells, the robust elevation of NEAT1_2 transcription results in the appearance of paraspeckles, and the cells are set to differentiate^94^.

The biological function of PGBD1 is not, however, restricted to support the neural progenitor identity, and further functions of PGBD1 overlap with roles previously associated with (lnc)NEAT1/paraspeckles.Similar to (lnc)NEAT1/paraspeckles, the stress responsiveness of PGBD1 includes sensing and handling cellular stress (e.g. oxidative stress or protein aggregation stress)^163, 164^. PGBD1 sequesters central protein components of these biological processes in a cloud/hub of its interactome. Via its recruited chaperone molecules, PGBD1 is able to maintain proteins in their soluble form. Whether, in addition to a transcriptional control of NEAT1, PGBD1 controls paraspeckle biogenesis by regulating polymerization properties of its structural proteins, prone to polymerase and assemble around the NEAT1_2 to form the higher order paraspeckle structure, is yet to be deciphered. Nevertheless, remarkably, PGBD1 is even capable of dissolving protein aggregates in the nucleus (in contrast to several recently identified proteins that initiate phase separation^165, 166^).

Notably, PGBD1 is specific for regulating (lnc)NEAT1/paraspeckle biogenesis, and other nuclear bodies, namely nuclear speckle and CS body assembled around structural lncRNAs of MALAT1(NEAT2)^167^ and GOMAFU^168^, respectively,are not affected.Furthermore, the PGBD1-mediated regulation of paraspeckles might be specificto neural cells of the cerebellum, where PGBD1 is dominantly expressed, whereas paraspeckle regulation by other means (e.g. CARM1 (coactivator-associated arginine methyl-transferase1)^169^ might be more typical in other cell types.

Mammalian specificity associated with control of a mammal-specific structure, paraspeckles, implies coevolution of new genes with new functionality. Taxonomically restricted duplicate genes associated with novel physical structures^170^ have also been described. Generally, however, well described examples of new genes associated with new structures (macroscopic or microscopic) are scarce. Perhaps the best comparison could be made to the repeated evolution TE derived syncytins involved in the formation of the multinucleated syncytiotrophoblast protoplasm of mammalian placenta^40^. However, in contrast to PGBD1, the evolution of syncytins was mediated by co-option of the fusogenic capacity of the parental *env* gene^40^. In PGBD1 the current biological activity has little resemblance to the ancestral activity, not least owing to the loss of transposase catalytic abilities and gain of function (e.g. by recruitment of SCAN/KRAB).

### The enigmatic evolutionary plasticity of PGBD1

Perhaps the most enigmatic feature of PGBD1 is that it is both core to a fundamental process (NPC self-renewal) in humans and evolutionarily labile. The knockout cells can be extracted but lose their progenitor status (Extended Data Fig. 6a). Not only are young genes typically “non-essential”^171^, typically “essential” genes have hallmarks of conservation not of turnover^172^. The plasticity is seen on several fronts: presence/absence of domains (both SCAN and KRAB) across the mammals, positive selection in the KRAB domain and expression fluidity that we observed comparing closely related primates.

Such evolutionary turnover is often owing to antagonistic coevolution^173, 174^. While interactions (suppression) of other transposable elements may be one cause of positive selection within the KRAB domains as TRIM28, among the most prominent interaction partners of KRAB, is also involved in TE suppression^175–177^. However, the mutation 227V (which is a well conserved F in other KRAB domains) is important for binding the transposable element suppressor TRIM28 binding, suggesting thatPGBD1 is under selection not to enable transposable element suppression (possibly including itself). Consistent with such a loss, we find no evidence from protein interaction data for classical KRAB-TRIM28 binding, similar to other observations of a lack of binding when KRAB is in the SCAN/KRAB context^89^. Furthermore, PGBD1 has affinity to bind other TEs (e.g. *Alu* elements).Specifically, we observed that PGBD1 recognizes *AluSx,* these being a subfamily of *Alu* that inserted after the divergence of the Old World and New World monkeys^178^. The significance, if any, of this primate-specific binding affinity or whether it is associated with the evolution of paraspeckles in primates, are yet to be elucidated.

Alternatively, such evolutionary flexibility could reflect gain or loss of functionality in different mammals owing to specific aspects of their development or ecology. It is possible, for example, that PGBD1 became essential only in human neural progenitors, and the biological processes associated with the domesticated PGBD1 (e.g. improving the fidelity of environmental interactions^179^ (DAP complex), fine-tuning adaptive behaviour in response to physiological stress (paraspeckle regulation)^180^,proved to be crucial to support the neural homeostasis of the evolving cerebellum. Indeed, the cerebellum is an unusual focus of selection within the great apes^181^. As both SCAN and KRAB are missing in rat Pgbd1, and even the transposase-derived domain is partially deleted in mouse, functional characterization of the protein in these species may be informative, explicitly because they are not models for the human protein.

### PGBD1 is a clinically relevant new gene

Given that the gene has so many functions and modes and is cell-level essential for self-renewal, it is, in retrospect, reasonable that PGBD1 might be disease-associated. The association with neuronal disease follows from its neuronal functionalities. First, PGBD1 represses a set of target genes, involved in inhibiting the neural differentiation process^95, 182, 183^. Second, in neurons, PGBD1 is involved in mitigating stress response. Curiously, PGBD1 functions in a partially overlapping mode to (lnc)NEAT1, butmodulates a set of additional target genes to (lnc)NEAT1 targets, consistent with PGBD1 being a key regulator of stress response in the cerebellum. Its ability to maintain the cellular homeostasis of neural progenitors and to mitigate stress response in the differentiated neurons make the functionally impaired/attenuated PGBD1 as a clinically relevant factor to contribute to neurodegenerative diseases.Our study spotlights a possible association of the reduced level of PGBD1 with Spinocerebellar ataxia type 1 (SCA1), a nuclear polyQ aggregation disease^111^. Via its interaction with the dystrophin associated protein complex (DAG), PGBD1 might be associated with Duchenne muscular dystrophy (DMD), a common X-linked recessive neuromuscular disorder, in which patients exhibit significant cognitive and behavioural abnormalities^184^ [reviewed in^179^.

Furthermore, we present multiple lines of evidence, supporting a potential association of PGBD1 with SCZ susceptibility. In addition to its reduced level in SCZ patients, our analysis substantiates a previously identified exonic rs33932084 SNP^143^. Together with the already reported SNPs^56, 139, 185, 186^, rs33932084 strengthens the significance of the association between PGBD1 and SCZ susceptibility. Moreover, we have multiple common targets between SCZ susceptibility genes and PGBD1, several of them affected by aberrant splicing. An additional link between PGBD1 and SCZ pathology might be the oxidative stress marker protein, thioredoxin (TXN). Indeed, multiple studies show evidence that oxidative stress-induced damage in neurons largely effect the cognitive functions and this deficit is a hallmark of psychopathology of SCZ^187,188^.

## Supporting information

PGBD1_supplementary

## Acknowledgments

MDC Advanced Light Microscopy (ALM) technology platform Anca Margineanu, Matthias Richter, Anje Sporbert. MDC Flow cytometry technology platform Hans-Peter Rahn and Kirstin Rautenberg. We tank Sandra Neuendorf for technical assistance. L.D.H is funded by European Research Council Grant EvoGenMed ERC-2014-ADG 669207.Z.I. was funded by European Research Council, ERC Advanced ERC-2011-ADG 294742.

## Author contributions

The work was conceptualized by Z.I., T.R. and L.D.H. The original draft was written by Z.I. The manuscript was edited by L.D.H. Experiments were performed and the methodologies were worked out by T.R., A.Z., C.S., G.W., O.K, G.I, M.B. Bioinformatic analysis were performed by A.P, K.R, M.S. The paper was reviewed by T.R., A.P., K.R., A.S., S.P., M.S., T.I.O., A.P.

## MATERIALS and METHODS

### PGBD1 and SCAN domain evolution

To identify all SCAN family members, the human genome hg19 was downloaded from the UCSC genome browser^189, 190^ and translated with EMBOSS (http://emboss.sourceforge.net/)191 in all 6 reading frames. The SCAN domain m motif was extracted from the pfam database (http://pfam.xfam.org/)192 and the motif search was performed with the HMMER software (http://hmmer.org/)193 on all potential ORF (including alternative start codons). All SCAN domain hits with score higher than 25 were considered as significant. Each hit was determined to be coding using blastp against the UniprotKB/Swiss-Prot (The UniProt Consortium, 2017) database and the other hits were located with tblastn, both from the NCBI (https://blast.ncbi.nlm.nih.gov/Blast.cgi). The results were compared with the KEGG^194^, NCBI, Uniprot and BioMart^195^ databases. Proteins which match the domain alignment only partly (<58aa) are not shown (ZFP69B), because they probably do not contain this domain. All other domains were assigned using PFAM and SMART (http://smart.embl-heidelberg.de/)^196^ with the additional option for pfam domains. One should be aware that other motif search tools differ from this annotation. The Fig. (Suppl-Fig.X) shows only the longest transcript of each gene. To identify genomes which contain PGBD1 sequences we performed BLASTt and BLASTN searches (NCBI online platform) and found sequence similarities in almost all Eu-and Metatherian species, including *Phascolarctoscinereus*(koala), *Sarcophilus harrisii* (Tasmanian devil) and *Dasypusnovemcinctus* (armadillo) but not in phylogenetically older species. To validate whether these first transcripts encoded both PGBD1 associated protein domains, available RNA-seq data of these species were downloaded, and mapped against their reference genome, using STAR. Alignments of PGBD1 amino acid sequences were performed with MEGA7 (http://www.megasoftware.net/) using MUSCLE^197^ algorithm with default settings. PGBD1 amino acid sequences were retrieved from the NCBI database.

### Phylogenetic tree of PGBD1 and 2

All sequences(∼12k) containing the *pfam* domain Transposase IS4 have been downloaded from *interpro*Uniprot DB and aligned with *mafft*(default settings). An initial tree has been calculated with the UPGMA algorithm (default settings) from which a subtree has been manually picked. The subtree includes the cluster of PGBD1 & 2 plus some closely clustering sequences. Identical sequences (CD-HIT 100% identical) and sequences shorter than 250 bp have been removed. The PGBD1 & 2 sequences (XP_020822236.1 and XP_020822393.1) from Koala have been added manually. The picked transcripts were realigned using muscle (default settings) and a phylogeny tree was built using *MrBayes*(settings: mixed rate model, single chain and average standard deviation of split frequencies < 0.05). The tree was visualized with*iTOL*. Protein domains have been annotated with *hmmerscan* from the pfam db. For visualisation reasons another tree of representative PGBD1 and 2 plus invertebrate sequences has been built using *MrBayes (*settings: mixed rate model, single chain and average standard deviation of split frequencies < 0.05). Protein domains were annotated with hmmerscan and CDD (NCBI). The KRAB domain was annotated with Phyre2.

### K_A_/K_S_ ratios

Eleven mammalian sequences were picked to represent a heterogeneous group of species. A multiple sequence alignment was performed with MUSCLE for mammalian and primate specific analysis. From the mammalian tree the Ka/Ks ratios were calculated for different regions this indicating that PGBD1 mRNA (CDS) sequences from 11 mammalian and 18 primate species were manually selected and downloaded from NCBI. The 11 mammalian sequences were picked to represent a heterogeneous group of species. A multiple sequence alignment was performed with MUSCLE (for translated amino acids, default parameters) in UGENE for mammalian and primate specific analysis. The following taxonomy trees were used (it was manually modified to an unrooted tree) and retrieved from the NCBI: mammalian: (KOALA,(MOUSE,(PONAB,(HUMAN,PANTR),MACMU)),(HORSE,(PIG,PHYMC),CALUR,DESRO)); primates:

(PROCO,TARSY,(SAIBB,AOTNA,((PONAB,(HUMAN,(PATNR,PANPA),GORGO)),((9PRIM,COLAB,RHIBE),(CHLSB, MANLE,THEGE,(MACNE,MACMU,MACFA),CERAT))))); The K_A_/K_S_ ratios were calculated for different regions using PAML (M0) (overall, N-terminal (aa 1-290), C-terminal (aa 291-809), SCAN (aa 40-142), KRAB (aa 211-267), DBD1 (aa 405-541) and DBD2 (aa 750-804), reference is the human protein sequence of PGBD1).

### Neutral and adaptive evolution

Adaptive evolution of primates in the mammalian tree was tested with M1a vs. M2a (primates were foreground, all others background), with a chi2 test. Foregrounds (in PAML) are manually marked branches, which are tested against a background (unmarked). Adaptive evolution of the KRAB(-like) region in primates was tested with M1 vs. M2 with chi2 test.

### Dating Horizontal gene transfer

Ensembl synteny browser could not allocate a syntenic region between monotremes and human around the PGBD1 locus. Thus rather than synteny data we employ a recent method that aims to infer whether absence of a candidate gene in a given taxon is evidence for homology search failure or reflects true absence. AbSENSE^65^was run to test the possibility that PGBD1 was not detected in monotremes and reptiles due to failure of homology detection.

Evolutionary distances of 9 species pairs (human-rhesus macaque, human-Ma‘s night monkey, human-goat, human-camel, human-koala, human-platypus, human-American alligator, human-green anole and African clawed frog) have been calculated as described in the^65^: Orthologs have been retrieved from BUSCO curated vertebrate dataset. A total of 73 genes were common to all selected species. Isoforms have been selected according to their IsoSel score^198^. Sequences were aligned with MUSCLE (default) on a gene by gene basis and concatenated to one alignment. The evolutionary distances were calculated with protdist (PHYLIP, default). The focal species was human. Bitscores were calculated with blastp (NCBI). Significance testing was performed as proscribed^65^.

### PGBD1 conservation in rodent model organisms

The PGBD1 exon architecture and conservation track was retrieved from the UCSC genome browser (hg19). Multiple sequence alignment of mammalian PGBD1 sequences were used to detect the conservation of the catalytic domain (mouse: XP_030103153.1, rat: XP_017456282.1)

### Cloning of plasmid DNA constructs

*MER75 and piggyBac excision constructs*: pCAGGS-Venus was cleaved with PstI, treated with T4 DNA polymerase and Antarctic phosphatase and finally gel-isolated. This vector was ligated to blunt, 5’-phosphorilated PCR fragment of the moth piggyBac transposon from pUC19-XL-Neo (with TTAA on both ends). The fragment was amplified with the high fidelity Pfu Ultra II Fusion HS DNA polymerase (Agilent) according to the manufacturer’s recommendations using the primers: Piggy-forw (5’-P-TTAACCCTAGAAAGATAATCATATTGT-3’) and Piggy-rev (5’-P-TTAACCCTAGAAAGATAGTCTGCG-3’).

The *pTR-CAG-HA-PGBD1 expression plasmid* overproducing the PGBD1 protein utilized for MS-SILAC experiments was generated as follows: The PGBD1 ORF was amplified from a cDNA library prepared from HeLa cells using DNA polymerase Pfu Ultra II Fusion HS (Agilent Technologies) and PGBD1 Forw (5’-ATGTATGAAGCTTTGCCAGGC-3’) and PGBD1 Rev (5’-TTGCGGCCGCCTAATCTGACAG-ATGAGCATTGT-3’) primer pairs. A NotI restriction site was added to the 3’ end of the fragment for further cloning processes. PCR program was as follows: 95°C 150s pre-denaturation, then 35 cycles of 95°C for 30s, 63°C for 30s, 72°C for 60s, and finally 60s final extension at 72°C. A PCR product ∼ 2500 bp was digested with NotI, gel isolated (Qiaprep Gel Isolation Kit) and cloned into pHA5 expression vector (EcoRV and NotI), resulting in pTR-CAG-HA-PGBD1. In pTR-CAG-HA-PGBD1, the expression of PGBD1 ORF is driven by CAG, and the expressed protein is tagged with an N-terminal hemagglutinin (HA). To generate a non-tagged version (as a control construct for MS-SILAC), the DNA sequence encoding the HA-tag was eliminated by NotI/EcoRI digestion from pHA5. Before NotI digestion the EcoRI site was filled up using Klenow DNA polymerase. The PCR product encoding the PGBD1 ORF was inserted.

The *pTOV-T11-TR-HA-PGBD1, inducible expression construct* was generated by NcoI and NotI fragment replacement from pTR-CAG-HA-PGBD1 plasmid into previously cleaved pTOV-T11 plasmid^199^ by SalI and filled up using Klenow DNA polymerase.

For *knocking down PGBD1*, miR-expressing SB transposon vectors were generated by inserting the following elements into the pT2/HB transposon plasmid^200^: MPSV promoter and 5’ intron of the retroviral vector MP71^201^, mCherry as a marker gene followed by the posttranscriptional regulatory element (PRE) of woodchuck hepatitis virus, and the poly(A) signal (PAS) of psiCHECK2 (Promega, Mannheim, Germany). Redirected miRs targeting human PGBD1 were generated as described previously^202^ and introduced between PRE and PAS. In short, RNAi target sites were identified using BLOCK-iT RNAi Designer (Thermo Fisher Scientific) and redirected miRs were generated by overlap PCR using synthesized DNA oligos encoding the 21-nt antisense sequences and a plasmid encoding mouse miR-155^203^ as template.

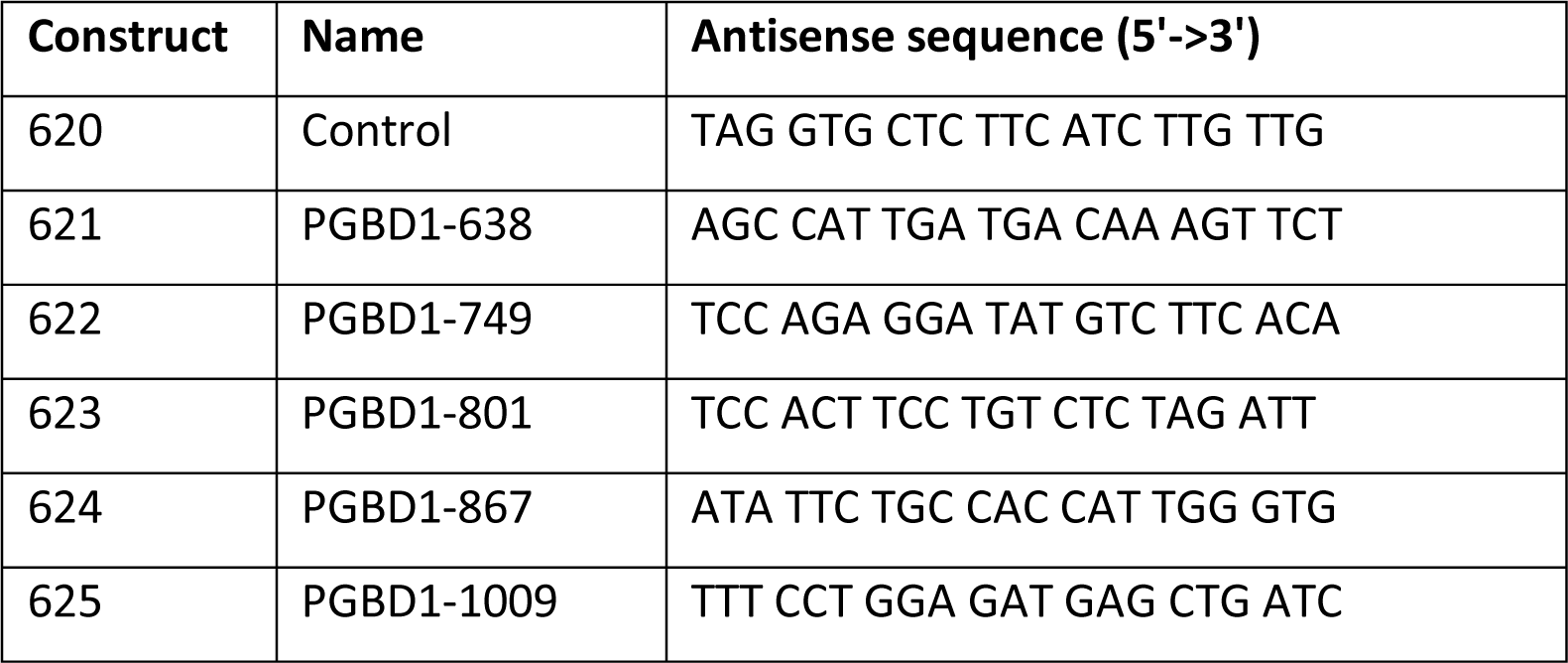

The plasmid constructs encoding *EGFP-tagged ATXN1 fusion protein of different length* of polyQ track (pT2-CMV/TetO_2_-EYFP-ATXNQ30 and Q82) were generated as follows: CMV promoter was excised from a pT2-CMV-EYFP-GW plasmid^204^ by HindIII and NheI digestion and replaced by a CMV/TetO_2_ promoter, generating a pT2-CMV/TetO2-EYFP-GW plasmid. ATXN1 Q30 or Q82 cDNA (GeneID: 6310) was shuttled into pT2-CMV/TetO2-EYFP-GW plasmid by LR recombination, as previously described^204^. Correct recombination was verified by BsrGI restriction digestion.

**Plasmid pEGFP-ATXN1-2Q** was from Prof.Dr. Huda Zoghbi’s lab contains the insert Ataxin-1 with Q2. (Addgene plasmid # 33239).

### Construction of CRISPR vectors

The pU6-sgRNA-CAGGS-Cas9-Venus-bPA (Addgene plasmid #86986) vector was cleaved by BbsI restriction endonuclease and ligated with the ds oligonucleotides encoding gRNA sequence. the following gRNA sequences were cloned:

Guide-79 sequence in the pU6-sgRNA79 plasmid: 5’-GGC GGC AAA TCT CCT GAG TG-3’

Guide-87 sequence in the pU6-sgRNA87 sequence plasmid: 5’-ATG ACA AAG TTC TCG GAG TT-3’

Guide-91 sequence in the pU6-sgRNA91 sequence plasmid: 5’-AGA TTT GCC GCC TGC GCT TT-3’

For **knocking out PGBD1**, a CRISP/Cas9 strategy was used. The pU6-sgRNA-CAGGS-Cas9-Venus-bPA (Addgene #86986) vector was cleaved using BbsI and ligated to the ds oligonucleotides encoding gRNA sequence. The following gRNA sequences were used: Guide-79 sequence in the pU6-sgRNA79 plasmid: 5’-GGC GGC AAA TCT CCT GAG TG-3’; Guide-87 sequence in the pU6-sgRNA87 sequence plasmid: 5’-ATG ACA AAG TTC TCG GAG TT-3’;Guide-91 sequence in the pU6-sgRNA91 sequence plasmid: 5’-AGA TTT GCC GCC TGC GCT TT-3’. The vectors were transfected into hESC_H9 and neural progenitor cells, respectively. 5 days post-transfection, the cells were FACS-sorted for the Venus fluorescence marker gene intensity.

For the***transposon excision assay*,** mPB is a plasmid vector expressing the mammalian codon optimized version of the insect piggyBac transposase, kindly provided by Allan Bradley (Wellcome Trust Sanger Institute, UK) (*repository number: pCyL43;*^205^*PB-PGK-Puro vector*: The pCyL50 PB cloning vector (*from Allan Bradley, Wellcome Trust Sanger Institute, UK;*^205^was used to insert the human phosphoglycerate-kinase (PGK) promoter driven puromycin resistance gene unit between the PB inverted repeat sequences. The transcription unit was PCR-amplified and ligated into the PB transposon cloning vector. *PGBD1 expression vector (pSB-cmv-GFP-cag-HUCEP4):*The coding sequence of the human PGBD1 fusion gene was PCR-amplified from cDNA prepared from HEK-293 total RNA. The primer sequences: PGBD1-forward: 5’-g**<u>gctagc</u>***cgccacc*atgtatgaagctttgccaggccctg-3’ (**<u>NheI</u>**-*Kozak sequence*-PGBD1 coding); PGBD1-reverse: 5’-tgc**<u>agatct</u>**ctaatctgacagatgagcattgtg -3’ (**<u>Bgl II</u>**-PGBD1 coding). After NheI /Bgl II digestion, the PCR product was ligated into an expression vector after a CAG promoter. The vector contains a separate CMV-GFP expression cassette for monitoring transfection efficiency. In addition, it also has the *Sleeping Beauty* (SB) inverted terminal repeat sequences in appropriate positions, in order to allow for establishing stable cell lines using the SB100X transposon system.

### MER75B-puro / MER85-Puro vectors

To select potentially the best MER sequences, the most important criteria were the high similarity to the consensus sequences and the presence of intact inverted terminal repeats (ITRs) at both 5’ and 3’ ends^46^. The selected human MERs with their short flanking genomic sequences were PCR-amplified from HEK293 gDNA preparations, using the following primers: MER75B-cloning-For: 5’-GGATCCTTTCCTTCACCCTCCCTGC -3’ MER75B-cloning-Rev: 5’-GGATCCAGTCACCCCAAGGAGAAAAGG -3’*(from human chromosome 4, Sequence ID: NC_000004.12, nt: 143088382 to 143088750*); MER85-cloning-For: 5’- GGATCCTTAACACATCAGACATGGAGGG -3’; MER85-cloning-Rev: 5’-GGATCCGGCCAAAGCATATGTTCTTAATC -3’*(from human chromosome 21, Sequence ID: NC_000021.9, nt: 34685694 to 34686125)*

The Sanger sequencing verified MERs were cloned into the pGEM®-T Vector Systems, then BsmAI and HindIII restriction sites were used to insert a PGK-Puro-polyA expression cassette into the MER75B and MER85 sequences, respectively. In the formed expression constructs, the ITRs at both ends of the MER sequences remained intact, allowing the potentially active transposase excising the expression cassette as a transposon. Note that MER75 is a predicted substrate of PGBD4, whereas MER85 is a predicted substrate of PGBD3.

### Excision repair reporter assay

Human HeLa cells were seeded in 12-well cell culture plates (100,000 cells/well). Transfections were performed using plasmids encoding HA-tagged PGBD expression construct (500 ng each) together with 500ng pEXC-mPB plasmid (in 1:1 ratio). 48 hours post-transfection cells were washed three times by DPBS and treated by trypsin for 2 min. After treatment 1 ml DPBS supplemented by 10% fetal serum was added to the cells and filtered through FALCON 5 ml Polystyrene round-bottom tube with cell-strainer cap. 60,000 cells GFP intensity was then measure by FACS CaliburTM System.

### Cell lines and transfection conditions for transposon excision assay

Human embryonic kidney cells (HEK293) cells were cultured in Dulbecco’s modified Eagle’s medium supplemented with 10% of fetal calf serum, 1% of L-glutamine, and 1% of penicillin–streptomycin (Thermo Fisher Scientific). Cell were transfected using the FuGENE 6 reagent (Bio-Science Ltd., Hungary), according to the manufacturer’s instruction. Briefly, 4 x 10^5^ cells were seeded onto 6-well plates and 1 day later, cell were transfected with the lipid-DNA mixture as specified by the protocol, using 500 ng of total DNA, composed of 250 ng transposon containing plasmid mixed with 250 ng of a transposase/putative transposase/control expression vector. Transfection efficiency was checked by fluorescence microscopy detecting GFP signal where appropriate, and quantified by FACS measurements using as FACSCanto instrument (BD Biosciences).

### Transposon excision assay

To detect transposon excision events from the donor plasmids, plasmid DNA was isolated from the transfected cells 48 hours post-transfection, using standard phenol/chloroform extraction method, followed by ethanol precipitation. To detect those plasmids that were re-circularized following transposon excision, followed by cellular DNA repair, the extracted plasmids were subjected to a nested PCR in 2 rounds. Primers used for the assay are as follows: Excision, first round, forward: 5’-GCGAAAGGGGGATGTGCTGCAAGG -3’; Excision, first round, reverse: 5’- TCTTTCCTGCGTTATCCCCTGATTC-3’; Excision, second round, forward: 5’- CGATTAAGTTGGGTAACGCCAGGG -3’; Excision, second round, reverse: 5’- CAGCTGGCACGACAGGTTTCCCG -3’. For normalization, PCR on the plasmid backbone (on the ampicillin resistance gene sequence) was carried out, resulting in a product, regardless of transposon excision. Excision, plasmid control, Amp64-Forward: 5’-TTTGCTCACCCAGAAACGC -3’; Excision, plasmid control, Amp403-Reverse: 5’-AGTTGGCCGCAGTGTTATCAC -3’

### Colony forming assay to detect stable integration

To detect stable integration after transposition, puromycin resistance gene expressing transposons were used for antibiotic selection experiments^206^.After 48 hours post-transfection, 1% of transfected cells were seeded onto cell culture Petri dishes, selected for 2-3 weeks with 1 µg/ml of puromycin (Sigma-Aldrich), and surviving cells were fixed with methanol and stained with Giemsa (Sigma-Aldrich). Colonies were quantified in a 75S model gel imager, using the Quantity One 4.4.0 software (Bio-Rad).

### Generation of stable PGBD1 knockdown and overexpression SHEP neuroblastoma cell lines

Note that to deplete PGBD1 in hESCs, first, we used a CRISPR/Cas9-mediated knocking out (KO) approach. However, using the KO strategy, no stable, proliferative KO line could be generated (Extended Data Fig. 6a). Similar, this KO approach has also failed in neural progenitor cells (NPC), suggesting an essential function of PGBD1 in cell survival. As an alternative, we used the knocking down (KD) RNA depletion strategy. to KD PGBD1, 300,000 human SHEP neuroblastoma cells were co-transfected using different pmiRNA constructs (marked by mCherry), pTOV-T11-TR-HA-PGBD1 together with pSB100X^207^ plasmid encoding the SB100X transposase in ratio 10:1. 3 days post-transfection, the cells were FACS-sorted for mCherry marker gene intensity. The sorted single cells were cultured for three weeks then subjected to an additional FACS sorting.

To generate overproducing HA-PGBD1 cell lines, SHEP cells were transfected using pTOV-T11-TR-HA-PGBD1 inducible expression construct together with pSB100X plasmid encoding the SB100X transposase in ratio 10:1. The transfected cells were cultured for four weeks in Dubelcco’s modified Eagle’s medium (DMEM) with 4.5 g/l D-glucose containing 10% fetal bovine serum, supplemented with G418 (gentamicin) in final concentration 500 µg/ml. PGBD1 protein level was determined using Western-blotting. To induce HA-PGBD1 protein expression, the media was supplemented by 1µg/ml doxycycline in final concentration.

### Cell culture and cell transfection for additional cell lines

Human cervical carcinoma cells (HeLa) and human neuroblastoma (SHEP) cells were cultured in Dubelcco’s modified Eagle’s medium (DMEM) with 4.5 g/l D-glucose containing 10% fetal bovine serum, supplemented with penicillin (100 μg/ml) and streptomycin (100 µg/ml) at 37°C and 5% CO_2_. In general, the cells were seeded in 10 cm or 6-well plates and transfection was performed using JetPrime^TM^ reagent following the recommended manufacturer’s protocol.

### Stable isotope labelling by amino acids in cell culture (SILAC)

Two populations of HEK293 cells were cultivated in cell culture for three weeks.One population of cells was fed with growth medium containing normal amino acids (“light cell population”). The second population of cells was cultured in growth medium, containing amino acids labelled with stable heavy isotopes (^13^C_6_-^15^N_4_ L-arginine; ^13^C_6_-^15^N_2_ L-lysine) (“heavy cell population”). In our experimental approach, untagged PGBD1 and HA-tagged PGBD1 were overexpressed in both conditions. The overexpressing plasmids encoding the HA-tagged or untagged PGBD1 were transfected into the two cell populations of HEK293 cells, respectively. 3 days post transfection protein purification was performed using EZview^TM^ red coloured Anti-HA agarose affinity gel, following the recommendations of the manufacturer. Purified protein mixtures were prepared as follows: (a) as forward experiment: protein mixture of Heavy HA-PGBD1 and protein mixture of Light PGBD1; (b) as reverse experiment: protein mixture of Light HA- PGBD1 and protein mixture of Heavy PGBD1. The purified protein mixtures including HA-tagged target proteins and their interacting partners were subjected to LS-MS/MS (mass spectrometry): Samples were processed by methanol-chloroform extraction, reduced, alkylated and digested with LysC and trypsin using standard protocols. After offline desalting, peptides were analysed by LC-MS/MS on a Proxeon EASY-nLC II system, connected to a Q Excative mass spectrometer (Thermo Scientific). Chromatography was performed using a 120 min acetonitrile gradient on a 25cm long inhouse prepared column (ReproSil-Pur 120 C18-AQ, 3 µm (Dr.Maisch GmbH HPLC)). The instrument was operated in the data dependent mode with the following settings for the full scans: resolution 70,000, AGC target value 3E6, maximum injection time 20 ms. The following settings were chosen for the MS2 scans: resolution 17,500, AGC target value 1E6, maximum injection time 60 ms. Raw files were processed with MaxQuant (version 1.4.1.2).

### SILAC-AP-MS Analysis

#### Calculation of significance

The significance was calculated as described in^208^ with small modifications. First, we filtered out contaminants and proteins for which no unique peptide was detected. We performed logarithmic transformation (log2 with pseudo count 1) of the normalized H/L ratios and confirmed a Gaussian distribution of the transformed data (Shapiro-Wilk test). The 15.87^th^, 50^th^ and 84.15^th^ percentiles were calculated and called r_-1_, r_0_ and r_1_, respectively. The z-transformation is defined as: 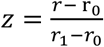 for r > r0 and 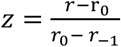 for r < r_0_, where r are the transformed H/L ratios.

The p-values were calculated with significance A formula. In the original paper, significance A gives two-sided p-values. Here we considered the sidedness of the test and adjusted one side of the distribution (one-sided):

significance 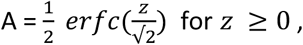

and significance 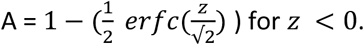

The sidedness of the label-switch experiment is reverse. Which means that *z* ≥ 0 was adjusted for sidedness. The p-values were adjusted for multiple testing with the Benjamini-Hochberg method. We only considered proteins as significant if they were also significant outliers in the label switch experiment (intersection). P-values of both experiments were combined with the berger method of the scran R package^209^. The script was written in Rstudio (R version 3.6.3), using erfc function from the pracma package.

#### Gene set enrichment analysis

All gene sets of the Molecular Signature Database v7.2^210, 211^ were downloaded. Terms with at least 5 gene members and less than 500 were tested for enrichment. The enrichment was tested with an overrepresentation test approach, the „HG“ algorithm from the tmod R package^212^.

#### Immunoflourescence microscopy

The cells were seeded on coverslips in 12-well cell culture plates (100,000 cells/well). 48 h after transfection, cells were fixed with 4% paraformaldehyde (Sigma) supplemented with Hoechst 33,342 (1:1,250, Invitrogen) in PBS for 15 min, and permeabilized with 0.1% Triton X-100 in PBS for 2 min. Coverslips were incubated with primary antibodies for overnight at 4°C, then washed three times with PBS, followed by an incubation using secondary antibodies for 60 min. After an additional washing step, the samples were mounted using ProLong® Gold antifade reagent (Invitrogen). The images were taken using a Leica LSM710 point-scanning single photon confocal microscope.

#### RNA-FISH and immunofluorescence

Cells grown on coverslips were fixed using 4% paraformaldehyde and permeabilized with 70% ethanol overnight. For RNA-FISH, Stellaris RNA-FISH probes labelled with Quasar 570 Dye for NEAT1_2 (SMF-2037-1) (1:100, Biosearch Technologies) were used according to the instructions provided. For subsequent immunofluorescence staining, SFPQ antibody (1:60, WH0006421M2, Sigma) and PGBD1 antibody (1:200, Abcam, ab180598) were used. Finally, cells were counterstained with DAPI (4ʹ,6-diamidino-2-phenylindole) in water for 15 min at room temperature. After an additional washing steps, the samples were mounted using ProLong® Gold antifade reagent (Invitrogen). The images were taken using a Leica LSM710 point-scanning single photon confocal microscope. Paraspeckles were defined as NEAT1_2 RNA-FISH signals that are colocalizing with SFPQ.

#### Co-immunoprecipitation

500.000 non-transfected or transfected (using plasmids encoding EGFP-ATXN1 with different polyQ tracks) human neuroblastoma SHEP cells were lysed in 200 µl lysis buffer containing 50 mM TRIS HCl pH 8.0, 10 mM EDTA, 100 mM NaCl, 5% glycerol, 1% NP-40 supplemented with COMPLETE protease inhibitor cocktail (Roche) and Benzonase®Nuclease (Sigma) for 40 mins at 4°C. After incubation, the lysates were centrifuged for 10 min at 12,000 rpm at 4°C to remove unbroken cells and cell debris. Supernatant was collected and protein concentration was determined. In parallel, 60 µl Dynabeads^TM^ ProteinG was washed as recommended by the manufacturer, and resuspended in 200 µl PBS+0,02 % Tween20. The mixture was incubated with 5 µg of primary antibodies for 1 h at room temperature on a turning wheel. After incubation, the magnebeads were collected and washed with 200 µl PBS+0,02 % Tween20, then 150-200 µg of cell lysates was added to the beads. Cell lysates together with the magnebeads were incubated for 3 hours at 4°C on a turning wheel. The beads were washed three times with 200 µl PBS+0,02 % Tween20. The proteins were eluted from the beads by adding a mixture of 20 µl 100 mM glycine pH 2,8 and 10 µl of 5 x SDS-loading dye (10% SDS, 10 mM DTT, 20 % glycerol, 0.2 M Tris-HCl pH 6.8, 0.05% Bromophenolblue) and boiling at 70°C for 5 min.

#### Western blotting

In general, the cells were collected from 6-well plates for Western blot analysis. The cells were subsequently washed with phosphate-buffered saline (PBS), and lysed on ice for 40 min in lysis buffer containing 50 mM TRIS HCl pH 8.0, 10 mM EDTA, 100 mMNaCl, 5% glycerol, 1% NP-40 and Protease Inhibitor Mini Tablets, EDTA Free (Pierce™) and Benzonase Nuclease (NOVAGEN) as recommended by the manufacturer. Total lysates were resolved by 8-12 or 4-20 % Mini-Protean® TGX™ SDS-PAGE Gel, and transferred to PVDF membranes by Bio-Rad Trans-Blot®Turbo™ Transfer System. The membranes were blocked with TBS-T (TBS supplemented with 0.05% Tween-20) containing 5% non-fat dry milk and were incubated overnight at 4°C with primary antibodies in appropriate dilutions in TBS-T containing 5% non-fat dry milk. Membranes were washed with TBS-T buffer, incubated for 1 hour at room temperature with Alkaline Phosphatase-conjugated (Sigma-Aldrich) or Horseradish Peroxidase-Conjugated (Promega) secondary antibodies in TBS-T containing 5% non-fat dry milk. The blots were subsequently washed with TBS-T. Bands were detected by ECL^TM^ Prime Western Blotting Detection Reagent (Amersham) and images were then analysed by ChemiDoc™MP Imaging System (Bio-Rad).

#### Antibodies and chemicals

Anti GFP polyclonal antibody (1:1000, Abcam, ab290) was used to monitor EGFP-tagged recombinant fusion ataxin1 proteins. Anti-PGBD1 antibody (1:1000, Abcam, ab180598), anti-TRX antibody (1:1000; abcam, ab133524), anti-HA monoclonal antibody (1:2000, Roche), anti-Ataxin 1 (phospho S776) antibody (1:1000, abcam, ab63376), Anti-Hsp70 antibody (1:1000, Abcam, ab2787), monoclonal anti-*α*-actinin (1:1000, Sigma, A7811), mouse anti-ZNF24 antibody (1:1000, Abnova, H00007572-M02), Anti-tubulin (1:1000, Millipore, MAB1637), aSFPQ (WH0006421M2, Sigma). Green fluorescence goat anti-rabbit antibody Alexa Flour® 488 (1:200, Invitrogen), green fluorescence donkey anti-rabbit antibody Alexa Flour® 488 (1:200, Invitrogen), red fluorescence goat anti-rat antibody Alexa Flour® 568 (1:200, Life Technology), red fluorescence donkey anti-mouse antibody Alexa Flour® 555 (1:200, Invitrogen), goat anti-rabbit IgG (1:5000, Thermo scientific, 31462), goat anti-mouse IgG (1:5000, Thermo scientific, 31432), goat anti-rat IgG (1:5000, Thermo scientific, 31470), red fluorescence donkey anti-mouse antibody Alexa Flour® 647 (1:200, Life Technology).

#### Filter retardation assays

Quantitative measurement of the polyQ-containing ataxin1 protein aggregates was performed by using a filter retardation assay^213^. Stable human neuroblastoma SHEP cells (300. 000) either overproducing or knocked down PGBD1 were transfected (as described previously) with 2 µg plasmids encoding EGFP-tagged ATXN1 with different polyQ tracks. 48 hours post-transfection, the cells were lysed on ice for 40 min in lysis buffer containing 50 mM TRIS HCl pH 8.0, 10 mM EDTA, 100 mMNaCl, 5% glycerol, 1% NP-40 and COMPLETE protease inhibitor cocktail (Roche). Equal amounts of crude cell extracts (100 µg total protein) were mixed and diluted in 150 µl DPBS supplemented with 0.1% SDS and filtered through a 0.2 µm cellulose acetate membrane using a 96 well dot-blot microfiltration apparatus. The membrane was washed twice with 100 µl DPBS with 0,1% SDS. ATXN1 aggregates retained on the membrane were detected using the polyclonal anti-GFP antibody (1:1,000).

#### Measuring ROS

The cells are fed with dichloro-dihydro-fluorescein diacetate (DCFH-DA) and, following an intensive washing step, the fluorescent signal is measured by FACS. Stable human neuroblastoma SHEP cells (100.000) either overproducing or knocked down PGBD1 were seeded on 12 well plates. After 24 hours, the cells were fed by dichloro-dihydro-fluorescein diacetate (DCFH-DA)(D6883, Sigma-Aldrich) for 60 min in fresh DMEM media. After removal of the media, the cells were gently washed three times with Dulbecco’s phosphate-buffered saline (DPBS), and resuspended in DPBS, supplemented by 10% fatal serum albumin. The intensity of the green fluorescence signal was quantified using a FACSCalibur^TM^ System (Becton-Dickinson), measuring 10.000 individual cells per sample. To determine the PGBD1 protein level after oxidative stress, 100.000 SHEP cells were seeded on 12 well plates, and treated with 2 mM H2O2 for 20 minutes in OPTI-MEM media. After treatment, the cells were washed with DPBS, and incubated in DMEM/COMPLETE media for the recovery intervals (4, 8, 24 hours). At the end of the recovery timepoints, the cells were collected and the total cell extract was analysed by Western blotting.

#### ChIP-exonuclease (exo) assay

The ChiP-exonuclease assay protocol was performed as in Serandour’s method^214^. The libraries were quantified by using the KAPA library quantification kit for Illumina sequencing platforms (KAPA Biosystems, KK4824) and sequenced on HiSeq following the manufacturer’s protocol.

#### ChIP-seq for histone tail modifications - peak calling

MACSv2 (Model-based Analysis for ChIP-seq)^215^ was utilized for the detection/analysis of genome-wide broad peaks representing histone tail modifications. Replicates were pooled separately for each histone modification. Peaks were called with input/mock DNA samples for identification of unspecific signals. Candidate peaks were selected according to the threshold values: q-value <= 0.01 and mfold = 10,100 (default 5, 10). The mfold parameter selects only those regions that are mfold or higher enriched for ChIP-seq reads compared to a random genome-wide distribution (fold enrichment for the peak summit against random Poisson distribution computed with the local lambda). Consensus peaks between biological replicates were calculated with DiffBind (v 2.10.0). Histone modification data for NPC and neurons were obtained from GSE119006 and GSE62193. Active promoters were defined as (H3K4me3+H3K27ac+H3K36me3), repressed promoters as (H3K4me3+H3K27me3-H3k27ac-H3K36me3) and poised enhancers as (H3K4me1+H3K27me3). The overlaps were calculated with BedToolsintersectBed command with –f 0.5 –r.

#### IRAlus identification in 3’UTRs

Our list for the 42 identified lRAlus had three levels of filtering: (i) PGBD1 binding at 3’UTR; (ii) within the Alu insert remove FLAM and FRAM (right and left arm) coordinates; (iii) It should be in the classical orientation favouring the formation of IRAlus.

### RNA-sequencing data analysis

#### Raw reads filtering

Software tools such as the FASTX-Toolkit and Trimmomatic were used to discard low-quality reads, trim adaptor sequences, and eliminate poor-quality bases. Outliers with over 30 % disagreement were discarded.

#### Read alignment

Salmon was used to build index and align the reads using the following commands: salmon index -t transcripts.fa -itranscripts_index --decoys decoys.txt -k 31, ./bin/salmon quant -itranscripts_index/ -l IU - 1 fastq -2 fastq --validateMappings –o output. Quantified data was checked for GC content and gene length biases using R package NOISeq to provide useful plots for quality control of count data.

#### Reproducibility

Mean variance and PCA were computed between biological replicates using the tximport package^216^ in R using lengthscaledTPM (CPM cutoff>2 and sample cutoff 2 between the replicates) for the analyzed groups (NPC-KD, KD-Neuron). Batch effects were removed using the RUv package of Bioconductor^217^. Subsequently, the samples were normalized using TMM (weighted trimmed mean of M-values) method for differential expression analysis. The differential expression analysis was conducted using the DESeq2 package. Dataset for the analysis of schizophrenia and control was obtained from GSE145656 and analyzed as above.

#### ChIP-exo data analysis for PGBD1 peak calling

To map the sequencing reads, the FASTQ files were aligned to the human genome hg19 using Bowtie2. To filter out PCR duplicates, we used Samtools in the following steps: (i) filter out low-quality (<20) reads; (ii) sort reads by name (sort -n); (iii) fix read pairs (fixmate); (iv) sort reads by chromosomal position (sort); (v) mark duplicates (markdup); (vi) extract only read_1 from each non-duplicated pair (view -f 0x40).As the majority of TFs bind as dimers (either homo or hetero), it was necessary to determine the optimal trim length. Therefore, we took 3 times the radius of the PGBD1 and converted that size into number of bps. After rounding, this resulted in the optimal trim length in bp.BEDTools “genome coverage” function was employed to generate the read profiles for both strands separately using the determined optimal trim length. Subsequently, both strands were combined and only base positions where there are reads on both strands were reported as the PGBD1 binding profile. After this step, the replicates were normalized based on their average background read count and then combined using their average read count per base position. GEMv3.4 was used to detect narrow peaks using the following command: --d Read_Distribution_ChIP-exo.txt --expt PGBD1.bam --ctrl PGBD1.bam --g hg19.chrom.sizes --f BAM -- genome hg/ --k_min 6 --k_max 18 --outBED --outNP --smooth 3 --mrc 20 –q 3

#### Transposable element identification and enrichment in PGBD1 peaks

The overlap of PGBD1 peaks with Repetitive elements (RepBase annotation) was performed with BedToolsintersectBed command (-f 0.1 –r). The enrichment of a particular TE class was calculated as following- 200 bp DNA-wide window was extracted from the center of the PGBD1 peak using Bedtools slop –b –l 200 –r 200 command^218^.Dataset of 10000 randomly chosen peaks, containing the same number of regions with the same length (200 bps) and the same nucleotide distribution as true regions was generated using the BedTools random command. To do this, each chromosome was divided in genomic windows of 1,000,000 bps and, for every real peak, the corresponding random peak was taken with flat distribution from the same window. All repeats falling in the peaks belonging to the real dataset and in the 10000 datasets of random peaks were annotated. This method delivered, for each transposable element, mean and variance, which was used to calculate a z-score zr = (xr − μr) / √sr, where xr is the occurrence of a particular transposable element r in the original dataset, while μr and sr are respectively its mean occurrence and its variance in the 10000 random sets. Similar analysis was performed with classes of transposable elements and the z-score zc = (xc − μc) / √sc, where xc is the occurrence of a particular class of transposable elements in the original dataset, while μc and sc are respectively its mean occurrence and its variance in the 10000 random sets.

#### ATAC-seq data analysis in PGBD1 peak regions genome-wide

Genrich tool was utilized for detection of peaks in the PGBD1 binding sites. Genrich calls peaks for multiple replicates collectively. First, it analyzes the replicates separately, with p-values calculated for each. At each genomic position, the multiple replicates’ p-values are then combined by Fisher’s method. The combined p-values are converted to q-values, and peaks are called.

FASTQ data were processed for alignment parsing, removal of multimapping reads, PCR duplicate removal, Genome length calculation, Control/background pileup calculation and p-value calculation using the following command: ./Genrich -t ATACseq.bam -o ATACseq.narrowPeak -f ATACseq.log -r -x -q 0.05 -a 20.0 -v -e chrM,chrY -E hg19_Ns.bed,wgEncodeDukeMapabilityRegionsExcludable.bed.gz ./Genrich -P -f ATACseq.log -o peaks.narrowPeak -p 0.01 -a 200 -v Peak-calling from log file: ATACseq.log.

#### GRO-Seq data analysis

GRO-seq data (sra format, GSE140486) were processed into the FASTQ format with the ‘fastqdump’ command (SRA toolkit)^219^. The resulting cDNAs were trimmed with Homer v 4.10 to remove 3ʹ terminal A-stretches, which had been attached during library construction (homerTools trim)^220^. Only cDNAs ≥25bp entered the analysis. Datasets were quality filtered with the FASTX (v 0.0.13) software tool (-q 10 - p 97) (http://hannonlab.cshl.edu/fastx_toolkit/), and resulting GRO-seq cDNAs were aligned to the human genome assembly (hg19) using Bowtie version 0.12.9 (-v 2 -k 3 -m 1 –best). BAM files were utilized to calculate GRO-seq peaks using the annotatePeaks function from HOMER (v 4.10). Protein coding genes were divided into 2 groups based on the number of PGBD1 peaks occurring in the entire gene body (1-2 vs. 3-7) Pausing using index for RNAPII in the dataset was calculated S as follows: S= log2 (d (RNAPII_PGBD1bindingsites_)) – log2 (d(RNAPII_gene body_)). Ratio of RNAPII read density at the PGBD1 binding sites within protein coding genes to that of the RNAPII read density in the gene body. d stands for the number of reads per nucleotide (nt) in the given region. The difference between the densities in log2 units equals to the ratio of fold enrichment in these regions, meaning a value of 1 would represent a 2-fold greater enrichment of RNAPII signal at the promoter region rather than in the gene body^221^.

#### Electrophoretic mobility shift assay (EMSA)

Approximately 11x10^6^ HEK293 cells were transfected with 12 μg pT2-CAG-HA-PGBD1 plasmids encoding HA-PGBD1 fusion protein. One day post-transfection cells were collected and washed with DPBS. Cells were lysed in 500 μl lysis buffer (50mMTris-HCl, pH8.0, 100mM NaCl, 10m MEDTA, 5% glycerine, 1% NP- 40 and protease inhibitor cocktail (Roche)) for 40 min at 4°C. Following removal of the cell debris by centrifugation at 20,000 g, HA-PGBD1 protein was purified by using EZview™ Red Anti-HA Agarose beads. The HA-tagged PGBD1 was eluted by 500 μl 100 μg/ml HA-peptide (Sigma) in 100 mMNaCl. The protein mixture was concentrated by Amicon Ultra-4, Ultracel -3K filter column. Centrifugation was performed at 7500 rpm for 40 min at 4°C. The protein mixture concentration was measured by standard BCA method and the level purity was analysed by loading 5 μl onto 10% SDS-PAGE.Binding reactions were performed in 25 μl volumes on ice for 20 min. DNA binding reactions contained FAM-labelled PGBD1-specific (GS-PGBD1-UP:5’-GCTTTC<u>AATGGAATGGAATG</u>CCTTCC-3’), complementary (GS-PGBD1-LOW:3’-CGAAAG<u>TTACCTTACCTTAG</u>GGAAGG-5’) dsDNA oligonucleotides (PGBD1 oligo), purified HA-PGBD1 protein, 10 mM Tris-HCl pH 8.5, poly(dI-dC), 1 mM EDTA, 50 mM KCl, 10 mM 2-mercaptoethanol. As competitor dsDNA oligonucleotide we used the PGBD1-specific oligonucleotide without FAM labelling. The gel buffer contained 25 mM Tris-borate pH8.3, 1 mM EDTA. Protein–DNA complexes were separated by electrophoresis in 6% non-denaturing polyacrylamide gels at 4°C. Electrophoresis was performed at constant voltage of 200V for 3 h. The fluorescent signal was detected by using a BioRadChemiDoc^TM^ MP Imaging System.

#### Generation of NPCs and neurons from hESCs

We used the human embryonic stem cell (hESC) line H1 according to the German law under a license approved by the Robert Koch Institute (license to A. Prigione # AZ: 3.04.02/0077-E01). The derivation of NPCs was the same as reported^78^. We performed the generation of midbrain dopaminergic neurons (mDAN) following a previously published protocol^222^. We first let the NPCs to differentiate onto Matrigel coated plates for 8 days using a medium containing: Neurobasal: DMEM/F12 (1:1), N2 (1x), B27 (1x), purmorphamine (1 µM), vitamin C (200 µM), and FGF8 (100 ng/ml). Afterwards, we cultured the cells for two additional days using a medium containing: Neurobasal: DMEM/F12 (1:1), N2 (1x), B27 (1x), purmorphamine (500 nM), and vitamin C (200 µM). We next split the cells using Accutase at 1:3 ratios and plated them onto matrigel-coated dishes. Finally, we switched the culture conditions to the maturation medium containing: Neurobasal/DMEM-F12 (1:1), N2 (1x), B27 (1x), vitamin C (200 µM), db-cAMP (500 µM), BDNF (10 ng/ml), GDNF (10 ng/ml), and TGFbeta3 (1 ng/ml). We kept the differentiating neurons in the maturation medium for eight weeks with medium changed every other day.We kept NPCs and neuronal cultures in a humidified atmosphere of 5% CO2 at 37°C under atmospheric oxygen condition and regularly monitored them against mycoplasma contamination.

#### Generation of stable PGBD1 knock-down NPC cell lines and samples preparation for RNA-sequencing

To deplete PGBD1, we first used a CRISPR/Cas9-mediated KO approach, however, no stable, proliferative KO line could be generated in either hESCs or neuroprogenitor cells (NPC) (Extended Data Fig. 6a), suggesting an essential function of PGBD1 in cell survival. As an alternative strategy, we used knock down (KD) RNA depletion. The established KD-mi621-NPC line was generated by transfecting a combination of miRNA KD constructs. This approach resulted in a ∼ 65% depletion of PGBD1 in NPCs on protein level (Extended Data Fig. 6b).

#### detailed protocol

300,000 human NPCs were transfected by different pmiRNA constructs together with pSB100 plasmid encoding SB100 transposase in ratio 10:1. 3 days post-transfection the pmiRNA contained cells were FACS sorted based on mCherry marker gene intensity. The sorted single cells were cultured for additional three weeks then cells were FACS sorted repeatedly. To increase the knock-down efficiency 6*10^5^ smNPC-miRNA621 (p23) stable cells (in 6 wells plate) were additionally transfected by 3 μg (1-1-1 μg) pmiRNA623,624,625 plasmids. 3 days post-transfection in 1 ml STEMdiff Neural Progenitor Medium (Stemcell Technologies) cells were washed by DPBS and 2 wells of the plate were put together and collected by centrifugation at 10.000 rpm for 2 min generating 3 biological replicates. 1/4 part of the collected cells (per samples) was tested by Western blot to determine the PGBD1 level. Total RNA was extracted from cells by using the Trizol kit (Invitrogen) following the manufacturer’s instructions. The sequencing libraries were prepared by using TruSeq RNA Indexes Set A, (Illumina 20020492) and TruSeq®Stranded mRNA LT kit 48 Samples (Illumina, 20020594), according to manufacturer’s instructions. High throughput single-indexed, paired-end 150 bp sequencing was performed in a HiSeq4000 instrument (Illumina, USA).

### qRT–PCR validation of selected DEGs in PGBD1-depleted NPC

Total RNA was extracted from cells by using the Direct-zol™RNA MiniPrep kit following the manufacturer’s instructions. 1 μg purified DNaseI-treated RNA was used for reverse transcription (RT) by High Capacity RNA-to-cDNA kit (Applied Biosystems). 1 μg purified DNaseI-treated RNA was used for reverse transcription (RT) (High Capacity RNA-to-cDNA kit, Applied Biosystems). Quantitative RT–PCR (qRT–PCR) was performed using the Power SYBR Green PCR Master Mix (Applied Biosystems) on the ABI7900HTsequence detector (Applied Biosystems). Data were normalized to GAPDH expression using the ΔΔCt method. Error bars represent the standard deviation (s.d.) of samples carried out in triplicates.

Sequence of the primers used in qPCR: NEAT1-qFwd: 5’-TTGTTCCAGAGCCCATGAT-3’

NEAT1-qRev: 5’-TGAAAACCTTTACCCCAGGA-3’

NEAT1_2-qFwd: 5’-GATCTTTTCCACCCCAAGAGTACATAA-3’

NEAT1_2-qRev: 5’-CTCACACAAACACAGATTCCACAAC-3’

MSI1-forward2: 5’-GTCACTTTCATGGACCAGGC-3’

MSI1-reverse2: 5’-CCGGTTGGTGGTTTTGTCAA-3’

NEUROD1-forward1: 5’-GAGACGCATGAAGGCTAACG-3’

NEUROD1-reverse1: 5’-CTGAACGAAGGAGACCAGGT-3’

PAX3-forward2: 5’-GGCATGTTCAGCTGGGAAAT-3’

PAX3-reverse2: 5’-TGCTGTGTTTGGCCTTCTTC-3’

PAX6-forward1: 5’-ACCGGTTTCCTCCTTCACAT-3’

PAX6-reverse1: 5’-GGAGTATGAGGAGGTCTGGC-3’

GAP43-forward2: 5’-CTCATAAGGCCGCAACCAAA-3’

GAP43-reverse2: 5’-GGTGCCTTCTCCCTTCTTCT-3’

ZNF24-forward1: 5ʹ-ATGTCTGCACAGTCAGTGGAAGAAGATTCA-3’

ZNF24-reverse1: 5ʹ-CGGAAAATCTCTGGGTCTGGGAGATGGTTC-3’

GAPDH-forward1: 5’-CTTTGGTATCGTGGAAGGACTC-3’

GAPDH-reverse1: 5’-CTCTTCCTCTTGTGCTCTTGCT-3’

### Preparation of human brain homogenate

Approximately 100 mg human brain cerebellum was resuspended in 800 µl 50mM Tris pH: 7,5, 1/25 ROCHE Protease Inhibitor a, 1:1000 Benzonase. The suspension was transferred into a 2 ml Precellys Tube (CK 14 Ceramik beads, Ref. Nr. KTO 3961-1-003.2) and homogenised by Precellys Evolution with 5000rpm, 2x 14sec. The homogenisate was suplemented with 200µl 5x RIPA buffer in the 2 ml Precellys tube (5x RIPA: 250 mM Tris pH 7,5; 750 mMNaCl; 0,5 % SDS; 2 % Sodium Deoxycholate; 5 % Triton) and incubated for 30 min at 4°C in a rotating wheel. Debris was removed by centrifugation at 1500 g for 20 min at 4°C. Protein concentration was measured by using standard BCA protocol.

### Statistics

All data are shown as mean and standard deviation (s.d.) of multiple replicates/experiments (as indication in Fig. legends). Analysis of all experimental data was done with GraphPad Prism 5 (San Diego, CA). P values were calculated with two-sided, unpaired t-test following the tests. P values less than 0.05 were considered significant.

## Fig. legends

**Extended Data Fig. 1 Schematic representation of the domain structure of the human SCAN-domain protein family.**

**a,** Protein sequence alignment of the SCAN domain template c3lhrA_ (identified by Phyre2) and PGBD1 from various mammalian and marsupial species. **b,** Protein sequence alignment of the KRAB domain template d1v65a_ (identified by Phyre2) and PGBD1 from various mammalian and marsupial species. **c,** Schematic representation of the domain structure of the human SCAN-domain protein family of transcription factors. From each protein the longest isoform is presented, including the protein domain annotation from SMART and PFAM. Filled boxes indicate domains, which were annotated in both databases, continuous lines indicate domains, which are annotated in SMART, but not in PFAM and dashed lines indicate domains, which are annotated in PFAM but not in SMART. Note that in PGBD1 the ZNF_C2H2 domain is replaced by the transposase-derived domain. **d,** Phylogenetic tree of Pgbd1 and Pgbd2. Both Pgbd1 and Pgbd2 are specific to the Mammalian lineage, not including Monotemes (Therian). They are closely related but differ in their protein domain structure. All sequences (∼12k) containing the pfam domain Transposase IS4 have been downloaded from interproUniprot DB and aligned with mafft. An initial tree has been calculated with the UPGMA algorithm from which a subtree has been manually picked. The subtree includes the cluster of Pgbd1 and Pgbd2 plus some closely clustering sequences. Identical sequences and sequences shorter than 250 bp have been removed. The Pgbd1 and Pgbd2 sequences from Koala have been added manually. The picked transcripts were realigned using muscle and a phylogeny tree was build using MrBayes. Annotated protein domains originate from the pfam db. While Pgbd2 carries exclusively the IS4 transposase-like domain, Pgbd1 has an additional SCAN and occasionally KRAB domains at the N-terminal. Average pairwise similarity score of ∼ 63% of the aligned region which spans 1324 bp exceeding the borders of the annotated transposase IS4 domain, calculated by distance matrix of Ugene).

**Extended Data Fig. 2| EggNOG phylogeny analyses for PGBD1 and PGBD2.**

**a,** EggNOG phylogeny tree for PGBD1, PGBD2 and the closest relatives showing SMART domain predictions. Note that the KRAB domain is not predicted in *Homo sapiens*. **b,** EggNOG phylogeny analysis for PGBD1, PGBD2 and the closest relatives.

**Extended Data Fig. 3| The plasticity of PGBD1 evolution.**

**a,** abSENSE method was used to calculate the probability that homologs of PGBD1 and PGBD2 would fail to be detected by a homology search (using BLAST method) in platypus, alligator, lizard and frog. From the rate parameters of the two genes where orthologs are described (bitscore: y axis), given the evolutionary distances between these species (x axis), we can infer the probability that an ortholog is truly absent in other species, given their evolutionary distance from the focal species (human), rather than simply not findable. For all of the species without an identified ortholog we can reject the hypothesis that the ortholog would not be expected to be detected by homology search even if present. Thus, absence of an orthology can not be ascribed to failure of homology search. **b,** The transposase-derived domains of human (hPGBD1) and rat (rPgbd1) sequences are highly similar (87% identity). Amino acid sequence alignment of the human and rat transposase-derived domains of Pgbd1. **c,** Conservation of the rPgbd1 in the rat genome. Exon architecture of the human and rat Pgbd1 genes. The human hPGBD1 consists of 7 exons. The N-terminal (SCAN and KRAB) domains are not detectable in the rat. Arrows represent the positions of the PCR primers that were used to analyse the N-terminus of the rPgbd1. Sequences of the forward and reverse primer that were used in the final analysis are shown. **d,** PCR amplified DNA fragments were cloned into pJET1.2 vector (Fermentas) then the individually purified plasmids were digested by BglII restriction endonuclease then further analysed by agarose gel electrophoresis. DNA fragment of the lanes 1 and 6 PCR products were analysed by Sanger sequencing. **e,** The predicted amino acid sequences of two PCR products amplified from the rat genome (yellow) identify several STOP codons upstream of the transposase-derived domain (gray).

**Extended Data Fig. 4| Tissue specific gene expression profile of the human PGBD1 and PGBD2.**

**a,** Tissue specific gene expression profile of the human PGBD1 protein based on www.gtexportal.org. Note the enriched expression of PGBD1 in the cerebellum and cerebellar hemisphere. **b,** Tissue specific gene expression profile of the human PGBD2 proteins based on www.gtexportal.org. Note that PGBD2 is mostly enriched in spleen and thyroid tissues. **c,** Genome activation status of PGBD1 promoter (1kb upstream to TSS) in human embryonic stem cells (ECS) neural progenitor cells (NPC) and neurons, characterized by selected histone marks for active and poised states. Yellow lines represent the count per million (CPM) values at log2FC. **d,** PGBD1 expression is the highest in ESCs, followed by NPC and differentiated neuron. Transcript levels of PGBD1 in human embryonic stem cells (ESC), neural progenitor cells (NPC) and neurons (GSE118106). **e,** Expression level of the PGBD1 protein in ESC, NPC and differentiated neuron (Western blot). PGBD1:actin ratio is shown at the bottom. ESC (Embryonic stem cell H1_ESC), H1_mDAN1; human H1_ECS-derived neuron, N1H1; human H1_ECS derived neural progenitor cell (NPC). The numbers indicate the ratio of the endogenous PGBD1 and actin proteins (determined by densitometry analysis). **f,** PCR-based *transposonexcision repair assay* detects no activity of the PGBD1. (Upper panel) Schematic view of the PCR-based 2-step transposon excision assay in HEK-293 cells. Panel shows the position of the primers on the DNA plasmids that were used in the assay. In case of excision, the 2nd PCR step amplifies an approximately a 357 bp product. Luciferase (Luc) is used as a control protein. (Lower panel) Agarose gel electrophoresis image of the PCR products from the excision assay. Only the mouse *piggyBac* transposase (mPB) has detectable activity, indicated by the 374 bp PCR product. To monitor the efficiency of plasmid isolation, PCR on the plasmid backbone (on the ampicillin resistance gene sequence) was also carried out resulted in a 340 bp long PCR product.

**Extended Data Fig. 5| Genome-wide analyses of PGBD1 binding.**

**a,** Genome-wide distribution of the PGBD1 ChiP-exo peaks in the protein coding and non-protein coding regions in human NPCs and neurons. Note that the intronic PGBD1 ChiP-exo peaks also mapped to *AluSx* sequences (∼89.5%) in both NPCs and neurons (not shown). **b,** Dissection ofPGBD1 binding sites intersecting with different subfamilies of *Alu* elements in NPCs. Note that 32% of the *AluSx* elements form and inverted repeat structure in the 3’UTR exons. **c,** Comparative plot of the significant (p<0,05) Gene Ontology terms (Biological Function) of the target protein coding genes. **d,** Correlation plot of PGBD1 ChIP-exo peaks and ATAC-seq signals from NPCs and differentiated neurons (GSE95023). **e,** Heatmap of PGBD1 binding sites merged with various histone marks genome wide in NPCs or in neurons in NPCs and in neurons (1kb upstream from RefSeq annotated TSS). Note that the bottom panels are representing PGBD1 binding peaks merged H3K4me1 signals (2kb upstream from RefSeq annotated TSS).**f,** Active and repressed gene promoters intersecting with PGBD1 binding sites.

**Extended Data Fig. 6| Depletion of PGBD1 compromises the NPC identity.**

**a,** Representative images of knock-out (KO) CRISPR cells. Note that none of the used gRNA constructs was suitable to establish a stable KO-PGBD1 cell line. Long-term (two months) culturing resulted in a differentiated morphology. **b,** Determination of the endogenous PGBD1 protein levels following depletion in NPC. (Left panel) Total protein visualized by the BioRad-ChemiDoc™ MP Imaging System. (Right panel) Western blot. The quantification was performed by normalizing the PGBD1 level with the total protein amount. KD-efficiency of the unique samples is indicated in %. **c,** Stable knock-down (KD) and overexpressing (OE) of PGBD1 in neuroblastoma SHEP cells. (Left panels) Representative confocal microscopic images of stable miRNA based KD-PGBD1 cell lines (KD1-PGBD1, KD2-PGBD1) used in functional studies. Scr, scrambled control. mCherry is a marker of the integrated KD miRNA constructs in the stable SHEP cells. (Right panel) Representative confocal microscopy image shows the overexpressed HA-tagged PGBD1 protein in a stable human SHEP cell line that was used in functional experiments. **d,** (Left panels) Western blot analysis of the PGBD1 depletion in stable KD-PGBD1 SHEP cell lines. Red boxes mark the selected lines used in further studies. Numbers indicate the KD efficiency determined by the ration of endogenous PGBD1 and actin. (Right panel) Western blot analysis of stable TET-inducible SHEP cell lines overexpressing HA-tagged PGBD1 protein in presence and absence of doxycycline. IB. immunoblot; aPGBD1: anti-PGBD1 antibody; aActin: anti-*α*-actinin antibody. **e,** Depletion of PGBD1 compromises the NPC identity and results in neuronal differentiation. HeatMap demonstrates the comparison of a gene expression of a selected neuronal lineage marker set (n=99) (https://www.rndsystems.com/research-area/neural-stem-cell-and-differentiation-markers) in PGBD1-depleted neural progenitor cells (KD-PGBD1_NPCs) and differentiated neurons.

**Extended Data Fig. 7| MS-SILAC analysis of the PGBD1 proteome.**

**a,** The highly confident PGBD1 interactome. Scatter plot of the H/L ratios of the SILAC based AP-MS experiments (label-switch) in HEK293 cells.The normalized H/L ratios are log and z transformed. Significance A was calculated according to^208^ with modifications (see methods). The red dots represent the highly significant interacting partners (FDR < 0.05), whereasthe black dots are considered as background. Note that the approach does not distinguish between HSAPA6/HSPA7 or DTNA/DTNB, as the detected peptide is shared between both proteins. **b,** Relative transcriptional changes of interactors upon knockdown and overexpression of PGBD1. Heatmap of the TPM normalized expression from RNA-seq in wild type, PGBD1 knockdown and PGBD1 overexpression NPCs. Only interactors with > 10 TPM in wild type NPCs are shown in the heatmap. **c,** Enriched gene sets in the PGBD1 interactome (Molecular Signature Database v7.2). The colour intensities of the dots indicate the P-value, the size of the dot represents the effect size.

**Extended Data Fig. 8| Elevated NEAT1_2 transcription in stable, PGBD1-depleted SHEP cells.**

Elevated NEAT1_2 transcription in stable, PGBD1-depleted SHEP cells (KD1-PGBD1, KD2-PGBD1). The NEAT1_2 transcripts (gold) are visualized by fluorescent in situ hybridization (FISH), whereas PGBD1 (green) with immunostaining. blue, DAPI. Representative confocal microscopic images (mCherry-KD-Scr) miRNA scrambled control, Ctrl SHEP-non-transfected SHEP cells). mCherry is a marker of the integrated KD miRNA constructs in the stable SHEP cells (see Extended Data Fig. 9).

**Extended Data Fig. 9| Formation of ATXN1(Q30) aggregates in stable, PGBD1-depleted SHEP cells.**

**a,** Formation of EGFP-tagged ATXN1(Q30) aggregates in stable KD-PGBD1 SHEP cells. Representative confocal microscopy images. Arrows point to the ATXN1 aggregates. mCherry is a marker of the integrated KD miRNA constructs in the stable SHEP cells. mCherry-KD-Scr, scrambled control miRNA. **b,** Formation of EGFP-tagged ATXN1(Q30) aggregates in stable SHEP cells, where PGBD1 was overexpressed (OE) from a TET-inducible (Dox+) expression construct. Representative confocal microscopy images. Arrows point to the ATXN1 aggregates. **c,** Quantification of the observed aggregates in the confocal microscopy study. Student’s t-test was used for statistical comparisons, *** p<0.0005. Scale 20μm.

**Extended Data Fig. 10| Formation of ATXN1(Q2) aggregates in stable, PGBD1-depleted SHEP cells.**

**a,** No physical interaction is detectable between PGBD1 and EGFP-ATXN1(Q2) proteins in SHEP cells by co-immunoprecipitation. IB, immunoblot; aPGBD1, anti-PGBD1 antibody; aGFP, antiGFP antibody. **b,** Formation of EGFP-tagged ATXN1(Q2) aggregates in stable KD-PGBD1 SHEP cells. Representative confocal microscopy images. Arrows point to the ATXN1 aggregates. **c,** Formation of EGFP-tagged ATXN1(Q2) aggregates in stable SHEP cells, where PGBD1 was overexpressed (OE) from a TET-inducible (Dox+) expression construct. Representative confocal microscopy images. Arrows point to the ATXN1 aggregates. **d,** Quantification of the observed aggregates in the confocal microscopy study. Student’s t-test, NS: not significant at p<0.05. Scale 20μm.

**Extended Data Fig. 11| The quantitative dichloro-dihydro-fluorescein diacetate (DCFH-DA) assay to determine cellular oxidative stress.**

**a,** Schematic representation of the quantitative dichloro-dihydro-fluorescein diacetate (DCFH-DA) assay to determine cellular oxidative stress. **b,** Representative FACS analysis of the fluorescence emission (equivalent with GFP fluorescence emission) generated by the DCFH dye in human neuroblastoma SHEP cells.

---

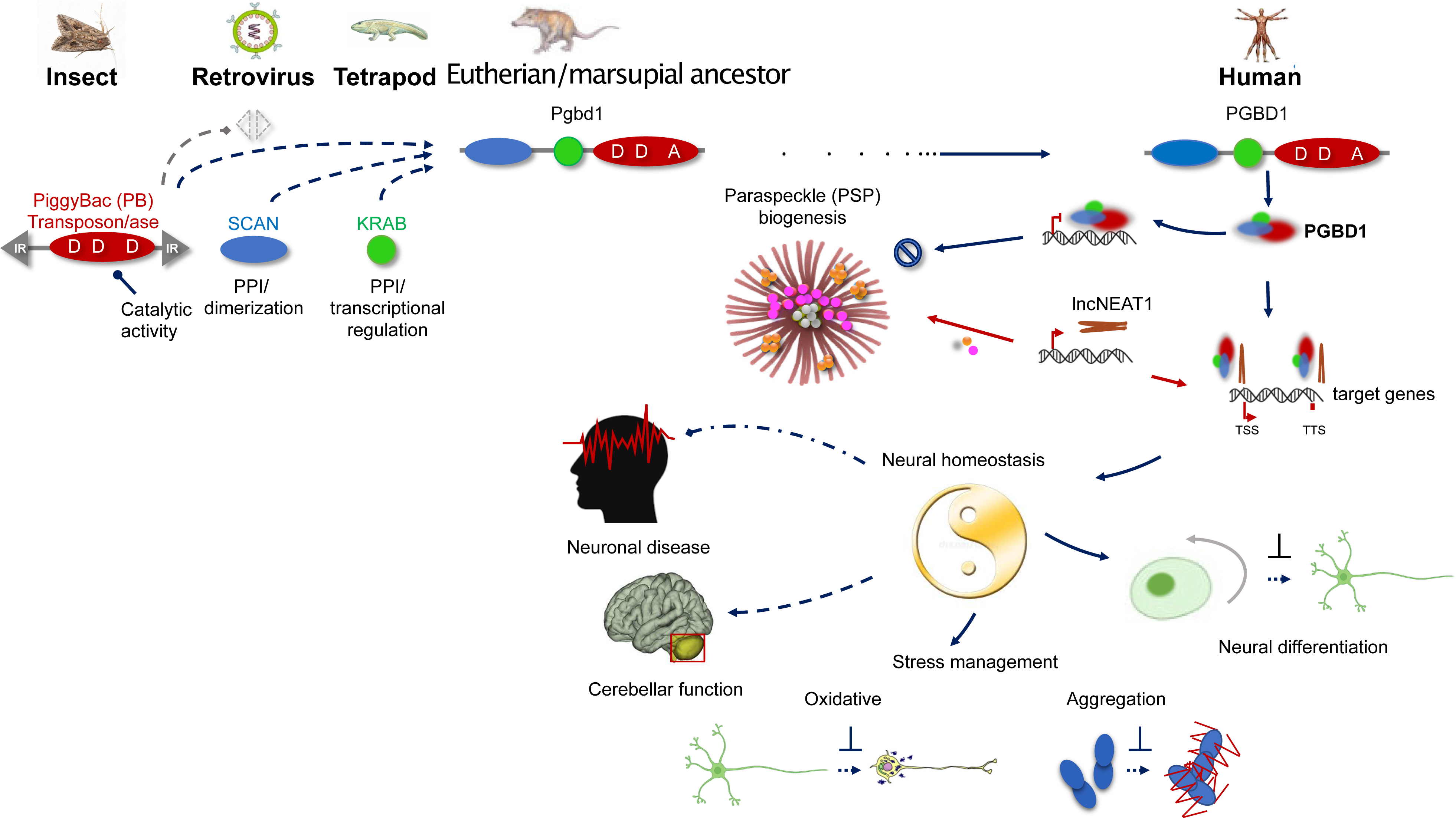

