## Supplementary material for "PiggyBac Transposable Element-derived 1 controls Neuronal Progenitor Identity, Stress Sensing and mammal-specific paraspeckles": PGBD1_supplementary

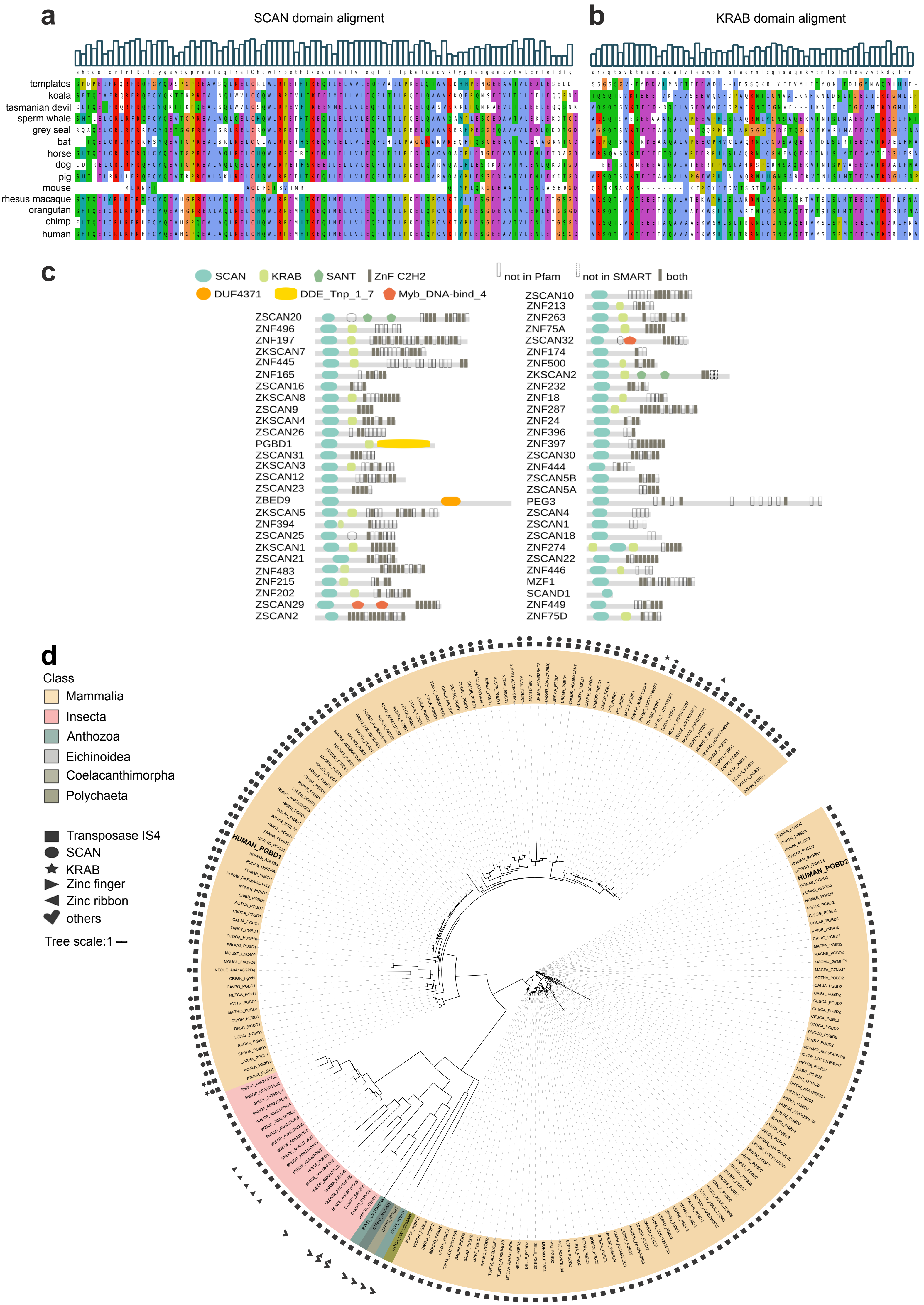

a
SCAN domain aligment
b
KRAB domain aligment
templates
koala
tasmanian devil
sperm whale
grey seal
bat
horse
dog
pig
mouse
rhesus macaque
orangutan
chimp
human
c
SCAN
KRAB
SANT
ZnF C2H2
not in Pfam
not in SMART
both
DUF4371
DDE_Tnp_1_7
Myb_DNA-bind_4
ZSCAN20
ZNF496
ZNF197
ZKSCAN7
ZNF445
ZNF165
ZSCAN16
ZKSCAN8
ZSCAN9
ZKSCAN4
ZSCAN26
PGBD1
ZSCAN31
ZKSCAN3
ZSCAN12
ZSCAN23
ZBED9
ZKSCAN5
ZNF394
ZSCAN25
ZKSCAN1
ZSCAN21
ZNF483
ZNF215
ZNF202
ZSCAN29
ZSCAN2
ZSCAN10
ZNF213
ZNF263
ZNF75A
ZSCAN32
ZNF174
ZNF500
ZKSCAN2
ZNF232
ZNF18
ZNF287
ZNF24
ZNF396
ZNF397
ZSCAN30
ZNF444
ZSCAN5B
ZSCAN5A
PEG3
ZSCAN4
ZSCAN1
ZSCAN18
ZNF274
ZSCAN22
ZNF446
MZF1
SCAND1
ZNF449
ZNF75D
d
Class
Mammalia
Insecta
Anthozoa
Eichinoidea
Coelacanthimorpha
Polychaeta
Transposase IS4
SCAN
KRAB
Zinc finger
Zinc ribbon
others
Tree scale:1
HUMAN_PGBD1
HUMAN_PGBD2

Figure S2

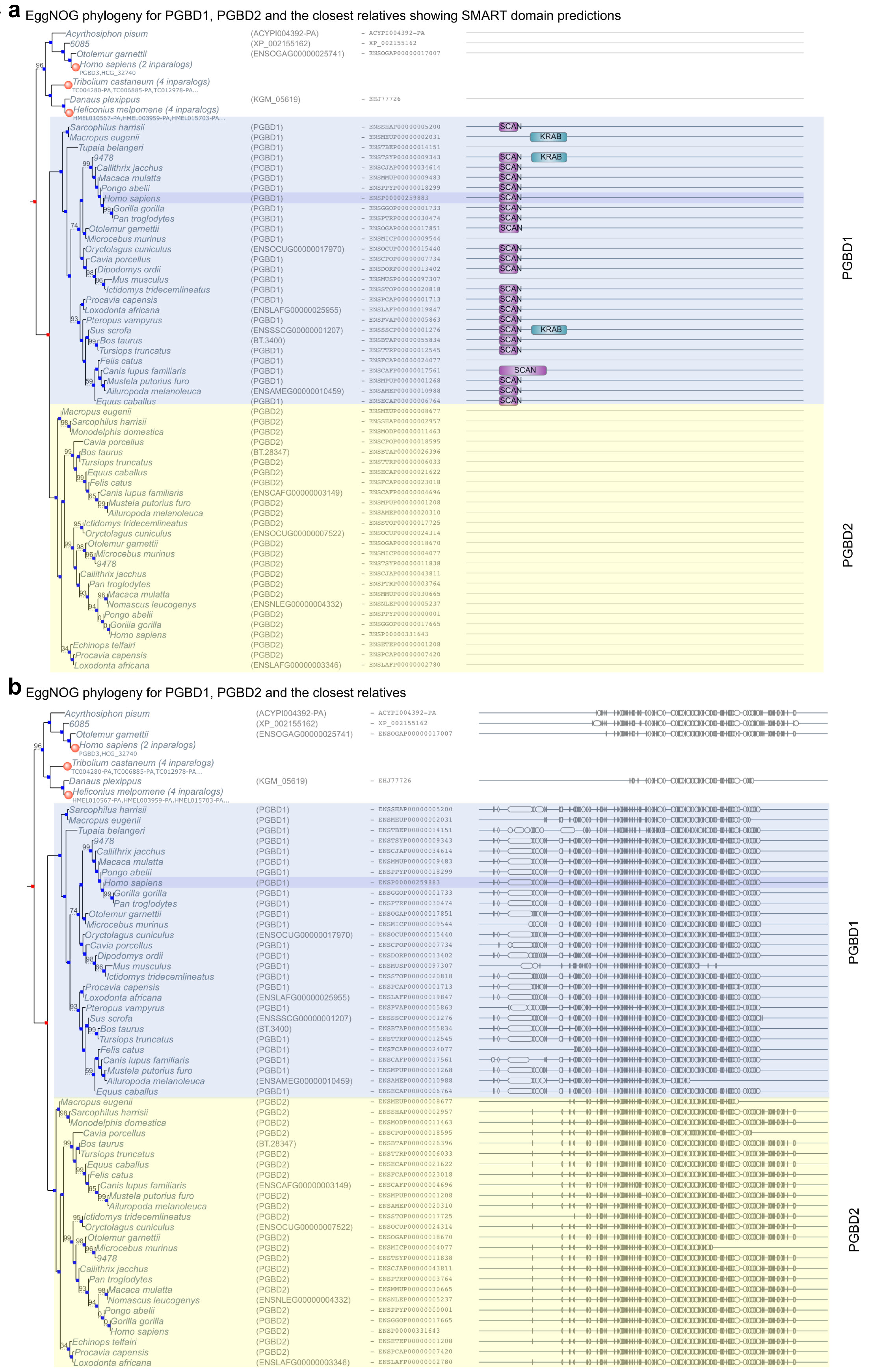
a
EggNOG phylogeny for PGBD1, PGBD2 and the closest relatives showing SMART domain predictions
SCAN
KRAB
PGBD1
PGBD2
b
EggNOG phylogeny for PGBD1, PGBD2 and the closest relatives
PGBD1
PGBD2

**a**

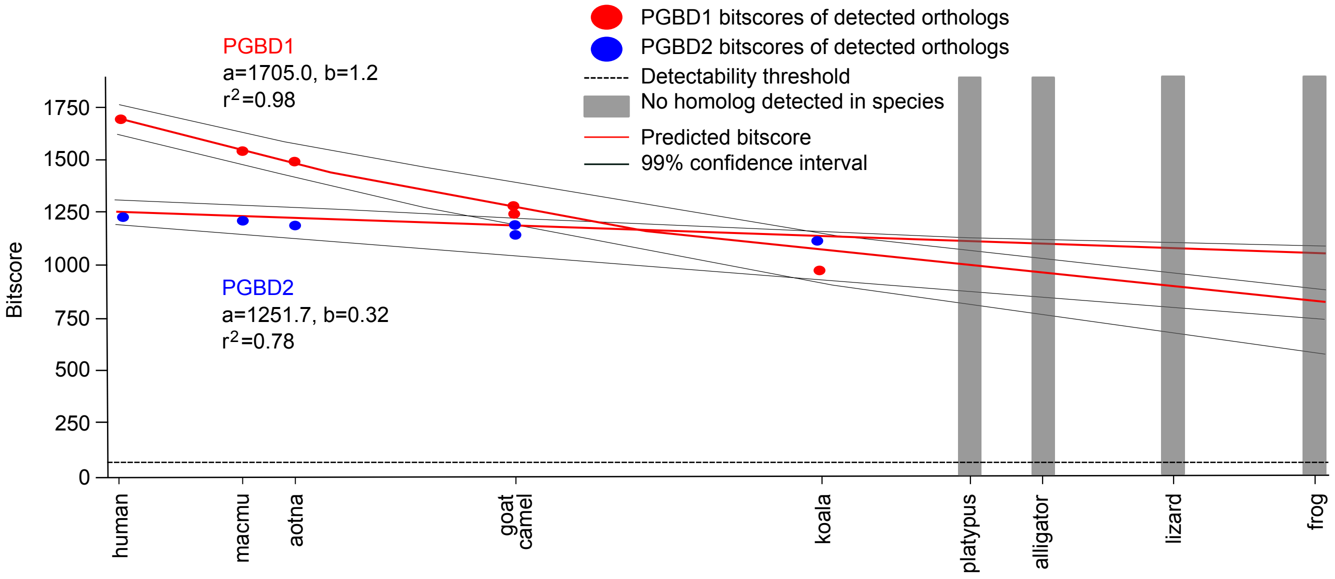


| Species | Homolog bitscore | |
|---|---|---|
| | PGBD1 | PGBD2 |
| Human | 1693.0 | 1241.0 |
| macmu | 1608.0 | 1233.0 |
| aotna | 1543.0 | 1207.0 |
| goat | 1264.0 | 1199.0 |
| camel | 1349.0 | 1099.0 |
| koala | 1059.0 | 1081.0 |
| platypus | No homolog detected | |
| alligator | No homolog detected | |
| lizard | No homolog detected | |
| frog | No homolog detected | |

| Species missing a detected homolog | Probability of homolog detection failure (E=0.001) | | Predicted bitscore | | 99 percent confidence interval | |
|---|---|---|---|---|---|---|
| | PGBD1 | PGBD2 | PGBD1 | PGBD2 | PGBD1 | PGBD2 |
| platypus | 0.0 | 0.0 | 1565.02 | 1223.6 | 1468.66 - 1628.14 | 1174.45 - 1294.35 |
| alligator | 0.0 | 0.0 | 1294.59 | 1163.64 | 1176.2 -1383.83 | 1091.5 - 1258.48 |
| lizard | 0.0 | 0.0 | 947.44 | 1071.3 | 799.27 - 1069.29 | 951.41 - 1213.76 |
| frog | 0.0 | 0.0 | 900.65 | 1057.03 | 742.89 - 1025.36 | 930.48 - 1204.94 |

**b**

```
hPGBD1  457  MRCVFGVLLLSGFMRHPRREMYWEVSDTDQNLVRDAIRRDRFELIFSNLHFADNGHLDQK  516
             M CVFGVLL SGF+ HPR  MYWE+SD+DQ LVR+AIRRDRFELIFS LHFA N HL QK
rPgbd1  1    MWCVFGVLLWSGFVMHPRMGMYWEISDSDQTLVRNAIRRDRFELIFSYLHFAGNSHLHQK  60

hPGBD1  517  DKFTKLRPLIKQMNKNFLLYAP-LEEYYCFDKSMCECFDSDQFLNGKPIRIGYKIWCGTT  575
             DKF+ LRPLIKQMNKNFLLYAP LEEYYCFDKSMCECFDSDQFLNGKP+RIGYKIWCGTT
rPgbd1  61   DKFSILRPLIKQMNKNFLLYAPRLEEYYCFDKSMCECFDSDQFLNGKPLRIGYKIWCGTT  120

hPGBD1  576  TQGYLVWFEPYQEESTMKVDEDPDLGLGGNLVMNFADVLLERGQYPYHLCFDSFFTSVKL  635
             TQGYLVWFEPYQE S ++ D++ DLGLGGNL+M+FADVLLE+G YPYHLCF+SFFTSVKL
rPgbd1  121  TQGYLVWFEPYQEYSAVETDKELDLGLGGNLIMSFADVLLEKGHYPYHLCFESFFTSVKL  180

hPGBD1  636  LSALKKKGVRATGTIRENRTEKCPLMNVEHMKKMKRGYFDFRIEENNEIILCRWYGDGII  695
             +SALKKKGV+ATG+IRENR EKCPLMNVEHMKKMKRG+F+FR+EEN+EI L  W+GD  I
rPgbd1  181  MSALKKKGVKATGSIRENRMEKCPLMNVEHMKKMKRGHFNFRVEENDEIFLFHWHGDSFI  240

hPGBD1  696  SLCSNAVGIEPVNEVSCCDADNEEIPQISQPSIVKVYDECKEGVAKMDQIISKYRVRIRS  755
             SLCSNAVGIEPV+E+SC  A+ +  PQ+SQPSIV +Y++C++GVAKMDQIIS+YRV +RS
rPgbd1  241  SLCSNAVGIEPVSEISCV-ANGKASPQVSQPSIVNLYEKCRKGVAKMDQIISRYRVGLRS  299

hPGBD1  756  KKWYSILVSYMIDVAMNNAWQLHRACNPGASLDPLDFRRFVAHFYLEHNAHLSD Stop  809
             KK    LVSYMI+VAMNNAWQLHR CNPG+ LD L F + VA FYL H+ +LSD
rPgbd1  300  KKRSLGLVSYMINVAMNNAWQLHRICNPGSPLDLLGFWKCVACFYLGHDINLSD Stop  353
```

**c**

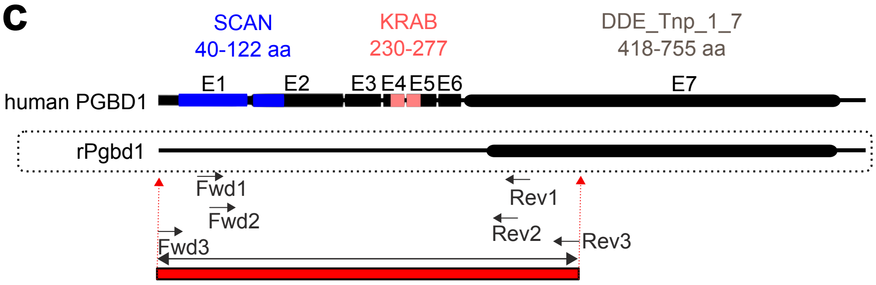


PCR fragment amplified from rat genomic DNA by Fwd3-Rev3 primers ~1200 bp

Fwd3: 5'- ATG TAT GAA GTT TTG CCG GGT -3'
Rev3: 5'- TCG ATC CTG TAG CCT TCA CTC -3'

**d**

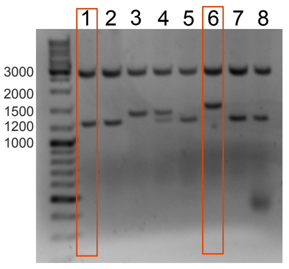


amplified PCR fragments cloned into pJET1.2 vector, plasmids were digested by BglII enzyme

**e**

PCR product 1

```
C I F Stop _ F R G R G R Q Stop P Stop G A A Stop E K A K G I Q H A Stop E E L
D E K R H Stop A Q L S K L V S T G F W T F E S Q K Stop E V K P S R A L Stop I
I F Stop _ Stop N I Q L N Stop I Q Stop N Q Stop F N E T N N Y A S Q K N V N L
E V T L Q E M W C V F G V L L W S G F V M H P R M G M Y W E I S D S D Q T L V
R N A I R R D R F E L I F S Y L H F A G N S H L H Q K D K F S I L R P L I K Q
M N K N F L L Y A P R L E E Y Y C F D K S M C E C F D S D Q F L N G K P L R I
G Y K I W C G T T T Q G Y L V W F E P Y Q E Y S A V E T D K E L D L G L G G N
L I M S F A D V L L E K G H Y P Y H L C F E S F F T S V K L M S A L K K K G V
K A T G S I R E N R M E K C P L M N V E H M K K M K R G H F N F R V E E N D E
I F L F H W H G D S F I S L C S N A V G I E P V S E I S C V A N G K A S P Q V
S Q P S I V N L Y E K C R K G V A K M D Q I I S R Y R V G L R S K K R S L G L
V S Y M I N V A M N N A W Q L H R I C N P G S P L D L L G F W K C V A C F Y L
G H D I N L S D Stop
```

PCR product 6

```
Stop _ F R G R G R Q Stop P Stop G A A Stop E K A K G I Q H A Stop E E L D E K
K H Stop A Q L S K L V S T G F W T F E S Q K Stop E V K P S R A L Stop I I F
Stop _ Stop N I Q L N Stop I Q Stop N Q Stop F N E T N N Y A S Q K N V N L E V
T L Q E M W C V F G V L L W S G F V M H P R M G M Y W E I S D S D Q T L V R N
A I R R D R F E L I F S Y L H F A G N S H L H Q K D K F S I L R P L I K Q M N
K N F L L Y A P R L E E Y Y C F D K S M C E C F D S D Q F L N G K P L R I G Y
K I W C G T T T Q G Y L V W F E P Y Q E Y S A V E T D K E L D L G L G G N L I
M S F A D V L L E K G H Y P Y H L C F E S F F T T L N X M S A L K K K G V K A
T G S I R E N R M E K C P L M N V E H M K K M K R G H F N F R V E E N D E I F
L F H W H G D S F I S L C S N A V G I E P V S E I S C V A N G K A S P Q V S Q
P S I V N L Y E K C R K G V A K M D Q I I S R Y R V G L R S K K R S L G L V S
Y M I N V A M N N A W Q L H R I C N P G S P L D L L G F W K C V A C F Y L G H
D I N L S D Stop
```

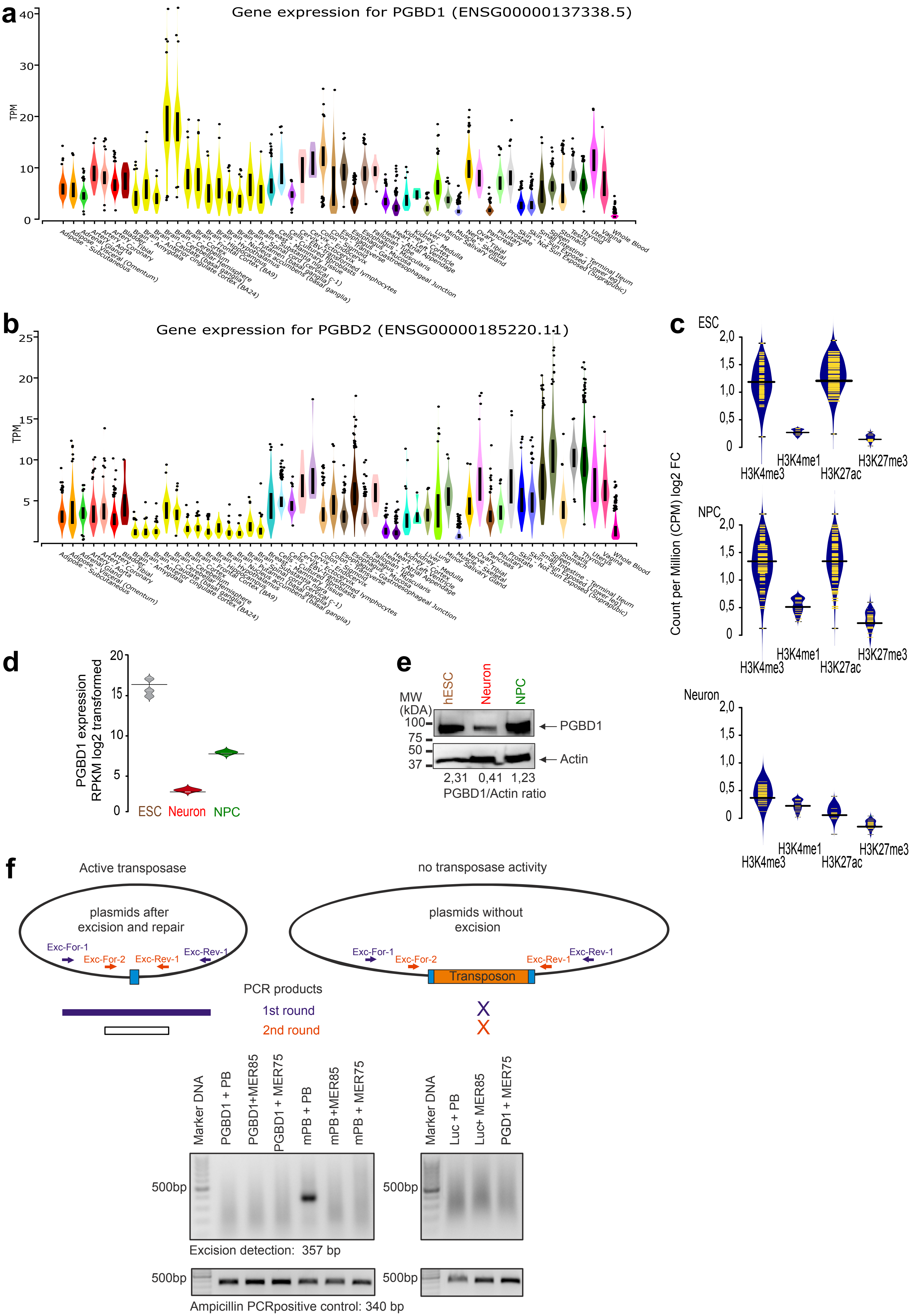
a
Gene expression for PGBD1 (ENSG00000137338.5)
b
Gene expression for PGBD2 (ENSG00000185220.11)
c
ESC
NPC
Neuron
Count per Million (CPM) log2 FC
H3K4me3
H3K4me1
H3K27ac
H3K27me3
d
PGBD1 expression RPKM log2 transformed
ESC
Neuron
NPC
e
MW (kDA)
hESC
Neuron
NPC
PGBD1
Actin
2,31 0,41 1,23
PGBD1/Actin ratio
f
Active transposase
no transposase activity
plasmids after excision and repair
plasmids without excision
Exc-For-1
Exc-For-2
Exc-Rev-1
Exc-Rev-1
Transposon
PCR products
1st round
2nd round
Marker DNA
PGBD1 + PB
PGBD1+MER85
PGBD1 + MER75
mPB + PB
mPB +MER85
mPB + MER75
Luc + PB
Luc+ MER85
PGD1 + MER75
500bp
Excision detection: 357 bp
Ampicillin PCRpositive control: 340 bp

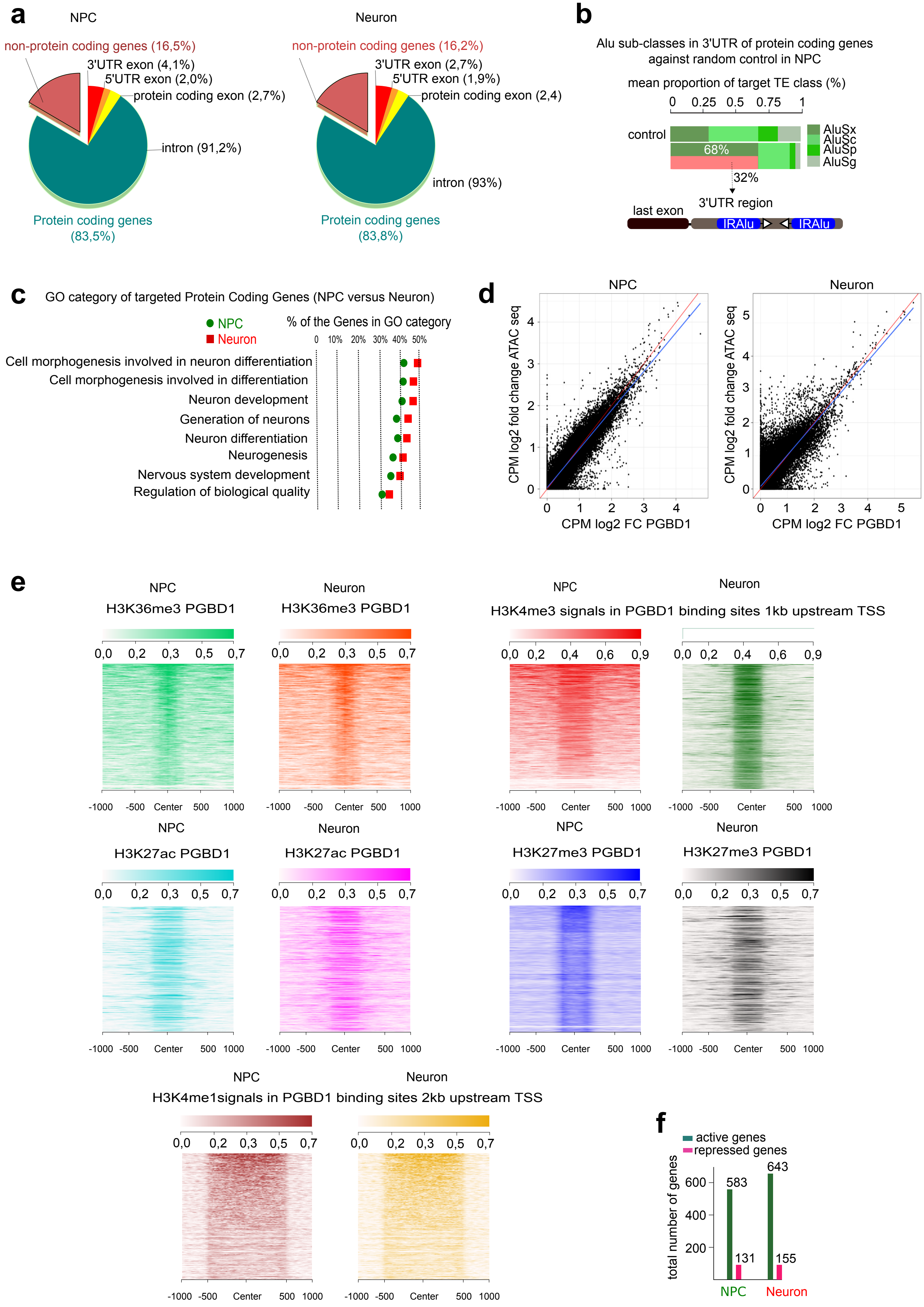
a
NPC
non-protein coding genes (16,5%)
3'UTR exon (4,1%)
5'UTR exon (2,0%)
protein coding exon (2,7%)
intron (91,2%)
Protein coding genes
(83,5%)
Neuron
non-protein coding genes (16,2%)
3'UTR exon (2,7%)
5'UTR exon (1,9%)
protein coding exon (2,4)
intron (93%)
Protein coding genes
(83,8%)
b
Alu sub-classes in 3'UTR of protein coding genes
against random control in NPC
mean proportion of target TE class (%)
0 0.25 0.5 0.75 1
control
68%
32%
AluSx
AluSc
AluSp
AluSg
3'UTR region
last exon
IRAlu
IRAlu
c
GO category of targeted Protein Coding Genes (NPC versus Neuron)
NPC
Neuron
% of the Genes in GO category
0 10% 20% 30% 40% 50%
Cell morphogenesis involved in neuron differentiation
Cell morphogenesis involved in differentiation
Neuron development
Generation of neurons
Neuron differentiation
Neurogenesis
Nervous system development
Regulation of biological quality
d
NPC
Neuron
CPM log2 fold change ATAC seq
CPM log2 FC PGBD1
e
NPC
H3K36me3 PGBD1
Neuron
H3K36me3 PGBD1
NPC
Neuron
H3K4me3 signals in PGBD1 binding sites 1kb upstream TSS
0,0 0,2 0,3 0,5 0,7
0,0 0,2 0,4 0,6 0,9
-1000 -500 Center 500 1000
NPC
H3K27ac PGBD1
Neuron
H3K27ac PGBD1
NPC
H3K27me3 PGBD1
Neuron
H3K27me3 PGBD1
NPC
Neuron
H3K4me1signals in PGBD1 binding sites 2kb upstream TSS
f
active genes
repressed genes
total number of genes
583
131
643
155
NPC
Neuron

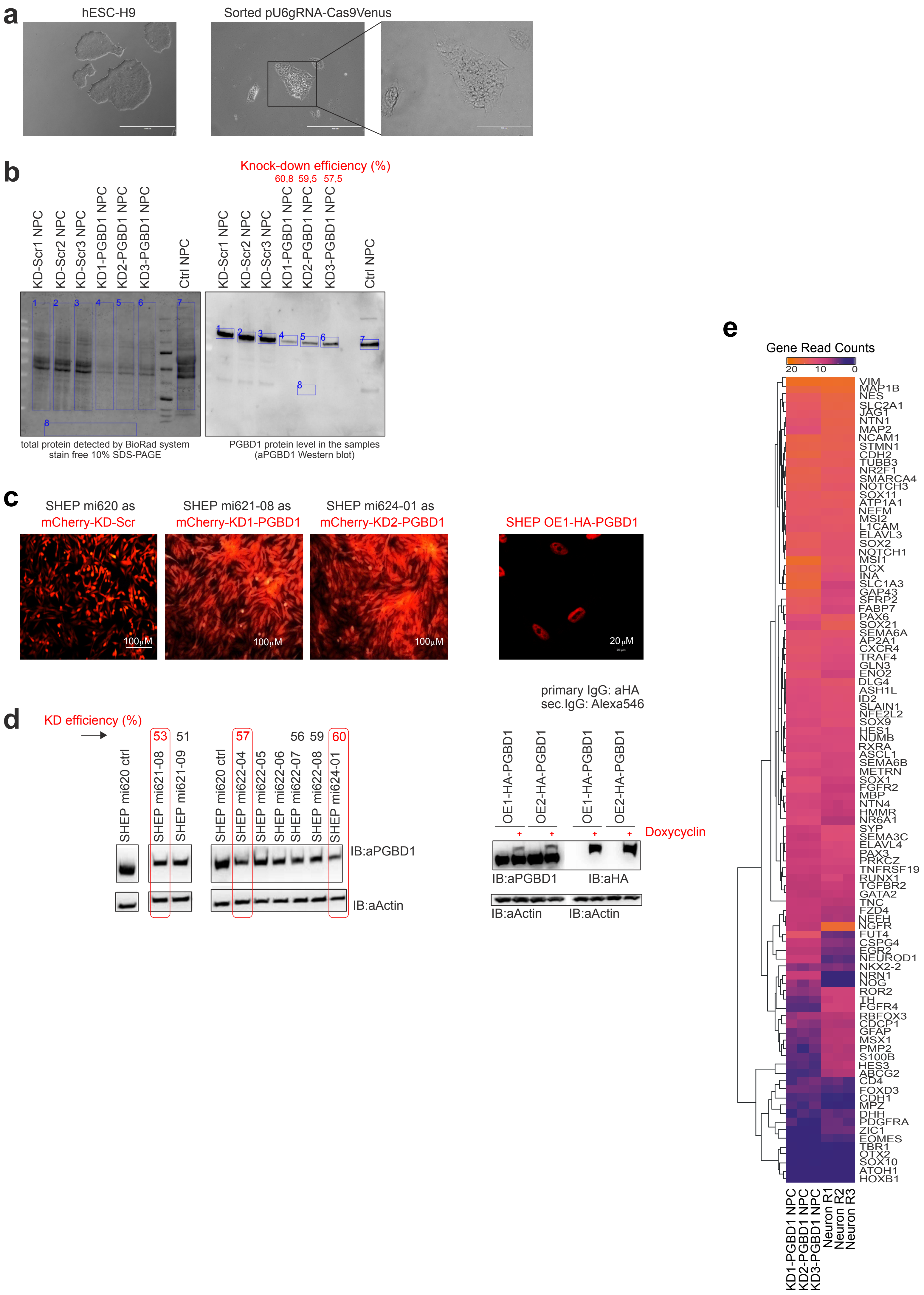
a
hESC-H9
Sorted pU6gRNA-Cas9Venus
b
Knock-down efficiency (%)
60,8 59,5 57,5
KD-Scr1 NPC
KD-Scr2 NPC
KD-Scr3 NPC
KD1-PGBD1 NPC
KD2-PGBD1 NPC
KD3-PGBD1 NPC
Ctrl NPC
total protein detected by BioRad system stain free 10% SDS-PAGE
PGBD1 protein level in the samples (aPGBD1 Western blot)
c
SHEP mi620 as mCherry-KD-Scr
SHEP mi621-08 as mCherry-KD1-PGBD1
SHEP mi624-01 as mCherry-KD2-PGBD1
SHEP OE1-HA-PGBD1
100μM
20μM
primary IgG: aHA
sec.IgG: Alexa546
d
KD efficiency (%)
53 51 57 56 59 60
SHEP mi620 ctrl
SHEP mi621-08
SHEP mi621-09
SHEP mi620 ctrl
SHEP mi622-04
SHEP mi622-05
SHEP mi622-06
SHEP mi622-07
SHEP mi622-08
SHEP mi624-01
IB:aPGBD1
IB:aActin
OE1-HA-PGBD1
OE2-HA-PGBD1
Doxycyclin
IB:aHA
e
Gene Read Counts
20 10 0
VIM
MAP1B
NES
SLC2A1
JAG1
NTN1
MAP2
NCAM1
STMN1
CDH2
TUBB3
NR2F1
SMARCA4
NOTCH3
SOX11
ATP1A1
NEFM
MSI2
L1CAM
ELAVL3
SOX2
NOTCH1
MSI1
DCX
INA
SLC1A3
GAP43
SFRP2
FABP7
PAX6
SOX21
SEMA6A
AP2A1
CXCR4
TRAF4
GLN3
ENO2
DLG4
ASH1L
ID2
SLAIN1
NFE2L2
SOX9
HES1
NUMB
RXRA
ASCL1
SEMA6B
METRN
SOX1
FGFR2
MBP
NTN4
HMMR
NR6A1
SYP
SEMA3C
ELAVL4
PAX3
PRKCZ
TNFRSF19
RUNX1
TGFBR2
GATA2
TNC
FZD4
NEFH
NGFR
FUT4
CSPG4
EGR2
NEUROD1
NKX2-2
NRN1
NOG
ROR2
TH
FGFR4
RBFOX3
CDCP1
GFAP
MSX1
PMP2
S100B
HES3
ABCG2
CD4
FOXD3
CDH1
MPZ
DHH
PDGFRA
ZIC1
EOMES
TBR1
OTX2
SOX10
ATOH1
HOXB1
KD1-PGBD1 NPC
KD2-PGBD1 NPC
KD3-PGBD1 NPC
Neuron R1
Neuron R2
Neuron R3

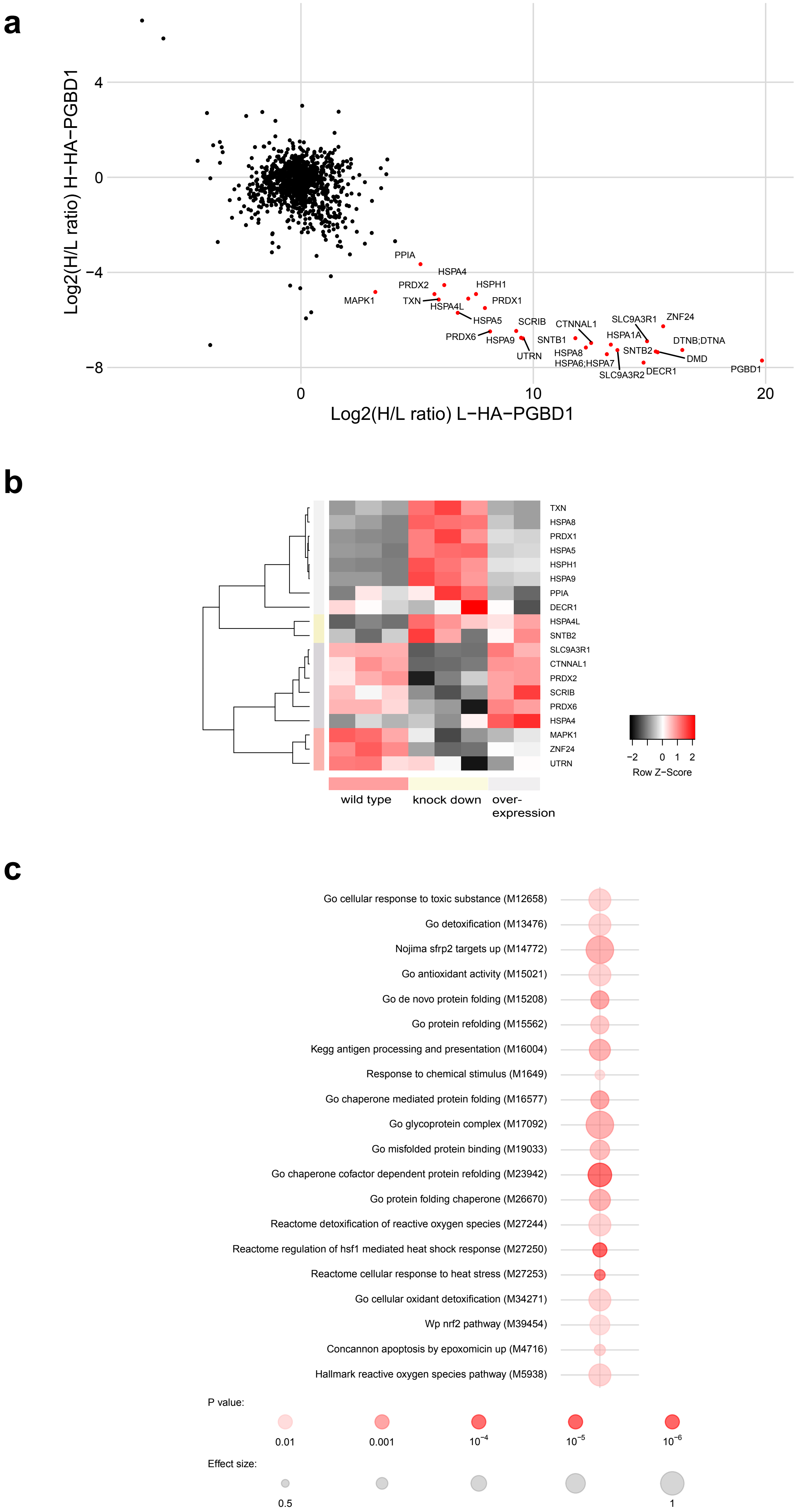
a
Log2(H/L ratio) H−HA−PGBD1
Log2(H/L ratio) L−HA−PGBD1
PPIA
HSPA4
PRDX2
HSPH1
MAPK1
TXN
HSPA4L
PRDX1
HSPA5
SCRIB
CTNNAL1
SLC9A3R1
ZNF24
PRDX6
HSPA9
SNTB1
HSPA1A
DTNB;DTNA
UTRN
HSPA8
SNTB2
DMD
HSPA6;HSPA7
SLC9A3R2
DECR1
PGBD1
b
TXN
HSPA8
PRDX1
HSPA5
HSPH1
HSPA9
PPIA
DECR1
HSPA4L
SNTB2
SLC9A3R1
CTNNAL1
PRDX2
SCRIB
PRDX6
HSPA4
MAPK1
ZNF24
UTRN
Row Z−Score
wild type
knock down
over-
expression
c
Go cellular response to toxic substance (M12658)
Go detoxification (M13476)
Nojima sfrp2 targets up (M14772)
Go antioxidant activity (M15021)
Go de novo protein folding (M15208)
Go protein refolding (M15562)
Kegg antigen processing and presentation (M16004)
Response to chemical stimulus (M1649)
Go chaperone mediated protein folding (M16577)
Go glycoprotein complex (M17092)
Go misfolded protein binding (M19033)
Go chaperone cofactor dependent protein refolding (M23942)
Go protein folding chaperone (M26670)
Reactome detoxification of reactive oxygen species (M27244)
Reactome regulation of hsf1 mediated heat shock response (M27250)
Reactome cellular response to heat stress (M27253)
Go cellular oxidant detoxification (M34271)
Wp nrf2 pathway (M39454)
Concannon apoptosis by epoxomicin up (M4716)
Hallmark reactive oxygen species pathway (M5938)
P value:
0.01
0.001
Effect size:
0.5
1

Figure S8

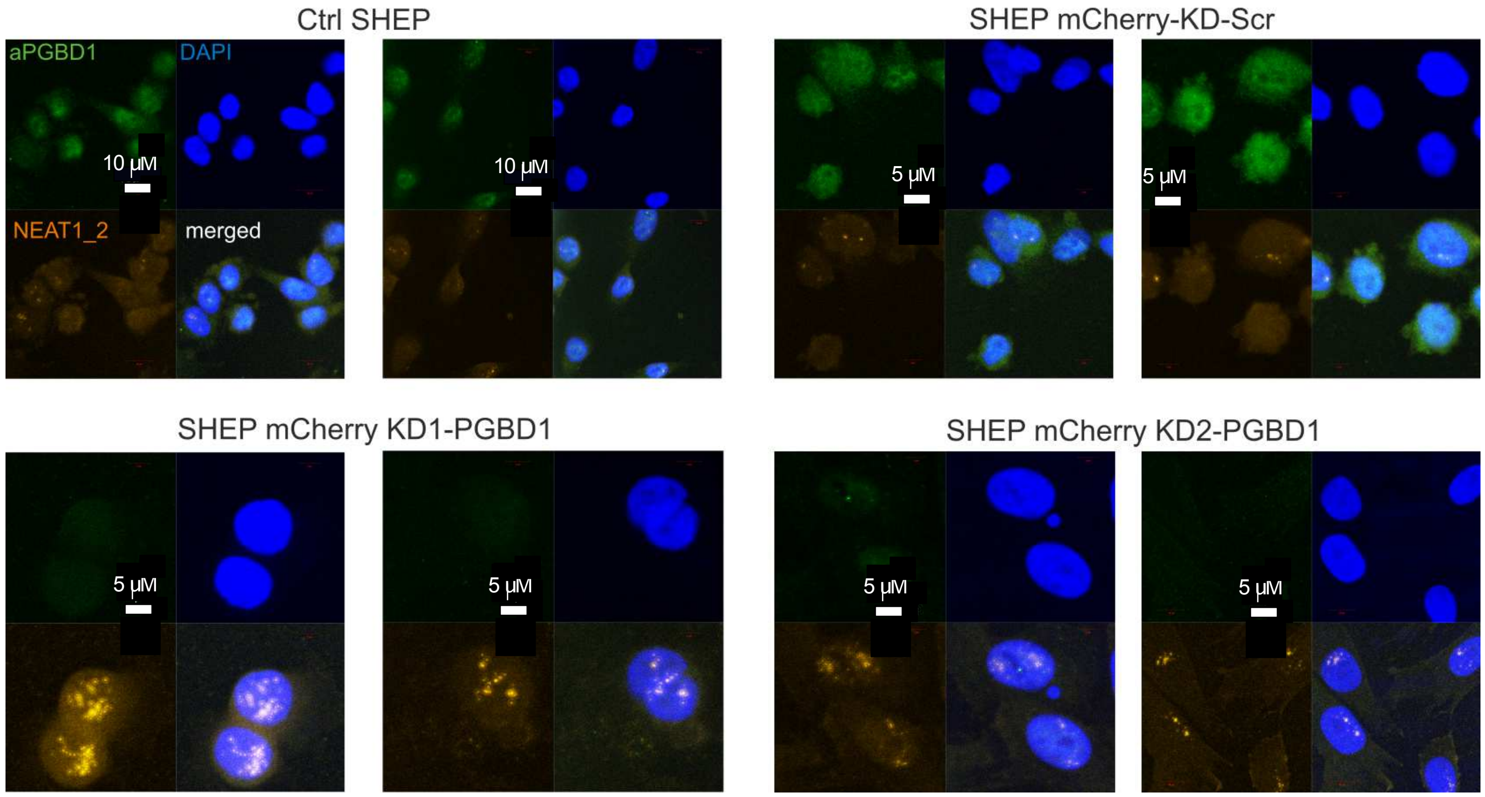
Ctrl SHEP
SHEP mCherry-KD-Scr
aPGBD1
DAPI
NEAT1_2
merged
10 µM
10 µM
5 µM
5 µM
SHEP mCherry KD1-PGBD1
SHEP mCherry KD2-PGBD1
5 µM
5 µM
5 µM
5 µM

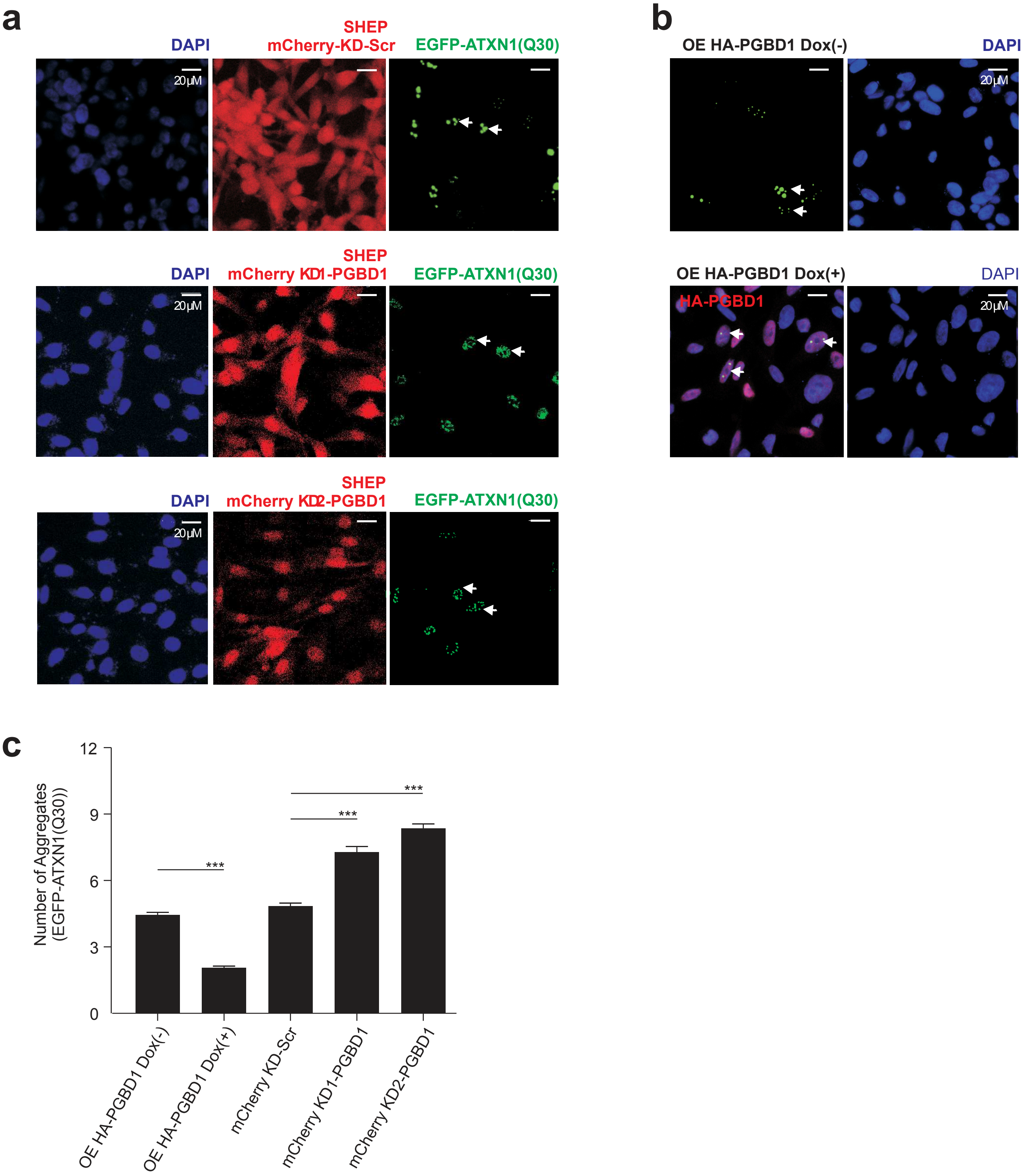
a
DAPI
SHEP
mCherry-KD-Scr
EGFP-ATXN1(Q30)
20μM
DAPI
SHEP
mCherry KD1-PGBD1
EGFP-ATXN1(Q30)
20μM
DAPI
SHEP
mCherry KD2-PGBD1
EGFP-ATXN1(Q30)
20μM
b
OE HA-PGBD1 Dox(-)
DAPI
20μM
OE HA-PGBD1 Dox(+)
DAPI
HA-PGBD1
20μM
c
Number of Aggregates (EGFP-ATXN1(Q30))
12
9
6
3
0
***
***
***
OE HA-PGBD1 Dox(-)
OE HA-PGBD1 Dox(+)
mCherry KD-Scr
mCherry KD1-PGBD1
mCherry KD2-PGBD1

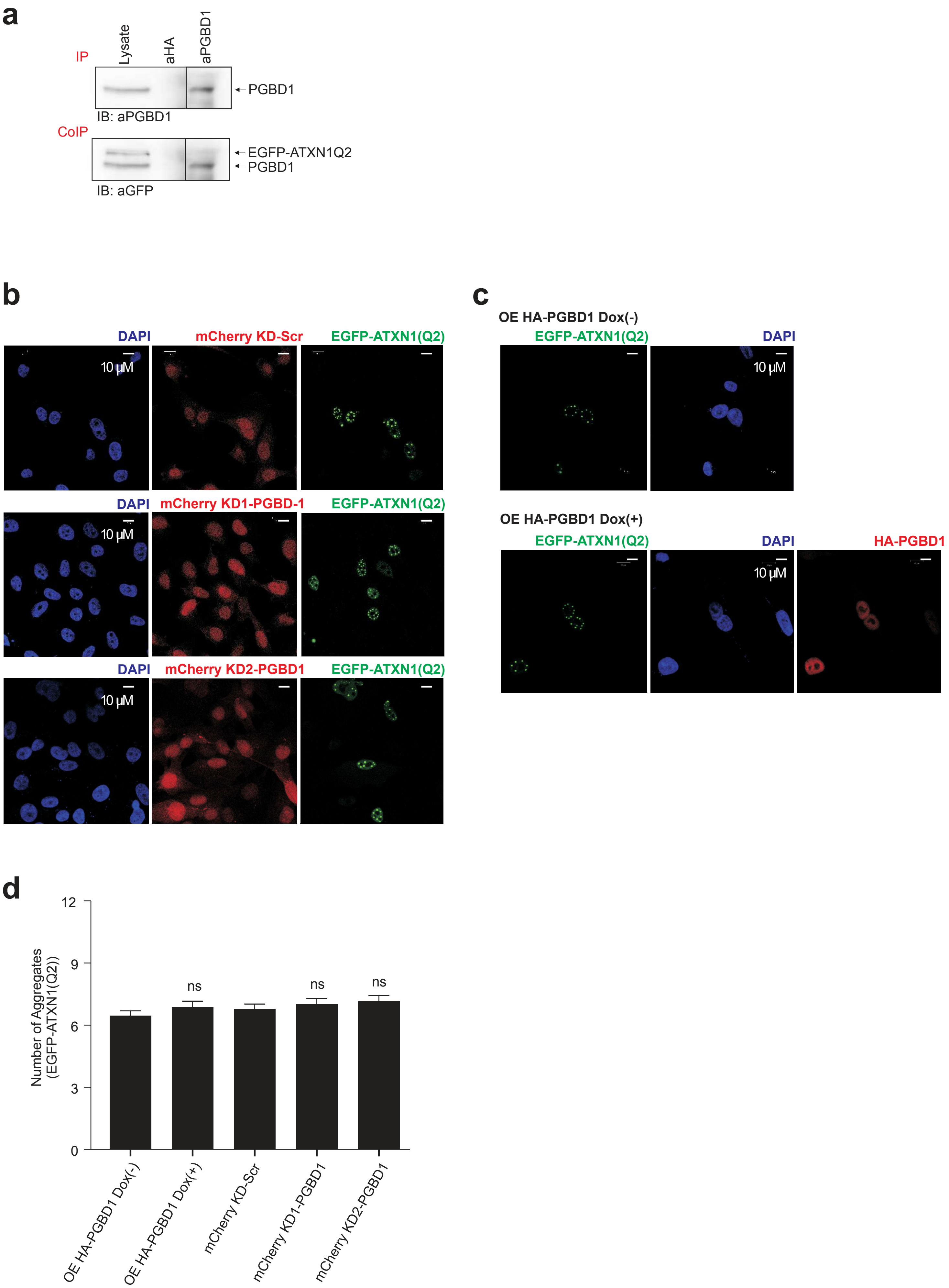

a
IP
Lysate
aHA
aPGBD1
PGBD1
IB: aPGBD1
CoIP
EGFP-ATXN1Q2
PGBD1
IB: aGFP
b
DAPI
mCherry KD-Scr
EGFP-ATXN1(Q2)
10 μM
DAPI
mCherry KD1-PGBD-1
EGFP-ATXN1(Q2)
10 μM
DAPI
mCherry KD2-PGBD1
EGFP-ATXN1(Q2)
10 μM
c
OE HA-PGBD1 Dox(-)
EGFP-ATXN1(Q2)
DAPI
10 μM
OE HA-PGBD1 Dox(+)
EGFP-ATXN1(Q2)
DAPI
HA-PGBD1
10 μM
d
Number of Aggregates (EGFP-ATXN1(Q2))
0
3
6
9
12
ns
ns
ns
OE HA-PGBD1 Dox(-)
OE HA-PGBD1 Dox(+)
mCherry KD-Scr
mCherry KD1-PGBD1
mCherry KD2-PGBD1

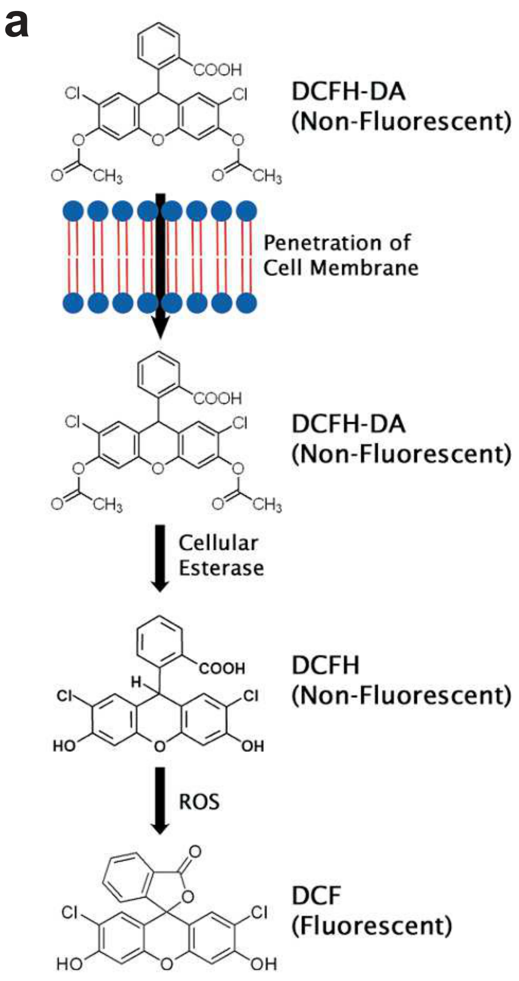


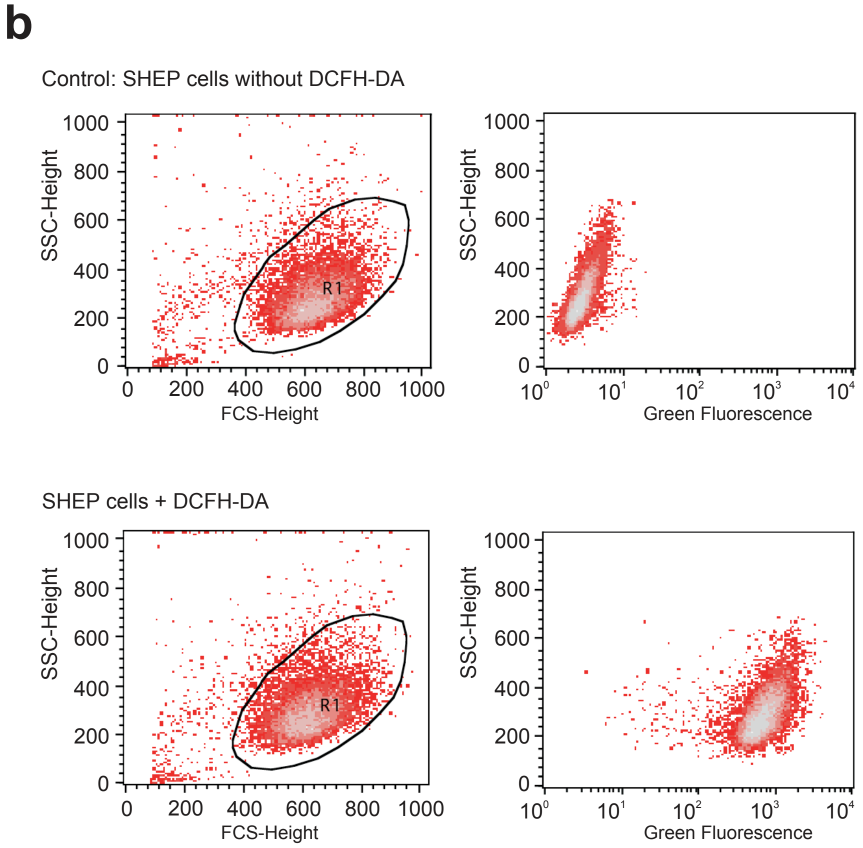


Based on the publication of Aranda et al., 2013.
Toxicol In Vitro. 2013 Mar;27(2):954-63. doi: 10.1016/j.tiv.2013.01.016
